# Nitrogen metabolism in *Pseudomonas putida*: functional analysis using random barcode transposon sequencing

**DOI:** 10.1101/2021.12.09.472016

**Authors:** Matthias Schmidt, Allison N. Pearson, Matthew R. Incha, Mitchell G. Thompson, Edward E. K. Baidoo, Ramu Kakumanu, Aindrila Mukhopadhyay, Patrick M. Shih, Adam M. Deutschbauer, Lars M. Blank, Jay D. Keasling

## Abstract

Pseudomonas putida KT2440 has long been studied for its diverse and robust metabolisms, yet many genes and proteins imparting these growth capacities remain uncharacterized. Using pooled mutant fitness assays, we identified genes and proteins involved in the assimilation of 52 different nitrogen containing compounds. To assay amino acid biosynthesis, 19 amino acid drop- out conditions were also tested. From these 71 conditions, significant fitness phenotypes were elicited in 672 different genes including 100 transcriptional regulators and 112 transport-related proteins. We divide these conditions into 6 classes, and propose assimilatory pathways for the compounds based on this wealth of genetic data. To complement these data, we characterize the substrate range of three promiscuous aminotransferases relevant to metabolic engineering efforts in vitro. Furthermore, we examine the specificity of five transcriptional regulators, explaining some fitness data results and exploring their potential to be developed into useful synthetic biology tools. In addition, we use manifold learning to create an interactive visualization tool for interpreting our BarSeq data, which will improve the accessibility and utility of this work to other researchers.

**IMPORTANCE:** Understanding the genetic basis of P. putida’s diverse metabolism is imperative for us to reach its full potential as a host for metabolic engineering. Many target molecules of the bioeconomy and their precursors contain nitrogen. This study provides functional evidence linking hundreds of genes to their roles in the metabolism of nitrogenous compounds, and provides an interactive tool for visualizing these data. We further characterize several aminotransferases, lactamases, and regulators--which are of particular interest for metabolic engineering.

## INTRODUCTION

As a free-living soil bacterium, *P. putida* encounters many different organic and inorganic nitrogen sources, and its responses to these conditions have been the target of recent study. During rhizosphere colonization, *P. putida* displays chemotaxis towards plant root exudates, which contain nitrogenous compounds such as benzylamines, polyamines, pyrrole derivatives, nucleotide derivatives, amino acids, and phenylpropanoids (1, 2). It also demonstrates chemotaxis towards and degradation of the phytotoxic and insecticidal benzoxazinoids exuded from the roots of maize seedlings (3, 4). Not only is *P. putida* able to withstand and metabolize these varied compounds in a nitrogen-rich rhizosphere, it is also capable of adapting to nitrogen-scarce conditions by increasing polyhydroxyalkanoate (PHA) production, repressing carbon catabolism, and increasing expression of transporters for nitrogen uptake (5).

Given its ability to adapt to the varied environment of the rhizosphere and its burgeoning role as a host for sustainable chemical bioproduction, it is not surprising that *P. putida*’s nitrogen metabolism has also been examined in the context of metabolic engineering. Due to the utility of PHAs as a next-generation bioplastic, the transcriptomic, proteomic, and metabolomic response of *P. putida* strains to nitrogen-limited growth conditions has been analyzed in order to better understand how PHA synthesis is triggered by nitrogen scarcity (6, 7). *P. putida*’s nitrogen metabolism and its regulation have also been a source of metabolic engineering parts. Some of its 39 predicted aminotransferases have been heterologously expressed as part of benzylamine derivative and glutaric acid production pathways (8, 9). The transcription factor regulating capro- and valerolactam degradation has been developed into a highly sensitive biosensor with the potential to be applied to increasing lactam production titers and used as an inducible system in pathway engineering (10). Moreover, many nitrogenous compounds are relevant building blocks for commodity chemicals, and understanding *P. putida*’s metabolism of them can enable more flux to be directed towards the desired product (11).

Despite its relevance to both metabolic engineering and basic scientific research, our understanding of *P. putida*’s nitrogen metabolism is far from complete. Many gene functions have been assigned through homology predictions with limited functional evidence, which can hamper the accuracy of metabolic modeling (12). Furthermore, *P. putida* has multiple paralogs of many of its enzymes, each potentially with different substrate preferences. Functional genomics can provide evidence for assigning gene functions and illuminate the specific roles of genes that have multiple paralogs (13).

In previous studies, we have employed barcoded transposon sequencing (BarSeq) to interrogate lysine, fatty acid, alcohol, and aromatic degradation in *P. putida* (13–15). In this work, we use BarSeq to study *P. putida* KT2440’s metabolism of 52 different nitrogen-containing compounds, nearly doubling the amount of publicly available BarSeq data for this bacterium. We provide evidence for many known nitrogen assimilation pathways and their regulatory systems, and also assign function to genes whose exact roles in nitrogen metabolism were not previously known. Due to their relevance in recent metabolic engineering efforts, we further examine the substrate specificity of *P. putida*’s 5-oxoprolinases and aminotransferases (8, 9, 16). To understand the regulation of these enzymes *in vivo*, we further characterized their cognate regulators. This work will bolster *P. putida*’s utility both as a host microorganism and as a source of metabolic engineering parts for sustainable chemical production.

## RESULTS AND DISCUSSION

### BarSeq reveals the genetic bases of diverse nitrogen metabolisms

Genes involved in nitrogen utilization from natural and unnatural compounds were identified using BarSeq. In these assays, a library of barcoded transposon insertion mutants was cultured in minimal media with glucose and a variety of sole nitrogen sources (Figure S1). A list of these nitrogen sources and the section in which each is discussed can be found in Table 1.

**Table 1:** Compounds used as nitrogen sources in BarSeq experiments and the section of the paper they are discussed in. Nitrogen sources indicated by (N) and amino acid dropout conditions indicated by (-).

| Section | Experiments included |  |
| --- | --- | --- |
| **Inorganic nitrogen sources and urea** | ammonium (N) | nitrite (N) |
|  | nitrate (N) | urea (N) |
| **Proteinogenic amino acids** | L-isoleucine (N) | glycine (N) |
|  | L-leucine (N) | L-histidine (N) |
|  | D-lysine (N) | L-alanine (-) |
|  | L-lysine (N) | L-arginine (-) |
|  | L-methionine (N) | L-asparagine (-) |
|  | L-ornithine (N) | L-aspartate (-) |
|  | L-pipecolic (N) | L-cysteine (-) |
|  | L-proline (N) | L-glutamate (-) |
|  | L-threonine (N) | L-glutamine (-) |
|  | L-phenylalanine (N) | glycine (-) |
|  | L-serine (N) | L-serine (-) |
|  | L-valine (N) | L-histidine (-) |
|  | 4-guanidinobutyrate (N) | L-isoleucine (-) |
|  | D-alanine (N) | L-leucine (-) |
|  | L-alanine (N) | L-lysine (-) |
|  | L-arginine (N) | L-methionine (-) |
|  | L-asparagine (N) | L-proline (-) |
|  | L-aspartate (N) | L-threonine (-) |
|  | L-cysteine (N) | L-tyrosine (-) |
|  | L-glutamate (N) | L-tryptophan (-) |
|  | L-glutamine (N) | L-valine (-) |
| **Quaternary amines + ethanolamine** | betaine (N) | choline (N) |
|  | carnitine (N) | ethanolamine (N) |
| **Purines and pyrimidines** | adenine (N) | hydantoin (N) |
|  | cytosine (N) | uracil (N) |
| **Lactams** | butyrolactam (N) | 5-oxoproline (N) |
|  | caprolactam (N) | valerolactam (N) |
| | γ-aminobutyric (N) | |
| **Polyamines, ω-amino acids, and GABA isomers** | 1,3 diaminopropane (N) | 6-aminocaproic acid (N) |
| | 1,6-diaminohexane (N) | β-alanine (N) |
|  | DL-2-aminobutyrate (N) | cadaverine (N) |
|  | DL-3-aminoisobutyrate (N) | putrescine (N) |
|  | 3-aminobutyrate (N) | spermidine (N) |
|  | 5-aminovalerate (N) |  |
| **Not applicable** | nicotinic acid (N) |  |

Wild-type growth on these substrates over 96 hours is shown in Figure S2. These assays revealed 672 genes with strong (|fitness| > 1) and significant (|t| > 5) fitness phenotypes. These include 100 transcription factors, 112 transport proteins, and numerous other enzymes with applications to engineered systems, such as aminotransferases (17, 18) (Figure 1A, Figure S3). Additionally, 529 of these genes encode proteins that are currently unreviewed in the Uniprot database, and 256 have not shown significant phenotypes in previous BarSeq studies. To visualize the fitness data, the manifold learning method, t-distributed stochastic neighbor embedding (t-SNE), was employed to cluster genes based on their fitness values in the tested conditions (Figure 1B, Figure S3) (19). The clusters were named based on the condition that elicited the largest and most frequent changes in fitness scores for genes within the cluster. In this visualization, we can identify genes that may take part in similar metabolisms. An interactive version that contains embedded hyperlinks to the Fitness Browser (20) can be accessed via this link: http://wintermute.ese.lbl.gov:8080/. Although genes with specific phenotypes to one condition are easy to identify, genes essential across many conditions are more challenging to assign a specific function. This is particularly problematic for genes that are essential in the majority of the tested conditions, like those involved in amino acid biosynthesis. To our surprise, clusters of genes involved in tryptophan, arginine, methionine, and branched chain amino acid biosynthesis were resolved by t-SNE (Figure I1-2). While not as exhaustive as pairwise comparisons of conditions, t-SNE provides a useful visualization of this diverse dataset.

**Figure 1:**
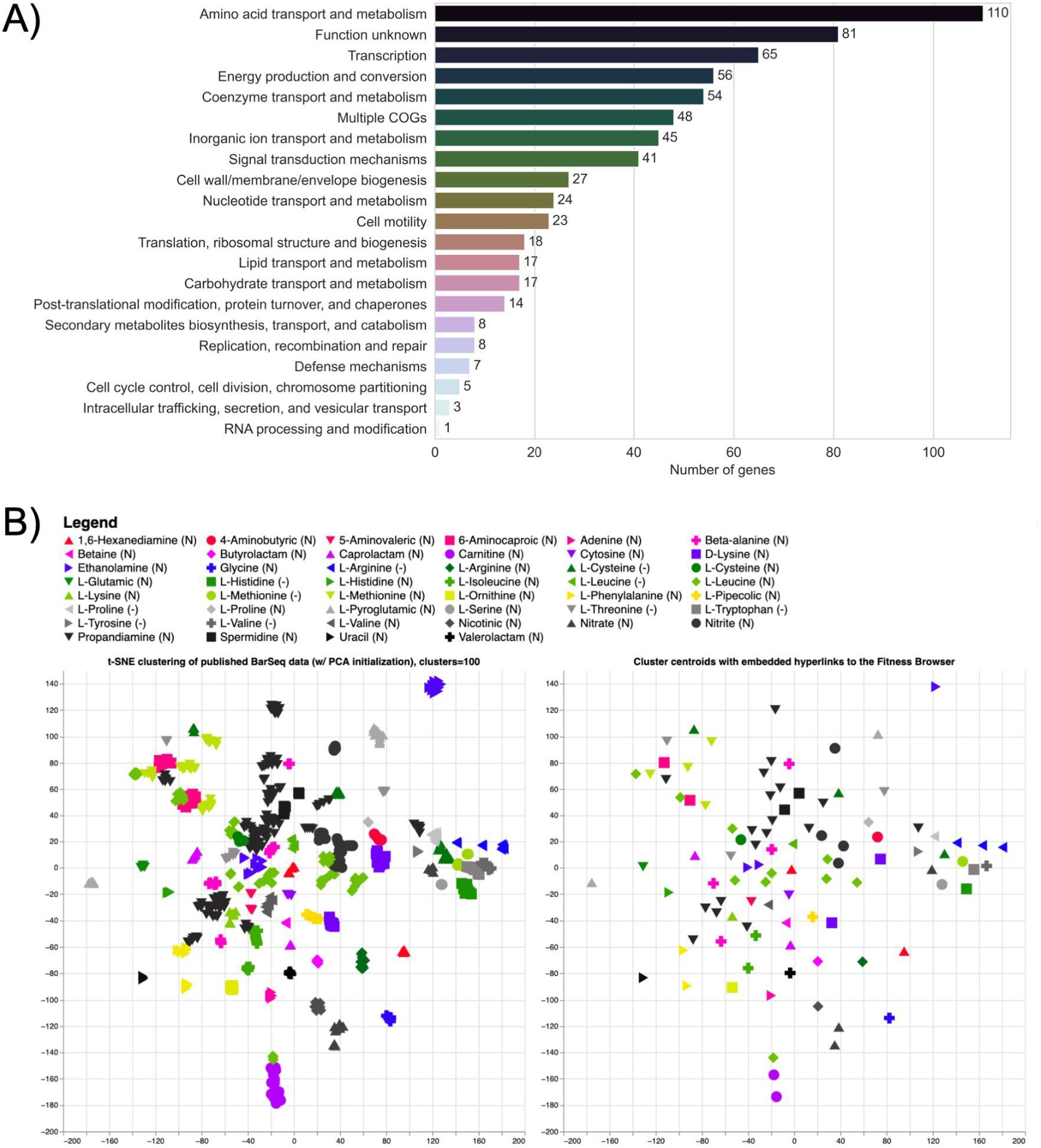
Global analysis of the *P. putida* KT2440 BarSeq data. A) Significant genes (|fitness| > 1 and |t| > 5) from all 71 tested conditions sorted by their Cluster of Orthologous Groups (COGs) based on the eggNOG database (21). B) Image of the interactive t-SNE visualization (<u>Figure I1</u>) showing the legend, t-SNE clustering (left) and cluster centroids (right). By clicking on a substrate in the legend, the corresponding cluster (left) and centroid (right) is highlighted, opening a list of cluster members and additional information. By clicking the highlighted centroid (right), the user is redirected to the Fitness Browser (https://fit.genomics.lbl.gov) (20), where the fitness data for all significant genes in the condition cluster is shown. An additional t-SNE visualization including COG identifiers is presented in <u>Figure I3</u>. More information about the interactive figures can be found in the description of Figure S3.

**Figure 2:**
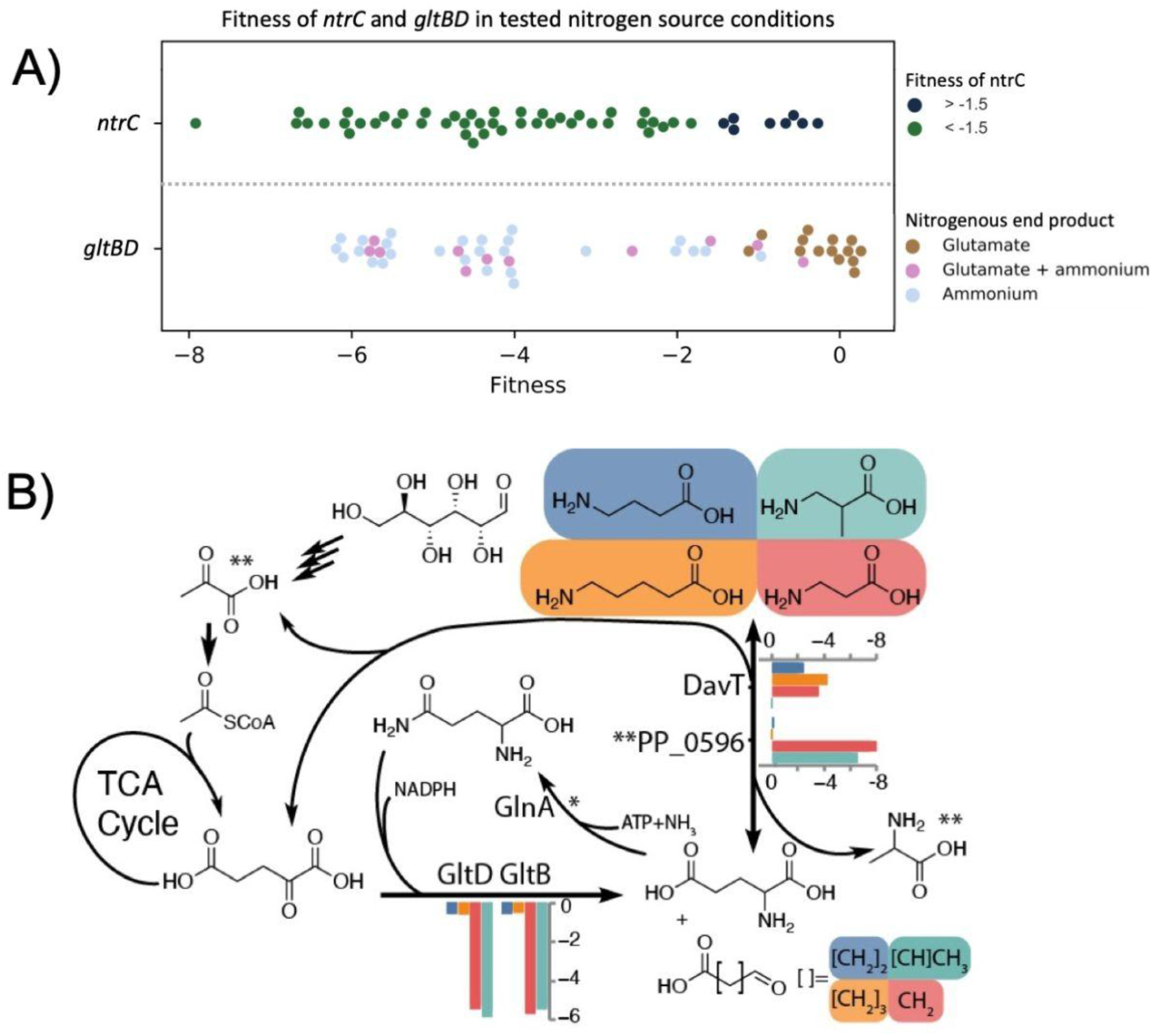
A) Scatter plot of the average fitness values (n=2) for *ntrC* and *gltBD* in all the tested nitrogen conditions. For *ntrC*, nitrogen conditions are grouped based on whether the fitness phenotype of *ntrC* is < -1.5 (blue) or > -1.5 (green). Conditions where ntrC fitness is >-1.5 (green) may be less dependent on *ntrC* activation and are shown in Table 2. *gltBD* phenotypes are sorted based on putative glutamate (brown), glutamate+ammonium (pink), or ammonium (grey) release during nitrogen source utilization. Fitness values for conditions resulting in *gltBD* fitness >-1.5 are shown in Table 3. B) The role of the GS/GOGAT cycle in the beta-alanine (red), 3-aminoisobutyrate (green), 4-aminobutyrate (blue) and 5-aminovalerate (orange) nitrogen source conditions. Average fitness values (n=2) are shown for *gltBD*, *davT,* and the pyruvate dependent transaminase PP_0596. *GlnA lacks fitness data because the library has no insertions. **Pyruvate and alanine are the specific nitrogen acceptor and product of PP_0596

**Table 2:** Nitrogen source conditions in which regulation may be less dependent on NtrC activation, indicated by *ntrC* fitness > -1.5 and illustrated by the dark blue points in Figure 2A.

| Conditions on which NtrC fitness is >-1.5 |  |
| --- | --- |
| Condition | NtrC fitness |
| L-glutamine | -0.27 |
| putrescine | -0.46 |
| L-asparagine | -0.57 |
| ammonium chloride | -0.66 |
| 4-aminobutyric acid | -0.86 |
| 4-guanidinobutyric acid | -1.30 |
| L-histidine | -1.31 |
| adenine | -1.43 |

**Table 3:**
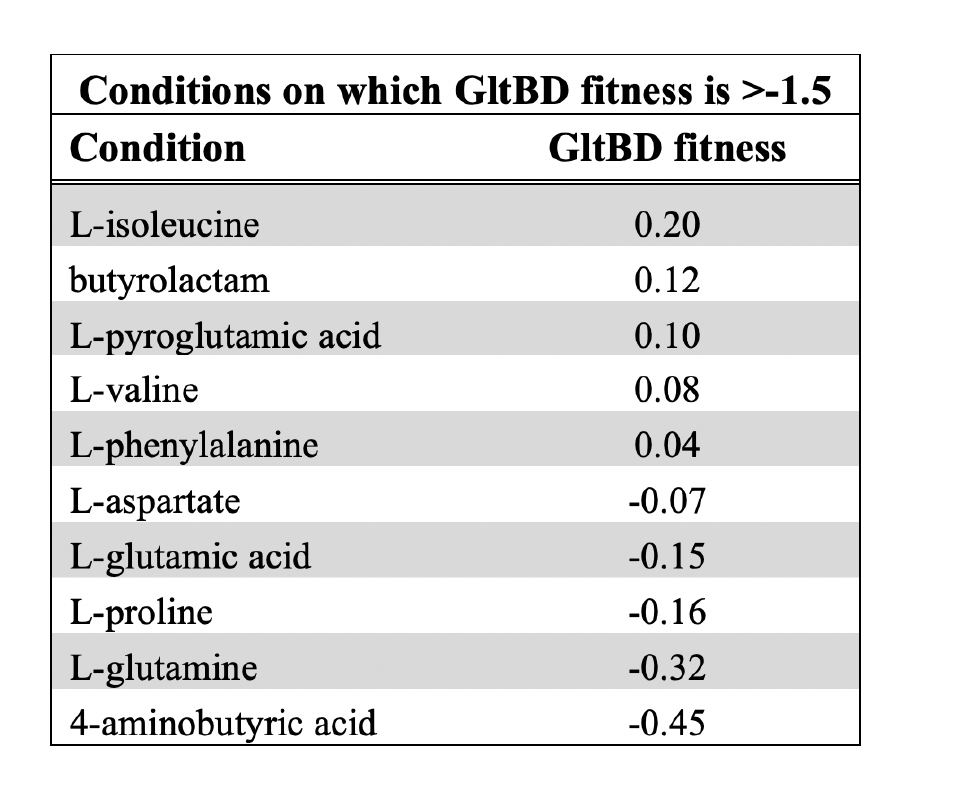
Nitrogen source conditions in which *gltBD* fitness > -1.5, illustrated by the scatter plot in Figure 2A.

### Global effectors of nitrogen metabolism

The regulators NtrB (PP_5047) and NtrC (PP_5048) have diverse phenotypes in the conditions tested. NtrBC is a two-component system that regulates the expression of numerous nitrogen assimilatory genes in *P. putida* (5). When the cell is nitrogen starved, the uridylyltransferase protein GlnD (PP_1589) modulates the activity of GlnK (PP_5234, protein PII). GlnK then activates NtrB, which affects the phosphorylation state of NtrC. Subsequently, NtrC modulates expression of its target regulon (5). While the library used for these assays has no insertions in *glnK*, the three other members of the signalling cascade are represented in the dataset and occupy the same cluster determined by t-SNE (Figure I1). It is unclear why certain nitrogen sources trigger a strong NtrC response (Figure 2A). The general role of NtrC is to counteract nitrogen starvation by activating the majority of nitrogen assimilatory genes, such as transporters (5). By setting an arbitrary fitness score cutoff at > -1.5, we identified at least 8 out of 52 nitrogen sources that might not rely on a functional copy of *ntrC* (Table 2). The utilization pathways for those compounds seem to be less dependent on NtrC activation, suggesting specific regulation systems or the presence of constitutively expressed transporter and degradation pathways for these nitrogen sources. The fitness profile of the extracytoplasmic function sigma factor SigX (PP_2088) also seems to be correlated with *ntrBC* based on cofitness and the clustering resolved from t-SNE. Previously, we found *sigX* to be partially essential for growth on D-lysine as a carbon source (15). SigX might be involved in regulating the catabolism of certain organic nitrogen sources. While no clear pattern in the fitness data could be determined for NtrC, another global factor involved in nitrogen metabolism, GltBD, illustrates a clear relationship between the conditions tested and their downstream metabolites.

GltB (PP_5076) and GltD (PP_5075) comprise the glutamate synthase (GOGAT) of *P. putida*, which plays an important role in the regulation of nitrogen assimilation. Interestingly, *gltBD* has diverse phenotypes across the conditions we tested (Figure 2). However, the second component of the central GS/GOGAT cycle, glutamine synthetase (*glnA*, PP_5046), has no insertions in our BarSeq library, indicating that it might have been essential during library construction. We were able to observe particularly strong fitness defects for *gltBD* in nitrogen conditions that either produce free ammonium or do not yield L-glutamate (Figure 2A) (22). For example, *gltBD* exhibits a very strong fitness defect in the nitrate (−6.17) and L-serine (−5.95) conditions but not in the L-phenylalanine condition (Figure 2A, Table 3). Another example is in the comparison of 5-AVA (5-aminovalerate), 4-ABA (4-aminobutyrate), β-alanine, and 3-AIBA (3-aminoisobutyrate). 3-AIBA utilization likely uses pyruvate as an amino-acceptor (via PP_0596), 4-ABA and 5-AVA require 2-oxoglutarate (via PP_0214/DavT), and β-alanine requires both aminotransferases (Figure 2B, Figure S4). An exemption to this rule is the nitrogen source ammonium chloride. The fitness defects for *gltBD* in this condition were much lower (|fitness| < 2), but still significant (|t| > 5). These data suggest that at high concentrations of ammonium (>10 mM), its assimilation is achieved by both the glutamate dehydrogenase (GdhA) PP_0675 and GltBD (5, 23, 24).

The regulators *gacS* (PP_1650) and *gacA* (PP_4099) show strong fitness phenotypes in our dataset. This two-component system is homologous to the well-studied *barA*/*uvrY* system of *E. coli*, and it has been identified as a global regulator of cellular physiology in diverse organisms (25, 26). The conditions that elicited the strongest negative fitness phenotypes for *gacSA* were L-alanine (−1.3), β-alanine (−1.8), spermidine (−2.0), and propanediamine (−7.4). The direct targets of GacSA regulation are the *rsm* noncoding RNAs which modulate the activity of translational repressors, resulting in global changes in gene expression (27–29). Through this mechanism, GacSA may be required for regulation of some portions of L-alanine, β-alanine, propanediamine, and spermidine metabolism.

### Inorganic Nitrogen Sources and Urea

*P. putida* KT2440 is a known obligate aerobe, and cannot use alternative terminal electron acceptors during oxidative phosphorylation (30). Because of this, oxidized nitrogen species can only be used as nitrogen sources. Although the preferred inorganic nitrogen source for many bacteria is ammonium (31), other inorganic nitrogen species such as nitrate and nitrite can be utilized via the assimilatory nitrate reduction system, often organized in a single gene cluster (32, 33). The enzymes associated with the initial steps of the assimilatory nitrogen system in *P. putida* are the reductases NarB and NirDB, for which we see significant growth phenotypes (Figure 3). In Pseudomonads, the two-component system NasS/T is a common regulator of this operon (33–35). In the absence of oxidized nitrogen sources, NasS and NasT form a complex that represses production of nitrate and nitrite reductases. When nitrate or nitrite are present, NasS dissociates from the NasS/T complex and the free RNA-binding antiterminator, NasT, enables full translation of the nitrate reduction operon (36) (Figure 3).

**Figure 3:**
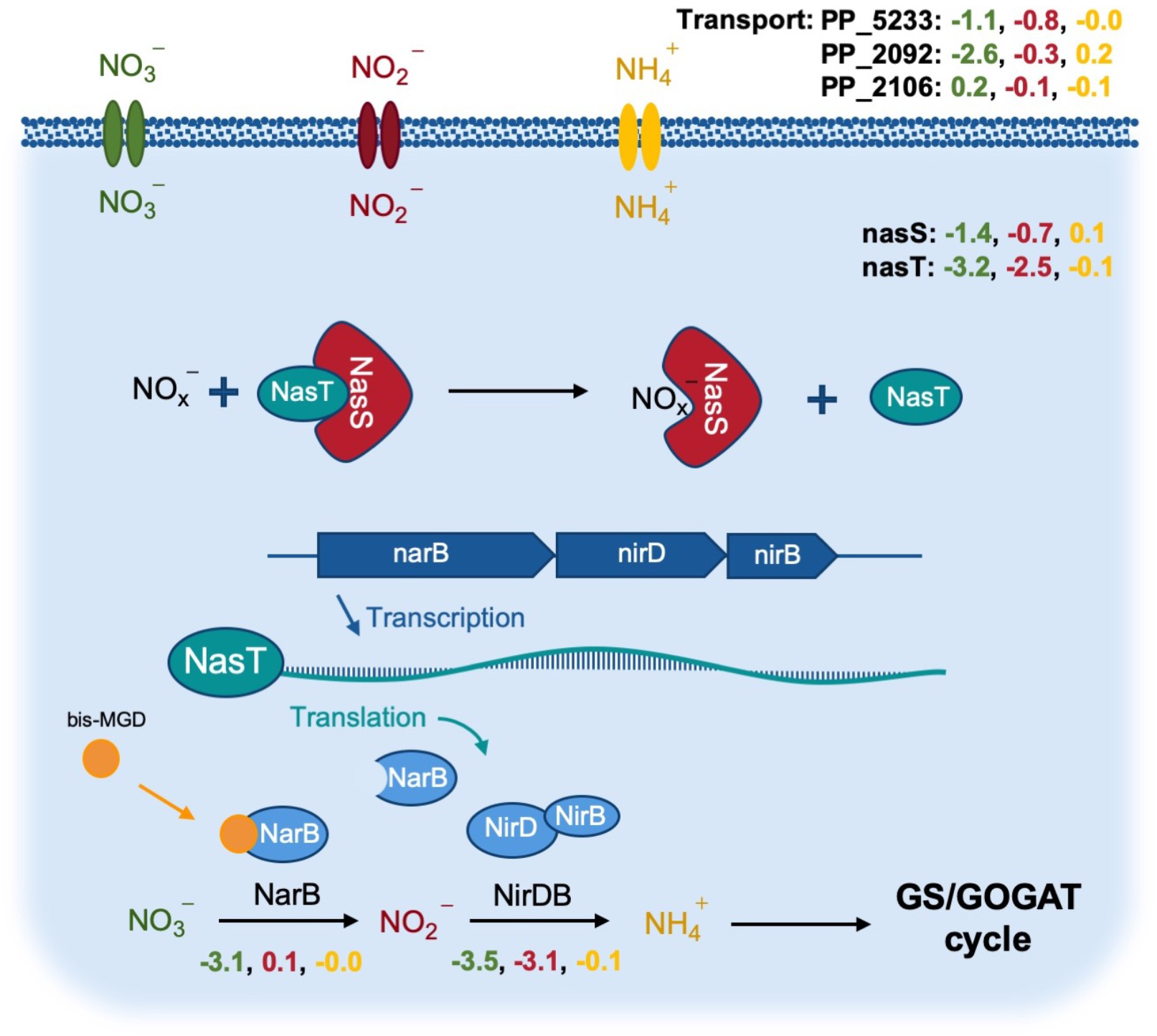
The assimilatory nitrate reduction system in *P. putida* KT2440. Average fitness values (n=2) exhibited in the nitrate (green), nitrite (red) and ammonium (yellow) sole nitrogen source experiments. Shown are putative transporters, action of the NasST regulatory system, the assimilatory pathway, and the role of the bis-molybdopterin guanine dinucleotide (bis-MGD) cofactor. bis-MGD is the required cofactor for NarB (48). The fitness phenotypes for bis-MGD biosynthesis cluster together with the nitrate phenotypes and can be found in the interactive t-SNE visualization (<u>Figure I1</u>).

Our competitive growth assay using BarSeq indicated that the same system may also operate in *P. putida* KT2440. In the nitrate condition, the specific growth phenotypes for *nasT* (PP_2093) and *nasS* (PP_2094) are represented by the significant fitness values of -3.2 and -1.4, respectively. We were also able to identify a distinct nitrate transporter PP_2092 (−2.6). However, no significant phenotype could be detected for this gene in the nitrite condition, suggesting that it may be specific to nitrate. Since there were no predicted transporters with strong fitness phenotypes in the nitrite condition, it is also possible that PP_2092 or other unidentified transporters are responsible for nitrite transport. At pHs less than 7.2 and concentrations greater than 100 µM, nitrite is also known to be passively transported into the cell via diffusion of nitrous acid, likely explaining the lack of fitness data for this transport reaction (37).

Overall, the nitrite condition demonstrates a pattern of fitness phenotypes that resembles a stress response. In its free acid form, nitrite has been shown to have antimicrobial properties and is hypothesized to have wide-ranging mechanisms of action, including DNA damage, collapse of the proton motive force, and deleterious nitrosylation of cofactors and proteins (38). Our fitness data indicate that disruption of the recently identified RES-Xre toxin-antitoxin (TA) module (PP_2433-4) is detrimental to growth in the nitrite condition. In this system, the toxin (PP_2434) rapidly degrades NAD^+^ to halt bacterial growth and give the organism time to adapt its survival strategies, while the antitoxin (PP_2433) inhibits the toxin to allow restoration of NAD^+^ levels (39, 40). The significant fitness defect of the antitoxin PP_2433 (−6.4) might indicate that either the RES-Xre system is part of *P. putida*’s stress response to nitrite, or that nitrite disturbs the redox homeostasis of the cell and further disruption of redox balance by the PP_2434 toxin is lethal in strains lacking a functional PP_2433 antitoxin. We also observed significant phenotypes for several pathways that oxidize NAD(P)H, which supports the idea that NAD^+^ is depleted in the nitrite condition. One example is *P. putida*’s altered glucose utilization strategy. Fitness data indicates that glucose oxidation to 2-ketogluconate is preferred over the gluconate phosphorylation or direct glucose uptake pathways (41–43). Indicators for the accumulation of 2-ketogluconate in the nitrate condition include the specific fitness phenotypes for the transporter *kguT* (PP_3377; -1.5) and the NAD(P)H-dependent dehydrogenase *kguD* (PP_3376; -2.65). Compared to the other two pathways, glucose utilization via 2-ketogluconate produces more NAD(P)^+^. Other conditions that seemed to lead to similar phenotypes for *kguT* were 1,6-hexanediamine (−1.3), caprolactam (−1.55), 2-aminobutyric acid (−1.5) and uracil (−1.2).

Urea, the simplest organic nitrogen source we tested, only elicited significant fitness phenotypes in *gltBD* (−5.45) and *ntrC* (−1.8). Transcriptome analysis has previously revealed that the expression of a putative urease operon in *P. putida* (PP_2842-9) is controlled by NtrC (5).

However, we observed no significant phenotypes in this operon. The presence of a second urea degradation pathway via urea carboxylase is unlikely, because it has been shown that organisms that possess the carboxylase pathway typically lack an urease (44–47). More research is necessary to further characterize urea metabolism in *P. putida*.

### Proteinogenic amino acids

The amino acid metabolism of microorganisms is of special interest for the food and bulk chemical industries, and has therefore been studied extensively for many decades (49). In contrast to the genus *Corynebacterium*, *Pseudomonas spp.* play a minor role in the industrial production of amino acids (50). However, their exceptional tolerance towards organic solvents makes them an excellent host for production of aromatic amino acid-derived compounds (11, 51–60).

Here, we have tested the 20 proteinogenic amino acids and two D-stereoisomers as the sole nitrogen source in a competitive growth assay using BarSeq. We also included BarSeq data from experiments where L-arginine, L-histidine, L-lysine, and D-lysine were used as sole carbon sources, and ammonium chloride was provided as a nitrogen source. No fitness data could be obtained for tyrosine and tryptophan due to insufficient biomass for DNA extraction and BarSeq analysis. Furthermore, although is has been reported that *P. putida* is unable to grow with the L-stereoisomers of methionine, threonine, or leucine as the sole source of carbon and nitrogen, we observed that *P. putida* could grow with these as sole nitrogen sources (Figure S2) (61). The flux of their carbon backbones into the TCA cycle might be too inefficient for these three amino acids to simultaneously be the sole source of both carbon and nitrogen.

Microorganisms may possess multiple pathways for the degradation of the same amino acid, especially in the case of L-arginine, for which there are five known degradation pathways in *P. aeruginosa* (62–64). Our nitrogen BarSeq experiments confirmed the presence of two of these pathways, and carbon source BarSeq data confirmed the presence of a third. Although another confirmed pathway in *P. aeruginosa*, the deiminase pathway, could not be identified in *P. putida* using BarSeq, the *P. putida* arginine deiminase AraA (PP_1001) has been successfully characterized *in vitro* (65).

The use of L-arginine as the sole nitrogen source led to significant growth phenotypes in both the common arginine succinyltransferase (AST) pathway (< -2.7) and the arginine decarboxylase/agmatine deiminase pathway. Ornithine can also be incorporated into the AST pathway, however, the only significant fitness defect we observed in this condition was for the dehydrogenase AstD (PP_4478; -2.3) (66, 67). Interestingly, the required succinyltransferase (PP_4479-80) to channel ornithine into the AST pathway exhibits a positive fitness value (1.5). Under these conditions, ornithine degradation through the AST pathway may not be optimal. Instead, direct deamination by the cyclodeaminase PP_3533 (−0.85) and conversion of proline to glutamate (PP_4947; -5.15) might be more favorable.

The presence of a functional arginine decarboxylase/agmatine deiminase (ADAD) pathway in *P. putida* is indicated by the fitness defect of the arginine decarboxylase PP_0567 (− 0.75). The amidase family protein PP_2932 has a strong fitness defect (−1.85) in the arginine condition, suggesting that it acts on N-carbomylputrescine as part of the ADAD pathway and should be more specifically annotated as an N-carbomylputrescine amidase.

When L-arginine is used as the sole carbon source, significant phenotypes for the arginine:pyruvate transaminase (APT) pathway appear as well. This pathway consists of the transaminase AruH (PP_3721, -1.2), the decarboxylase AruL (PP_3723, -0.9), the dehydrogenase KauB (PP_5278, -1.6) and a guanidinobutyrase (PP_4523, -1.5). The putative guanidinobutyrase PP_4523 not only has a fitness defect with L-arginine as the carbon source, but also with 4-guanidinobutyric acid as the nitrogen source (−0.95).

Due to a slight fitness defect (−1.0) of the predicted alanine racemase (Alr) PP_3722 and its localization in the APT operon, it is unclear whether AruH acts on the D- or L-stereoisomer of arginine, or on both. In a previous study, the deletion of *alr* led to significantly decreased growth of *P. putida* with arginine as the sole carbon and nitrogen source (68). This might indicate the presence of another catabolic route for arginine via its D-stereoisomer.

Carbon and nitrogen availability could determine the preference for one pathway over another. Since the AST pathway requires succinate for the conversion of arginine to glutamate, it is less efficient under carbon-limiting conditions, whereas the products of the other pathways can feed directly into the TCA cycle via putrescine and GABA degradation (62).

The predicted histidine-lysine-arginine-ornithine ABC transport system (PP_4483- PP_4486) shows significant phenotypes (> -2) only in the arginine condition, while the ornithine condition results in mild fitness defects (−0.44). Furthermore, a different transporter, PP_5031, seems to be the most important transporter for histidine (−0.55). Interestingly, PP_5031 has a much stronger phenotype in the histidine carbon source condition (−4.5). Since NtrC responds to nitrogen-scarce conditions by upregulating nitrogen transporters (5), it may be challenging to identify transporters that are distinctly expressed in the presence of a specific substrate. These mild fitness defects for transporters in the histidine and ornithine nitrogen source conditions suggest that multiple transporters capable of accepting histidine or ornithine are expressed as part of the NtrC response. This limitation of nitrogen source BarSeq assays might be circumvented by adding ammonium in excess or complementing nitrogen source BarSeq data with carbon source experiments.

Another interesting interaction between D- and L-stereoisomers can be observed in the catabolism of lysine and alanine. Lysine catabolism in *P. putida* has been extensively studied and was fully characterized by Thompson et al., who used D- and L- lysine as a carbon source for BarSeq experiments (15, 61, 68–73). By comparing our nitrogen source data with these previous BarSeq datasets, we can determine the differences between D- and L-stereoisomer catabolism and utilization as carbon or nitrogen sources (Figure S5).

When L-lysine is fed as a carbon source, we see similar significant fitness phenotypes in genes of the pathways for both enantiomers (Figure S6). Likewise, when D-lysine is provided as a carbon source, both pathways have fitness defects, but the L-lysine pathway does appear less essential (15). The importance of both pathways for both enantiomers is most likely due to an lysine-inducible or constitutive expression of the *P. putida* alanine racemase *alr*, which can then convert the provided isomer of lysine to its stereoisomeric counterpart (61, 74). The same behavior can be observed when both lysine enantiomers are used as the sole nitrogen source.

The key difference between the lysine carbon and nitrogen source datasets is in the genes PP_4493, PP_0213 and *gltBD*. PP_0213 and PP_4493 are involved in the flux of the carbon skeleton of D- and L-lysine into the TCA cycle and are therefore critical under carbon-limiting conditions. Since the substrates of both gene products are fully deaminated, their fitness values in nitrogen-limiting conditions are negligible. The strong phenotype for *gltBD* (−5.8) is likely caused by the release of free ammonium in the first deamination step.

Along with D-glutamate, D-alanine is an important compound for the synthesis of microbial peptidoglycan and therefore necessary for bacterial survival and growth (75). The main racemase in *P. putida* for alanine appears to be DadX (61, 76). Our BarSeq data for alanine suggest that the conversion reaction was essential for *P. putida*’s survival when the library was constructed, since there are no *dadX* (PP_5269) mutants present in our mutant library. The *dadX* paralog *alr* may be able to convert alanine as well, but its insignificant growth phenotype implies that it only plays a minor role in alanine degradation (61, 68, 76). Furthermore, neither of the alanine transaminase homologs show a growth defect when either L- or D-alanine is used as the sole nitrogen source. The only significant growth defect under these conditions is that of the deaminating D-amino acid oxidoreductase PP_5270 (*dadA*) and interestingly, its fitness defect is only significant with L-alanine (−3.6). For D-alanine, it only has a slight fitness defect of -0.65.

An indicator that D-alanine deamination is still dependent on PP_5270 is the significant growth defect caused by the transcriptional regulator PP_5271 (−2.05). The gene PP_5271 has high identity (93% with an E-value of < 9e-88) with the regulator *lrp* from *P. aeruginosa* PA14. In *P. aeruginosa* PA14, *lrp* is a transcriptional activator of the *dad* operon, which is required for L-alanine catabolism. The regulator is highly induced by L-alanine and also slightly less induced by D-alanine and L-valine (77). The strong negative fitness value of -2.05 also supports the theory that *lrp* is an activator rather than a repressor in *P. putida* (78).

Pseudomonads are well known for their ability to degrade and synthesize aromatic compounds (79). Therefore, it is not surprising that they have also developed a large repertoire of aminotransferases with activity on aromatic substrates (80, 81). The only aminotransferase we identified in an aromatic amino acid condition (L-phenylalanine) is PP_3590 (−4.0). Its gene product is annotated as the D-lysine aminotransferase AmaC, although it plays a minor role in D-lysine degradation (73). Our BarSeq experiments suggest that PP_3590 has a broad substrate range, with fitness defects appearing on other substrates as well. For example, it has a fitness defect of -1.25 on L-pipecolate, indicating potential L-2-aminoadipate transaminase activity. The homolog, PP_1972, seems to have little effect on phenylalanine (−0.35) or pipecolate (−0.15) growth. In a previous study, a double knock-out mutant of PP_3590 and PP_1972 did not lead to phenylalanine auxotrophy, underlining the redundancy of aromatic aminotransferases in *P. putida* (81).

Overlaps between aromatic amino acid transferases and branched-chain amino acid (BCAA) metabolism are also common in bacteria (82). Our data indicate that the aromatic aminotransferase PP_3590 also seems to be involved in the transamination of L-isoleucine (− 1.75). However, the main transaminase in BCAA degradation and biosynthesis is PP_3511 (*ilvE*). It shows a significant fitness defect (< -4) in almost all the tested conditions, including ammonium chloride (−5.55). This highlights the importance of PP_3511 in minimal media conditions and its unique role in the metabolism of BCAAs. In fact, IlvE-deficient strains require gene complementation or supplementation of all three BCAAs (valine, isoleucine, leucine) to restore growth in minimal media (83, 84). We observe a similar behavior for the histidinol-phosphate aminotransferase PP_0967, which is essential for all the tested conditions except histidine. Surprisingly, heterologous expression of PP_0967 has been used to facilitate the deamination of L-phenylalanine to phenylpyruvate (56).

Many steps of the amino acid catabolism in *P. putida* are already known or precisely predicted based on homology models. However, by using BarSeq, we were able to generate functional evidence for the pathways of at least eight additional amino acids beyond the ones discussed here. To further expand our understanding of the amino acid metabolism in *P. putida*, we also included drop-out growth experiments for all proteinogenic amino acids. By supplying all but one of the 20 proteinogenic amino acids, we created conditions where biosynthesis of amino acids is essential for growth. We refer to the Fitness Browser (https://fit.genomics.lbl.gov) and the interactive t-SNE visualization (Figure I1-2) for more details.

### Quaternary amines and ethanolamine

Choline is a trimethylated, positively-charged amine and a common component of ionic liquids used in the depolymerization of lignin (85). In nature, choline is metabolized by most bacteria to serve as a precursor for betaine, which is an important osmoprotectant (86). While biosynthetic pathways for those compounds are rarely present in bacteria, transporters and catabolic routes for quaternary amines are ubiquitous in these organisms (87, 88).

It is predicted that choline, betaine (trimethylglycine), and carnitine are metabolized through convergent pathways, with carnitine and choline entering betaine metabolism following thiolase cleavage and alcohol oxidation, respectively (89) (Figure 4A). In the subsequent reactions, betaine is demethylated to produce glycine. Even though the glycine cleavage system (PP_0986, PP_0988 and PP_0989) has a significant phenotype when glycine is used as the sole source of nitrogen (< -4), it is not essential for growth in the quaternary amine conditions (> - 0.3). Instead, conversion into serine is more efficient due to the generation of the required methyl-group donor 5,10-methylenetetrahydrofolate during demethylase activity of PP_0310-1 (90). As indicated by the strong fitness phenotypes elicited in the carnitine, choline, and betaine conditions, this reaction is likely catalyzed by PP_0322 (a predicted glycine/serine hydroxymethyltransferase) (< -2). Although PP_3144 (L-serine dehydratase) appears essential for growth on L-serine as a sole nitrogen source (−2.3), it is non-essential for growth on the tested quaternary amines. This suggests that other serine dehydratases (PP_0297 or PP_0987) or deaminating enzymes may be expressed in these conditions to release ammonium from serine.

**Figure 4:**
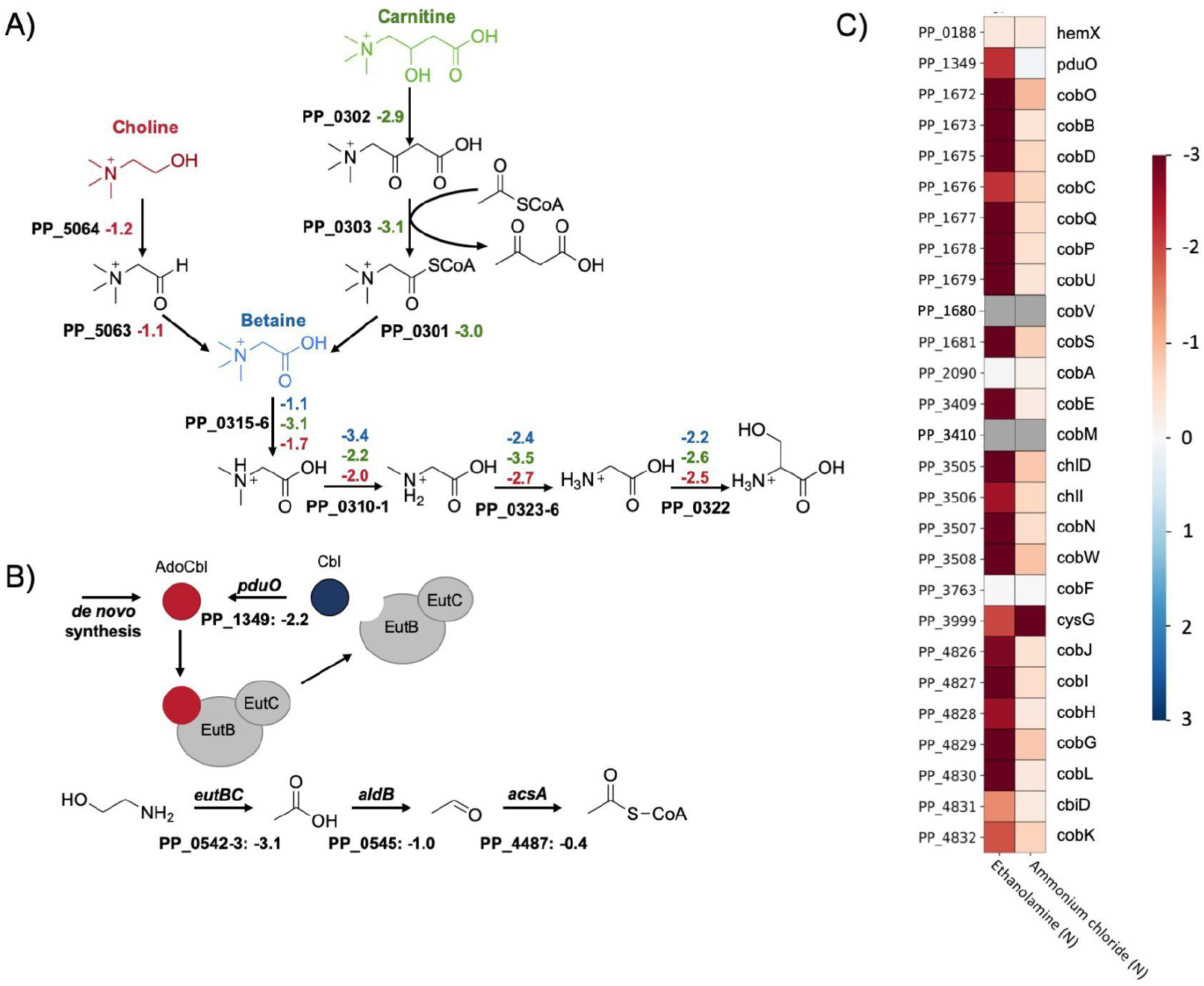
Quaternary amine and ethanolamine degradation in *P. putida*. A) Putative routes for the quaternary amine catabolism in *P. putida* KT2440. The figure shows the degradation of choline (red), carnitine (green) and betaine (blue). The corresponding average fitness scores (n=2) are shown next to each gene. B) Ethanolamine degradation pathway, shown with fitness values (n=2) and regeneration of the AdoCbl cofactor. C) Heatmap with average fitness scores (n=2) of genes that are putatively involved in *P. putida*’s adenosylcobalamin biosynthesis. No fitness scores (grey) could be obtained for the genes PP_1680 (cobV) and PP_3410 (cobM).

Based on the results of the BarSeq assay and sequence homology, the initial steps in carnitine metabolism were also identified. We propose that the beta-oxidation of carnitine to betainyl-CoA proceeds through the genes PP_0301 (a putative thioesterase), PP_0303 (a dehydrocarnitine cleavage enzyme), and PP_0302 (L-carnitine dehydratase) (Figure 4A). The carnitine metabolism of *P. aeruginosa* was originally believed to proceed via CoA activation of oxidized carnitine (91). Enzymatic characterization of a PP_0303 homolog has since suggested that this protein acts upon oxidized carnitine and acetyl-CoA, releasing acetoacetate and betainyl-CoA (92).

We also observed fitness phenotypes for the transporters and regulators involved in quaternary amine metabolism. Although all three quaternary amines cause a strong NtrC response, which tends to mask distinct transporters, PP_0294-6 (choline/betaine/carnitine ABC transporter) exhibits fitness defects for choline (−1.13) and carnitine (−2.55). The metabolism of betaine and its precursors is regulated by the repressor BetI (PP_5719) and the activators GbdR (PP_0298) and CdhR (PP_0305) (87, 93). While *gbdR* is essential in all three conditions (< -1.0), *cdhR* is specific to carnitine (−2.5) and *betI* is specific to choline (1.15). The negative and positive fitness defects correlate with the predicted mode of action for these regulators (93). The reason for the significant fitness phenotypes of PP_0308-9 in these conditions and their role in this operon are still unknown. Transcriptomic analysis suggested that they might be involved in the formation of filamentous biofilms (94).

Ethanolamine is a common molecule in nature and is involved in the choline and serine metabolism of plants (95, 96). Therefore, it is not surprising that several Pseudomonads are able to use it as a source for carbon and nitrogen (97). Bacterial ethanolamine degradation can be divided into two routes: (i) via acetyl-CoA or (ii) via ethanol (97, 98). However, the first step in both pathways is catalyzed by the adenosylcobalamin-dependent ethanolamine ammonia-lyase (EAL) EutBC (98–100). In *P. putida*, we observed a strong fitness phenotype for *eutBC* (PP_0542-3; -3.1) in the ethanolamine condition (Figure 4B). Furthermore, we were also able to support the requirement for its cofactor adenosylcobalamin (AdoCbl). Since there is no exogenous AdoCbl in our minimal media and no specific phenotypes for the corrinoid-specific transport system PP_0524-5, *P. putida* is likely capable of the *de novo* synthesis of AdoCbl (101–104). This is also supported by the strong fitness phenotypes we observed for the putative *cob* genes in this condition (Figure 4C). The additional requirement of the adenosyltransferase PduO (PP_1349; -2.2) is most likely caused by the release of Cbl during EAL activity (105–108). Re-adenylation of Cbl by PduO is likely less metabolically demanding than the *de novo* synthesis of AdoCbl.

### Purines and pyrimidines

Purines and pyrimidines are widely distributed chemical structures in nature, and many organisms also salvage them as nitrogen sources. In this study, we have tested the purine base adenine and the pyrimidine bases cytosine and uracil as sole nitrogen sources using BarSeq.

Similar to many other purines, the first steps in the degradation of adenine and guanine are the conversion into the shared intermediate xanthine (Figure 5A). While adenine is converted to xanthine via the intermediate hypoxanthine, guanine is directly converted to xanthine by the guanine deaminase PP_4281 (109). The slight fitness defect of the adenosine deaminase PP_0591 (−0.45) might indicate that it can also act on adenine as a substrate.

**Figure 5:**
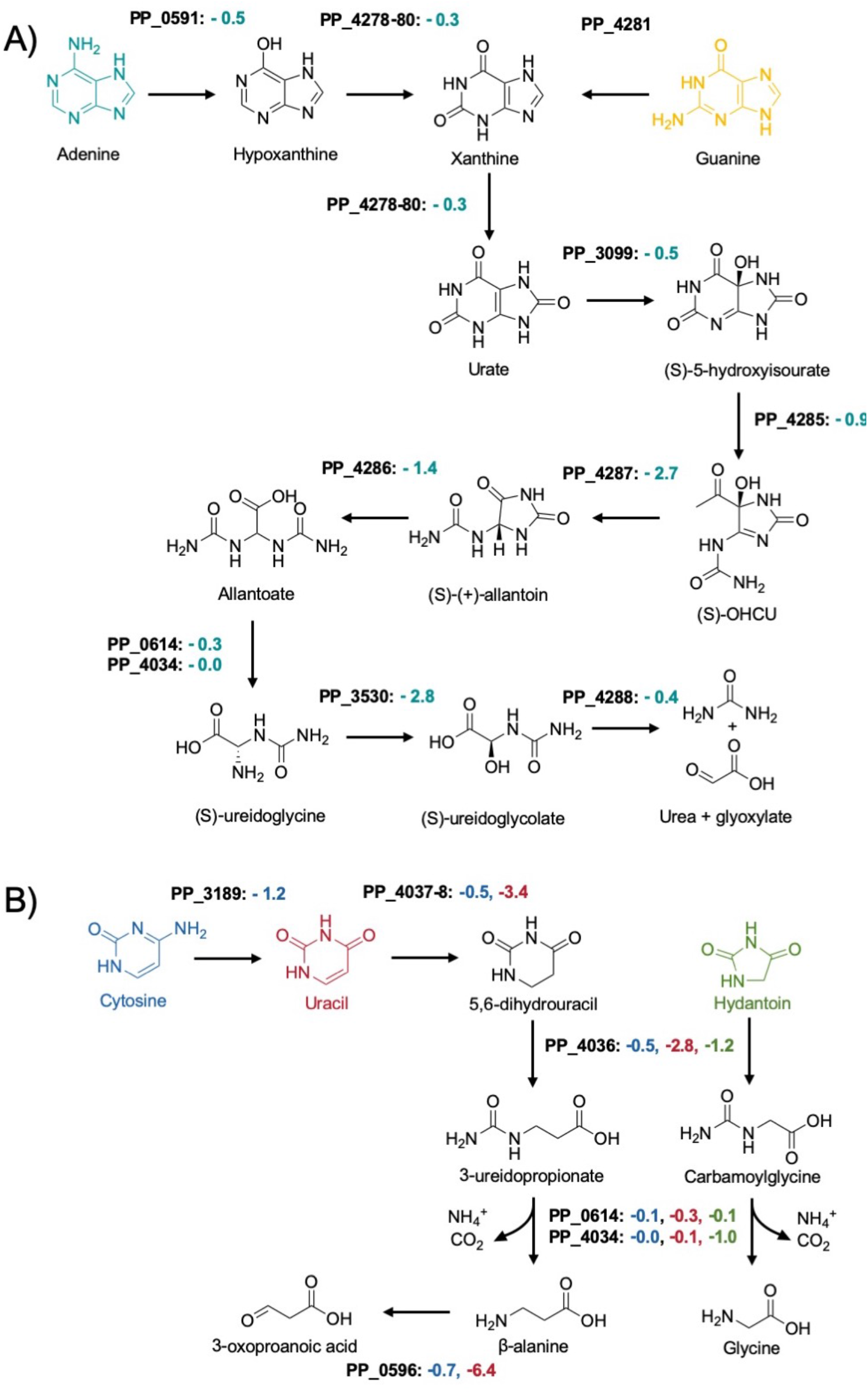
Putative routes for purine (A) and pyrimidine (B) catabolism in *P. putida* KT2440. Shown are the average fitness scores (n=2) for genes involved in adenine (teal), guanine (yellow), cytosine (blue), uracil (red) and hydantoin (green) degradation.

The deamination products of the purine nucleobases, hypoxanthine and xanthine, are then both converted by the xanthine dehydrogenase complex *xdhABC* (PP_4178-80) to yield urate. For many years it was believed that the urate oxidation reaction yields (S)-allantoin in a single step (110). However, it was recently discovered that in addition to urate oxidase, this reaction actually involves two other enzymes and goes through the intermediate 5-hydroxyisourate, releasing hydrogen peroxide (111). To date, no urate oxidase has been experimentally verified in *P. putida*, and the only potential urate oxidase (PP_3099) exhibits no growth phenotype in any of the tested conditions. An indirect indicator for this reaction could be the significant fitness defect of the LysR-type transcriptional regulator PP_2250 (−2.35) and the conserved membrane protein of unknown function PP_2251 (−2.25). PP_2251 has high identity with the proteobacterial antimicrobial compound efflux (PACE) transporters A1S_1053 (57% with an E-value of 6e-45) from *Acinetobacter baumannii* and PFL_4585 (64% with an E-value of 5e-56) from *P. protegens Pf-5*. It is possible that PP_2251 might also be a PACE transporter that functions as part of a defense mechanism against hydrogen peroxide or spontaneous peroxide radicals formed during urate oxidation (112, 113). The downstream reactions of 5-hydroxyisourate conversion to (S)- allantoin were strongly indicated by BarSeq. These reactions and further processing of (S)- allantoin to glyoxylate and urea are shown in Figure 5A.

The degradation of the pyrimidine base cytosine begins with the removal of the amine group by the cytosine deaminase CodB (PP_3189; -1.2) (Figure 5B). Furthermore, we were able to confirm the significance of the designated cytosine transporter CodA (PP_3187; -1.2).

Unexpectedly, the same transporter and deaminase cause fitness defects in the 3-aminobutyric acid condition (−1.15 and -1.15, respectively). The expression of the *codBA* operon and many other transport systems for nitrogenous compounds is controlled by the global regulator NtrC (5, 114). A general survival strategy for *P. putida* under nitrogen-limiting conditions seems to be the expression of numerous transporters for nitrogen-containing compounds that could potentially be present in the medium (5). Since 3ABA is an uncommon metabolite in nature, it might activate NtrC to such an extent that the cell starts scavenging nucleotides as a first response to nitrogen starvation (5, 114). Fitness defects for the hydantoinase PP_4036 (−1.8) and the dihydropyrimidine dehydrogenase PP_4037-8 (−2.78) are additional indicators for the degradation of nucleotides.

Since the deamination product of cytosine is uracil (Figure 5A), similarly strong fitness values in both conditions could be expected. However, the fitness values associated with uracil degradation were at least 5-fold smaller in the cytosine condition. Our data suggest that in some cases subsequent deaminations of the same compound lead to less significant phenotypes; other examples include lysine and arginine nitrogen source experiments (Figure S6).

The ring-opening reaction for 5,6-dihydrouracil is catalyzed by PydB (PP_4036; -2.8), and the resulting amide (S)-ureidoglycolate is then hydrolyzed by *hyuC* (PP_4034) or its predicted paralog PP_0614. Both (S)-ureidoglycolate hydrolysing genes have no significant fitness defect, likely due to functional redundancy. Surprisingly, the only significant growth phenotype we identified for *hyuC* was in the hydantoin condition.

Hydantoin was discovered as a reduction product of allantoin (115), and its derivatives are used as anticonvulsants and pesticides (116, 117). Because it is a xenobiotic compound, the ability of *P. putida* to utilize hydantoin is probably due to its structural similarity to 5,6- dihydrouracil. The pyrimidine permease PP_4035 (−1.05) appears to be involved in hydantoin transport, and the same dihydropyrimidinase (PydB) that acts on 5,6-hydrouracil is likely responsible for opening its 5-membered ring (−1.15), producing carbamoylglycine. The amidohydrolase *hyuC* has a fitness defect of -1.0 in hydantoin sole nitrogen source experiments, which is the strongest fitness defect we observed for this gene. This could be explained if hydantoin was not within the substrate range of its paralog PP_0614 (−0.15). It is known that R- substituted hydantoins are converted by bacteria to the corresponding D-amino acid (118). Although comparing the hydantoin condition with the glycine condition did not reveal any common phenotypes involved in these metabolisms, it is likely that the products of the carbamoylglycine hydrolysis are glycine, ammonia, and carbon dioxide.

Even though we have no data for thymine, we tested a racemic mixture of 3- aminoisobutyric acid (3-AIBA) as the sole source of nitrogen. Its D-stereoisomer is the final product of thymine degradation (119). To date, no aminotransferases have been identified in *P. putida* that show either D- or L-3-AIBA activity. However, the strong fitness phenotypes for the pyruvate transaminase PP_0596 (−6.3) and the dehydrogenase PP_0597 (−1.7) are most likely caused by the presence of the L-stereoisomer, which is a common intermediate in valine degradation (120). Another potential indicator that PP_0596 has 3-AIBA transaminase activity is the fitness defect for the transcriptional regulator *lrp* (−2.15) and the deaminating D-amino acid oxidoreductase PP_5270 (−1.4). As described earlier, these genes are part of the *dad* operon which is involved in alanine metabolism and therefore an indicator for pyruvate transaminase activity (77).

### Lactams

With applications ranging from manufacturing solvents to precursors for plastics and pharmaceuticals, lactams are an industrially relevant class of chemicals (121). Here, we tested four different lactams as nitrogen sources in our BarSeq experiments, gaining functional evidence for two previously uncharacterized lactam hydrolases, PP_2920-2 and PP_4575-7. Fitness data indicate that these lactam hydrolases are responsible for the hydrolysis of butyrolactam and 5-oxoproline (Figure 6A). In-frame deletions of each of these hydrolases abolished growth when their respective lactam was provided as the sole nitrogen source. Growth could then be restored by complementation with a pBADT-based plasmid containing the lactam hydrolase (Figure S7).

**Figure 6:**
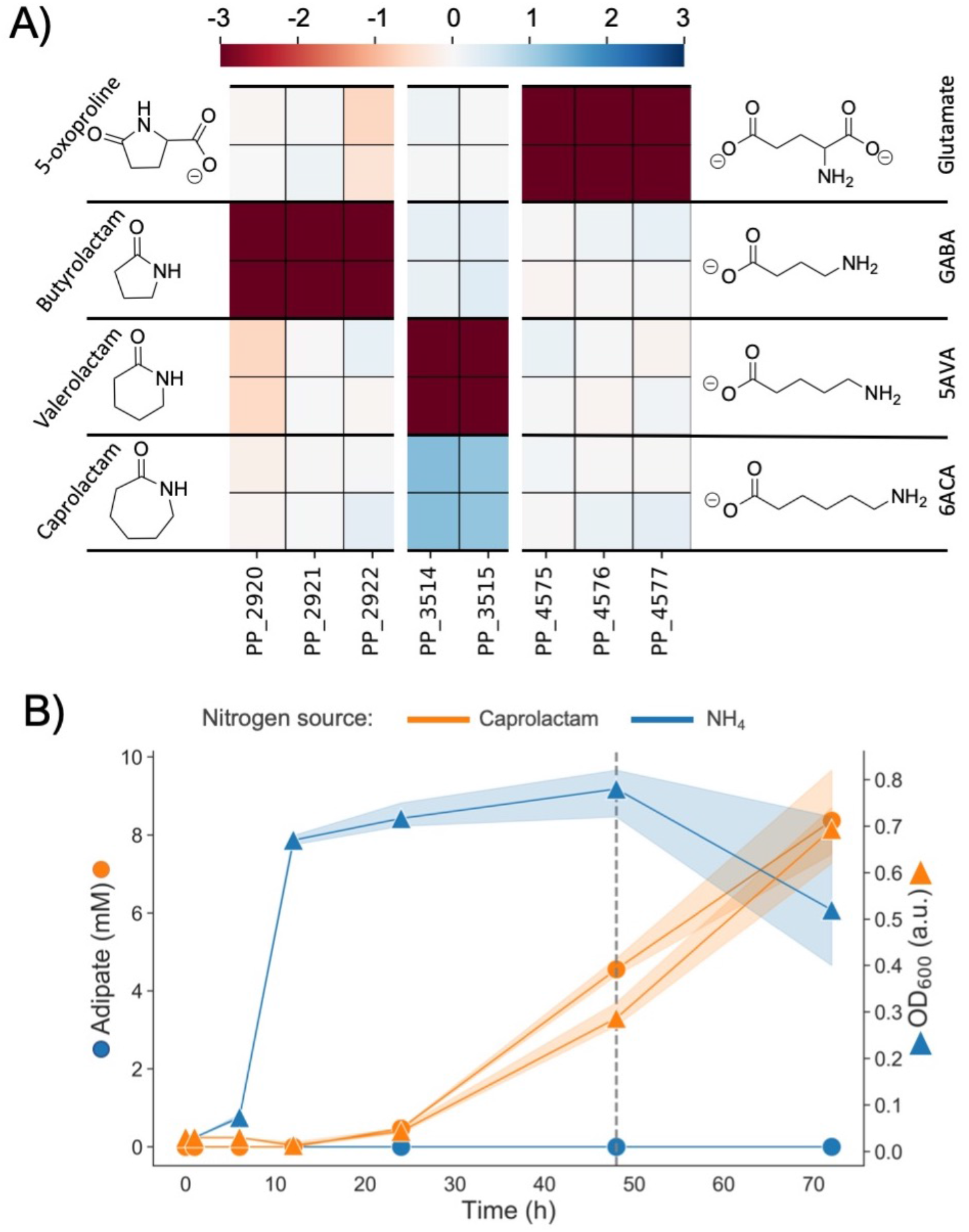
A) Heatmap with fitness scores (n=2) of genes putatively involved in the hydrolysis of caprolactam, valerolactam, butyrolactam and 5-oxoproline in *P. putida* KT2440. B) LC-MS analysis of caprolactam degradation in *P. putida* KT2440. Wild type cells were grown in MOPS minimal media with ammonium (blue) and caprolactam (orange) as the sole source of nitrogen. Shown is the OD (squares) and concentration of adipate in the supernatant (circles) over a time course of 72 hours. The dashed line marks the time point at which caprolactam was no longer detected in the media.

The three subunits of PP_2920-2 and PP_4575-7 are annotated by UniProt as paralogs of *pxpABC*, the previously described widespread prokaryotic 5-oxoprolinase (122). Formerly, 5- oxoprolinases were only known to exist in eukaryotes and bacteria possessing a γ-glutamyl cycle. Niehaus et al. hypothesized that prokaryotes must have a way to combat the spontaneous cyclization of glutamate and glutamine to 5-oxoproline and identified *pxpABC* (122). Our fitness data suggests that only PP_4575-7 acts on 5-oxoproline, although it is also possible that spontaneous metabolite cyclization is the reason for the evolution of *P. putida*’s two other lactam hydrolases, PP_2920-2 and the previously identified valerolactam hydrolase OplBA (PP_3514-5) (16). The ω-amino acids GABA and 5AVA are common metabolic intermediates in *P. putida*, appearing as part of arginine, putrescine, and lysine catabolisms (15, 123, 124). Previous work has demonstrated that these C4 and C5 ω-amino acids can cyclize following activation to an acyl-CoA thioester (16, 121, 125). With no functional hydrolase, these lactams would serve as dead-end metabolites, and the reduction in carbon and nitrogen availability might be detrimental to growth. To test whether these lactamase systems serve this purpose, we conducted growth assays of ΔPP_4575-7, ΔPP_2920-2, and Δ*oplBA* with their lactam substrate’s corresponding ω- amino acid as a nitrogen source (Figure S8). We observed slight growth lags and decreased maximal ODs in the knockouts versus the wild type, supporting the idea that these lactam hydrolases might exist for this purpose.

In previous work, the lactamase responsible for the hydrolysis of valero- and caprolactam, OplBA, was identified using proteomics data (16). Although the authors did search for the lactam hydrolase with BarSeq experiments using valerolactam as a sole carbon source, there was no significant fitness data for OplBA. Interestingly, we observed significant negative and positive fitness phenotypes for OplBA mutants with valerolactam and caprolactam as the nitrogen sources, respectively (Figure 6A). The positive fitness data for OplBA mutants on caprolactam is especially surprising, because previous work has shown that *oplBA* knockouts cannot grow with caprolactam as the sole nitrogen source (16). However, it could be explained by the nature of BarSeq experiments. Library mutants with a functional OplBA hydrolyze the caprolactam to 6ACA, which then accumulates due to *P. putida’s* slow utilization of this nitrogen source (Figure S2). Then, mutants with the *oplBA* genes disrupted can consume the freed 6ACA without needing to expend the resources to produce this protein in already nitrogen-limited growth conditions.

The slow utilization of caprolactam, and therefore 6ACA, as a nitrogen source may be because *P. putida* is unable to utilize the product of 6ACA transamination, adipic acid (126). We suspected that adipic acid was accumulating in the media, which we confirmed with a metabolomics experiment. Caprolactam was used as a sole nitrogen source, and samples of the supernatant were taken at 1, 6, 12, 24, 48, and 72 hours. Within 48 hours, caprolactam is no longer detectable in any of our samples, while adipic acid slowly accumulates at approximately the same rate that biomass increases (Figure 6B).

A possible explanation for the discrepancy between the fitness values of *oplBA* between valerolactam carbon and nitrogen source conditions is the growth kinetics of *P. putida*. Although there is no significant data for *oplBA* mutants when valerolactam is the carbon source, the aminotransferase that acts on 5AVA, PP_0214, still has a severe fitness defect (−4.1). It is likely that after valerolactam is hydrolyzed to 5AVA, it diffuses throughout the media and is effectively shared amongst the mutant library, eliminating the need for all library members to have a functional OplBA. However, cross-feeding would only be possible if *P. putida’s* consumption of 5AVA as a carbon source is slower than its transport out of the cell. In the valerolactam nitrogen source experiments, it is possible that mutants with a functional version of OplBA consume the provided glucose at such a rate that they also must deaminate the 5AVA much faster than it diffuses into the media. In this scenario, the inability to produce 5AVA from valerolactam would be detrimental, and *oplBA* mutants would find themselves nitrogen-starved, explaining the negative fitness value observed.

Adjacent to each of these three lactam hydrolases are three transcription factors: *oplR* (PP_3516), PP_2919, and PP_4579. The fitness data suggest that each of these transcription factors activates expression of its neighboring lactam hydrolase in response to the presence of its lactam substrate. However, we cannot tell from the fitness data if specificity for the different lactams is due to the transcription factor, the lactam hydrolase, or both. To answer this question, we examined the specificity of the lactam hydrolases through a complementation assay and the specificity of the transcription factors using RFP reporters.

For each of our three hydrolase knockout strains, we attempted to restore growth on its corresponding lactam with plasmid-based expression of the three different lactam hydrolases, resulting in a total of 9 strains tested. If a lactam hydrolase was more promiscuous than its regulator, we would expect it to restore growth in backgrounds with a different lactamase knocked out. However, plasmid-based expression of the lactam hydrolases only restored growth in the expected nitrogen source condition, indicating that the lactam hydrolases are fairly specific (Figure S9).

The transcription factor *oplR* has already been identified as the regulator of *oplBA* and developed into a suite of highly sensitive valero- and capro- lactam biosensor plasmids (10). However, although it is strongly suggested, we cannot say from the fitness data alone that the other adjacent regulators are responsible for inducing lactamase expression. To confirm the role of each transcription factor, we conducted assays of two-plasmid systems in *E. coli*, as done by Thompson et al with *oplR* (10). The transcription factor was expressed under control of an arabinose inducible promoter on a low copy plasmid (pSC101 origin), while the region 200 bp upstream of the lactamase was cloned 5’ of the RFP gene in a medium copy plasmid (BBR1).

We refer to these plasmids as the sensor and reporter plasmid, respectively. The transcription factor induction level and concentration of the expected lactam ligand were then combinatorially varied. RFP, normalized with OD, increased with increasing lactam concentrations, and the dynamic range of the assay changed with transcription factor induction (Figure S10B). This indicated that the transcription factors do in fact induce expression of their adjacent lactam hydrolases in response to their corresponding lactams. However, total induction and dynamic range were very small, likely because these transcription factors do not function optimally in *E. coli*. Therefore, we decided to probe the specificity of lactam regulation in *P. putida*.

Unfortunately, our sensor plasmids are not compatible with *P. putida*, and swapping the lactam hydrolase for RFP in the genome also did not result in sufficient fluorescent signal (data not shown). Instead, we opted to test the ligand range of these transcription factors by transforming our reporter plasmid into *P. putida* and relying on native expression levels of the transcription factors.

Both the butyrolactam and 5-oxoproline systems displayed high inducer specificity, with RFP induction occurring only in the presence of their expected ligand. The dynamic range of both of our reporter plasmid systems was quite wide, and both systems displayed high sensitivity. In particular, the butyrolactam reporter had normalized RFP signal roughly 900 times above zero induction at the lowest tested butyrolactam concentration (50 μM). This system has the potential to be highly effective as a biosensor for detecting butyrolactam production in *P. putida*. We also observed a response to butyrolactam from the reporter plasmid for OplR. Previous work by Thompson et al. showed that OplR is highly specific to valero- and capro- lactam, but the maximum butyrolactam concentration tested was 0.5 mM and the host was *E. coli* (10). We detected fluorescence at butyrolactam concentrations greater than ∼ 6.25 mM, however, since we are relying on native expression of the transcription factor, we cannot know for certain whether this is due to a response from OplR to the butyrolactam or crosstalk between PP_2919 and the *oplBA* promoter (Figure S10C).

### Polyamines, ω-amino acids, and GABA isomers

Polyamines are abundant in nearly all forms of life and are involved in numerous cellular functions, including gene expression, stress response, cell growth, and membrane homeostasis (127). Beyond their biological significance, polyamines also have great industrial value, for example, putrescine is a direct nylon precursor, and propanediamine is used in textile finishing and strengthening agents (128, 129). The ω-amino acids, several of which are the degradation products of polyamines, also have both biological and industrial relevance. Many are produced as metabolic intermediates in pathways such as proteinogenic amino-acid degradation, while some, such as GABA, can act as signaling molecules (130, 131). Industrially, the aforementioned 6ACA, 5AVA, and GABA are precursors to several polymers and plastics (121).

The three transaminases that appear to be most important for the utilization of polyamines, ω-amino acids, and GABA isomers are the pyruvate:alanine aminotransferases PP_0596, PP_2180 (SpuC-I), and PP_5182 (SpuC-II). They have different specific fitness phenotypes on both polyamines and amino acids, however, because the tested polyamines are converted to ω-amino acids, we could not determine if any acted directly on polyamines.

Therefore, we purified these enzymes to assay their substrate range *in vitro*. Surprisingly, all three of the aminotransferases showed some activity on nearly all of the substrates tested, including substrates on which they had no significant fitness phenotype (Table 4) (Figure S11).

**Table 4:**
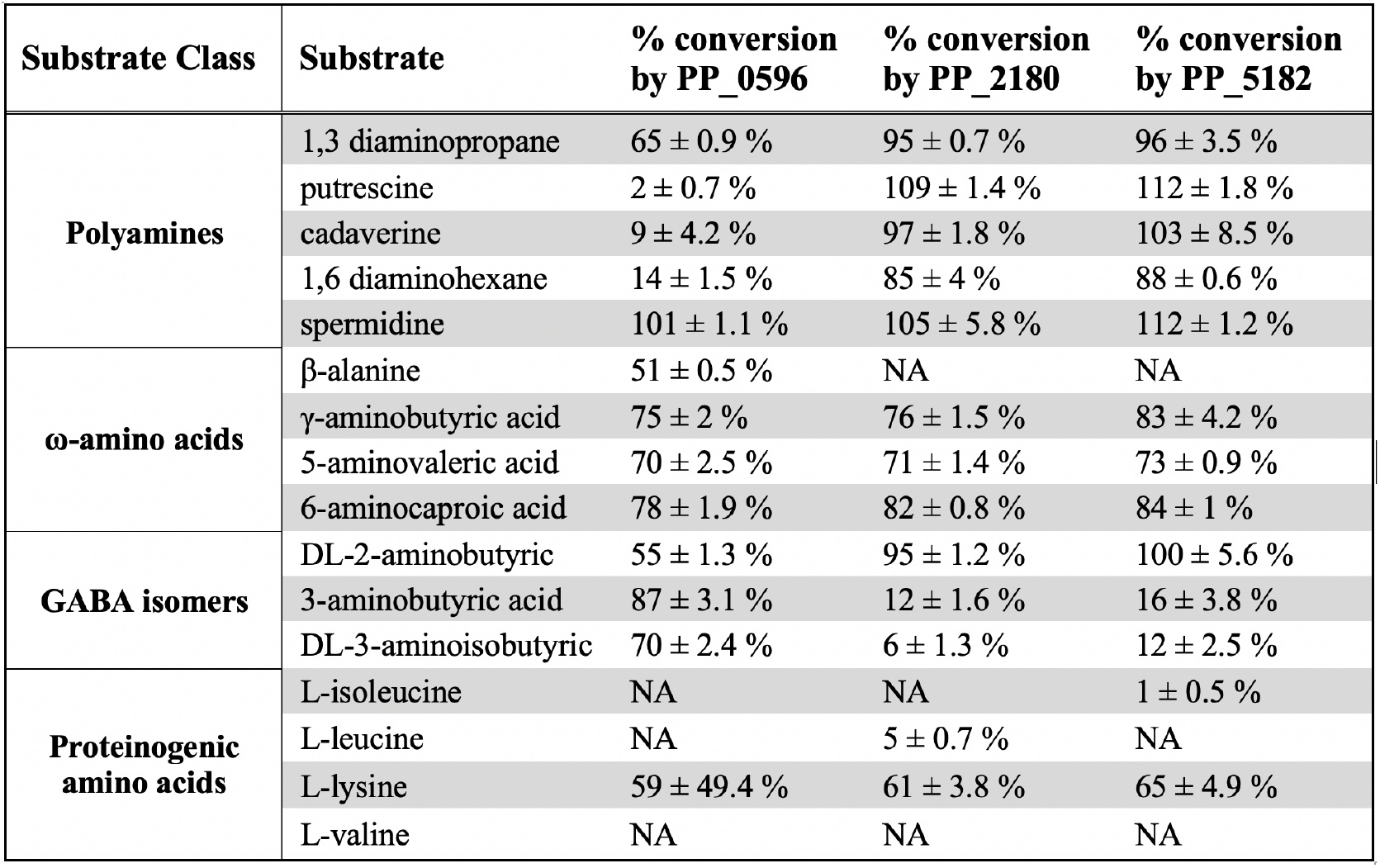
Substrate range of purified pyruvate:alanine aminotransferases. Tested substrates were added in excess to pyruvate. Activity of the aminotransferases on the tested substrates was measured indirectly through the quantification of the co-product alanine. Percent conversion is relative to the expected measurement of alanine if the amine acceptor pyruvate was completely transformed. Error represents the standard deviation of three replicates. NA indicates that no activity was statistically detected above controls.

The promiscuity of these enzymes directly contrasts with the specific phenotypes of the fitness data. For example, PP_2180 or PP_5182 have no fitness defect on 1,3-diaminopropane (1,3-DAP), yet both demonstrate greater activity on this substrate than PP_0596, which does have a significant fitness defect. This disparity between the fitness data and biochemical data is best explained by a difference in the promiscuity of these enzymes and their regulators. To investigate our hypothesis, we constructed fluorescent reporter systems.

As previously described in the lactam section, we used a two-plasmid approach with RFP as an output to test transcription factor and aminotransferase promoter combinations in *E. coli*. Based upon our fitness data and recently published DAP-Seq data from *P*. *aeruginosa*, we hypothesized that the transcription factor PP_5268 may regulate PP_5182 (132). However, no fluorescence signal was detected when testing the sequences upstream of PP_5182 or PP_5183 as promoters (data not shown). It is possible that the tested transcription factor-promoter systems are incompatible in *E. coli* or that PP_5268 is not responsible for the regulation of PP_5182. Nevertheless, we were able to identify the transcription factors likely responsible for regulation of the aminotransferases PP_0596 and PP_2180 (Figure S12). Adjacent to PP_0596 is the LysR-type transcription factor PP_0595, which has fitness data that closely mirrors the phenotypes of PP_0596. The two-plasmid system for this combination worked as anticipated, and it appears that PP_0595 activates the promoter in response to both DL-3-aminoisobutyric acid (3AIBA) and β-alanine (Figure S12). The aminotransferase PP_2180 had opposite fitness data to the adjacent MerR-family transcription factor PP_2181, which is oftentimes characteristic for transcriptional repressors. We first tested PP_2181 with the region 200 base pairs upstream of PP_2180 as a promoter, but detected no significant increase in fluorescence (data not shown). However, PP_2180 appears to be part of an operon and using the region upstream of PP_2177 as a promoter results in a response to 1,6-diaminohexane (1,6-DAH) and cadaverine (Figure S12).

The specificity of these transcription factors was much higher than the *in vitro* substrate specificity of the aminotransferases they regulate, suggesting that the role of PP_0596 and PP_2180 *in vivo* is determined by regulation.

Utilization of polyamines as a nitrogen source can proceed through two pathways, direct transamination by an aminotransferase or the γ-glutamylation pathway. However, because the product of the γ-glutamylation pathway is an ω-amino acid that is also then a substrate for an aminotransferase, we cannot say with certainty which pathways are used based on fitness data alone. In 1,6-DAH and cadaverine nitrogen source conditions, we see mild-fitness defects (< - 1.0) for the two aminotransferases PP_5182 and PP_2180. As previously discussed, both aminotransferases act on these substrates *in vitro* and the regulator PP_2181 responds specifically to 1,6-DAH and cadaverine. This suggests that transamination, not γ-glutamylation, is the preferred method of utilization for longer chain polyamines. In-frame deletions of either aminotransferase alone does not effectively abolish growth on 1,6-DAH and cadaverine, likely because they are functionally redundant. Strains with both PP_5182 and PP_2180 deleted show almost no growth with cadaverine and 1,6-DAH as sole nitrogen sources, although we did observe very slight growth in the double knock out strain after 72 hours (Figure S13). This could be due to the background activity of another aminotransferase or the γ-glutamylation pathway.

Conversely, the γ-glutamylation pathway appears to be equally if not more important than direct transamination for the utilization of putrescine, 1,3-diaminopropane (1,3-DAP), and spermidine, with each condition resulting in significant fitness phenotypes for all steps of the pathway (Figure 7B). It is possible that putrescine, 1,3-DAP, and spermidine may also be directly deaminated by aminotransferases, although this is less likely for putrescine, which results in no significant fitness phenotypes for any aminotransferases.

**Figure 7:**
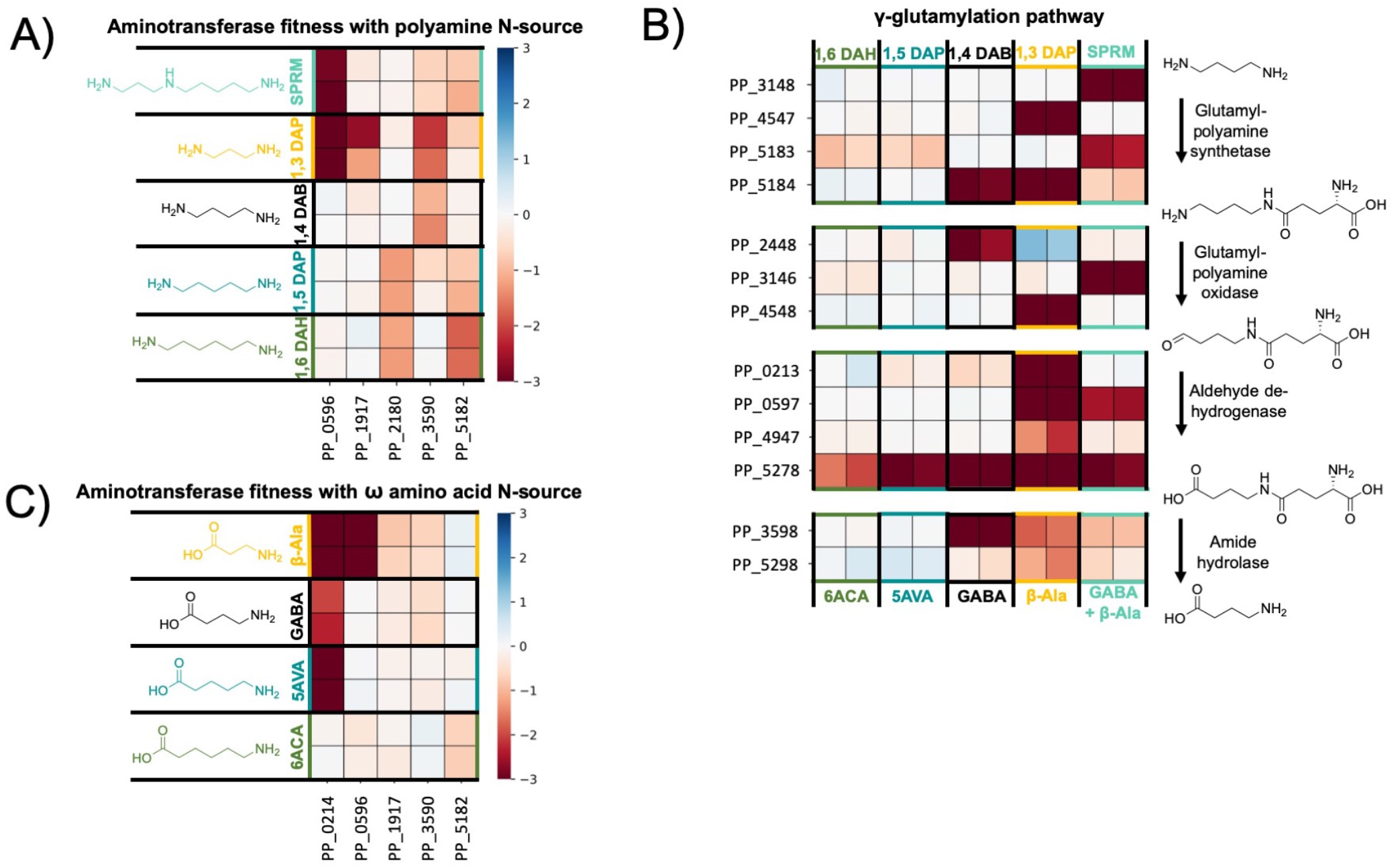
Heatmaps of polyamine and ω-amino acid catabolism in *P. putida.* For each enzyme class shown, HMMER was used to identify all putative genes present in *P. putida* corresponding to that pFam in *P. putida*. Data was filtered to find genes in each class with fitness values >-1.0 and t > |4| in the nitrogen sources shown. A) Heatmap with fitness values (n=2) for genes putatively involved in polyamine transamination in *P. putida* KT2440. B) Heatmap with fitness values (n=2) for genes putatively involved in polyamine γ-glutamylation in *P. putida* KT2440. C) Heatmap with fitness values (n=2) for genes putatively involved in omega-amino acid transamination in *P. putida* KT2440. Spermidine (SPRM) = teal; 1,3 diaminopropane (1,3 DAP) = yellow; β-alanine = yellow; putrescine/1,4-diaminobutane (1,4 DAB) = black; γ-aminobutyric acid (GABA) = black; cadaverine/1,5- diaminopentane (1,5 DAP) = blue; 5-aminovalerate (5AVA) = blue; 1,6-diaminohexane (1,6 DAH) = green; 6- aminocaproic acid (6ACA) = green.

The ω-amino acids, whether fed directly or released from polyamine catabolism, are degraded by transamination to an aldehyde then oxidation to a dicarboxylic acid. The most poorly utilized ω-amino acid we tested is 6ACA, which typically results in little to no visible growth until close to 72 hours. This is likely due to a combination of regulatory issues and the accumulation of its product adipic acid, which *P. putida* is unable to utilize further (Figure 6B). The *spuC* paralogs PP_5182 and PP_2180 are likely involved in 6ACA catabolism, despite their weak (> -1) phenotypes. Deleting either of these aminotransferases individually does not fully abolish growth on 6ACA, however, a double knockout strain is unable to grow on 6ACA (data not shown). If the aminotransferases for which we tested the *in vitro* substrate range are just as promiscuous *in vivo* and regulatory issues do contribute to the slow growth rate of 6ACA, we would expect that plasmid-based expression of these aminotransferases would improve growth. To test this theory, we provided functional gene copies to knockout mutants of PP_0596, PP_5182, and PP_2180 by introducing the arabinose-inducible pBADT plasmid harboring the corresponding gene. (Figure S14). Surprisingly, the control strain, wild-type *P. putida* harboring pBADT-RFP, was unable to grow under these culture conditions. We hypothesize that the burden of plasmid maintenance and insufficient kanamycin-resistance protein levels lead to early stage cell death when 6ACA is used as the sole source of nitrogen. However, our functionally complemented strains all demonstrated growth on 6ACA, indicating that that regulation contributes to the slow utilization of 6ACA and expressing any of the three aminotransferases with known activity on 6ACA independently of native regulation improves growth.

As previously discussed, the transcription factor PP_0595 is specific for β-alanine, and the main substrate of the aminotransferase it regulates, PP_0596, appears to be β-alanine *in vivo*, although we also found it necessary for utilization of uracil, 3ABA, and 3AIBA (Figure S15). Both of these genes have a very strong fitness defect in the β-alanine nitrogen source condition (− 7.8). Located in the same operon is the methylmalonate-semialdehyde dehydrogenase PP_0597 (−5.9), which catalyzes the conversion of malonate-semialdehyde to acetyl-CoA. The requirement for PP_0596, *davT*, and *gltBD* in the β-alanine condition may indicate a higher-order metabolic pathway for efficient metabolism of β-alanine. In the β-alanine condition, DavT may catalyze the reverse reaction (succinate-semialdehyde and glutamate to 4-ABA and 2- oxoglutarate) to reduce intracellular alanine concentrations produced by PP_0596. This is supported by the requirement of DavD (PP_0213, succinate/glutarate dehydrogenase) (Figure 2B). Alternatively, DavT and PP_0596 may both have activity on β-alanine, and a knockout in either gene could result in detrimental loss of flux to malonate-semialdehyde. This peculiarity of β-alanine metabolism requires further investigation.

*In P. putida*, GABA can serve as the sole source of both carbon and nitrogen (133). Since GABA is abundantly present in root exudates, *P. putida* possesses sensitive and specific receptors that detect GABA and facilitate chemotaxis (133, 134). Unlike *E. coli*, the genes for the degradation and transport of GABA are not clustered in a single operon (70, 135).

Furthermore, the observed fitness defects are mostly unspecific, which makes it more difficult to identify distinct phenotypes. However, the main aminotransferase acting on GABA appears to be DavT (PP_0214; -2.15), producing succinate-semialdehyde that is then converted to succinate by DavD (70). The fitness defect for *davD* (−0.45) is not significant (|t| < 5) suggesting that additional dehydrogenases such as PP_2488 (0.0) or PP_3151 (−0.2) catalyze this reaction.

Moreover, our biochemical assay revealed that at least three other aminotransferases (PP_0596, PP_2180, PP_5182) accept GABA as a substrate (Table 4). Although the second identified aminotransferase (PP_3590) by BarSeq has an insignificant phenotype (−0.55; |t| = 3.7), it exhibits a significant fitness defect in the DL-2-aminobutyric acid condition (−0.95).

2-aminobutyrate (2-ABA) is an unnatural amino acid that is used in the production of pharmaceuticals, and has been a target molecule for biological production in *E. coli* (136, 137). The only known usage of 2-ABA in biological systems is the substitution of L-cysteine during glutathione synthesis to form ophthalmic acid (138). Biosynthetically, it can occur during isoleucine synthesis via threonine by IlvE transamination of 2-oxobutyrate (137). Due to the essentiality of *ilvE* in minimal media conditions, we are not able to confirm this in *P. putida* using BarSeq. However, we were able to identify the D-amino acid oxidoreductase PP_5270 (− 2.75) which acts on the alanine produced during pyruvate transaminase activity. Although there is no significant phenotype for PP_0596 in this condition, we have shown that PP_0596 can transaminate 2-ABA (Table 3). Furthermore, the transamination product 2-oxobutyrate is an intermediate in isoleucine biosynthesis. Because of this, there appears to be a requirement for the downstream steps of isoleucine biosynthesis. Instead of producing isoleucine, its precursor (S)-3- methyl-2-oxopentanoate might be directly funneled into the isoleucine degradation pathway via the branched-chain ɑ-ketoacid dehydrogenase (BCKD) complex (−1.54). This decarboxylative route is further supported by the requirement of the methylcitrate cycle (PP_2334-6; -1.5). These genes are important for the degradation of propionyl-CoA, which can only be produced from 2- ketobutyrate via a decarboxylative reaction (139).

The last GABA isomer we tested was 3ABA, previously described in the pyrimidine section. 3ABA is one of several GABA isomers that can prime plant immunity, and it is applied agriculturally to prevent crop blight (140). Although the most effective route for 3ABA degradation seems to be the pyruvate transaminase, PP_0596 (−5.25), it appears that the cotranscribed gene, PP_0597 (malonyl semialdehyde dehydrogenase), is deleterious during growth on 3ABA (2.85). The dehydrogenase PP_0597 is not the correct enzyme for further processing of the downstream product acetoacetate, and is likely a waste of resources.

Furthermore, the reaction catalyzed by PP_0597 is identical to that of PP_4667, and it is possible that expression of PP_0597 results in a metabolic disturbance in valine metabolism.

## FUTURE DIRECTIONS

In this work, hundreds of genes critical for growth of *Pseudomonas putida* KT2440 on diverse nitrogen sources were identified. This provides a roadmap for the characterization of other microbes using functional genomics. Deeper analysis of related γproteobacteria through similar omics methodologies could enable investigations into the evolution and diversification of these metabolic phenotypes. The Pseudomonas genus alone contains a litany of species with interesting metabolisms. Further interrogation of this genus through BarSeq or multi-omics could shine a light on the evolutionary and ecological importance of these metabolisms.

The majority of our functional genomics data validates what has already been predicted by major databases such as KEGG or SEED. We and others have confirmed the reliability of TnSeq data as a guide for successful metabolic engineering (11, 14, 16, 29). These data provide a level of *in vivo* evidence for gene function that is distinct from prior mutational interrogations.

We contend that these data are highly predictive and can be used in conjunction with existing databases (BioCyc, SEED, KEGG) to inform genetic, biochemical, and bioengineering studies. A process for integrating the acquired information from these and other TnSeq studies into public-facing databases would be desirable.

The identification of genetic interactions through t-SNE and correlation analyses could also enable more detailed interrogations of their translated proteins. With the recent publication of Alphafold2 and RoseTTA fold, it is now possible to elucidate structures and protein-protein interfaces using purely *in silico* approaches (141, 142). Using computer-generated protein structures and docking simulations, one could propose and test more detailed hypotheses regarding structure-function relationships. This could further deepen our understanding of the molecular mechanisms that enable these diverse phenotypes.

## METHODS

### Media, chemicals, and culture conditions

General *E. coli* cultures were grown in lysogeny broth (LB) Miller medium (BD Biosciences, USA) at 37 °C while *P. putida* was grown at 30 °C. When indicated, *P. putida* and *E. coli* were grown on modified MOPS minimal medium, which is comprised of 32.5 µM CaCl2, 0.29 mM K2SO4, 1.32 mM K2HPO4, 8 µM FeCl2, 40 mM MOPS, 4 mM tricine, 0.01 mM FeSO4, 9.52 mM NH4Cl, 0.52 mM MgCl2, 50 mM NaCl, 0.03 µM (NH4)6Mo7O24, 4 µM H3BO3, 0.3 µM CoCl2, 0.1 µM CuSO4, 0.8 µM MnCl2, and 0.1 µM ZnSO4 (102). For most experiments, nitrogen-free MOPS was used, in which case the NH4CL was omitted and the nitrogen source of interest was added at concentrations ranging from 4-10 mM. Cultures were supplemented with kanamycin (50 mg/L, Sigma Aldrich, USA), gentamicin (30 mg/L, Fisher Scientific, USA), or carbenicillin (100 mg/L, Sigma Aldrich, USA), when indicated. All other compounds were purchased through Sigma Aldrich (Sigma Aldrich, USA).

### Strains and plasmids

The strains and plasmids used in this work are listed in Table 5, and plasmids used in this work are listed in Table 6. All strains and plasmids created in this work are available through the public instance of the JBEI registry (public-registry.jbei.org/folders/456). Device Editor and Vector Editor software were used to design the plasmids, and primers used for the construction of plasmids were designed using j5 software (143–145). All primers were purchased from Integrated DNA Technologies (IDT, Coralville, IA). Plasmids were assembled via Gibson Assembly using standard protocols and isolated with the Qiaprep Spin Miniprep kit (Qiagen, USA) (146).

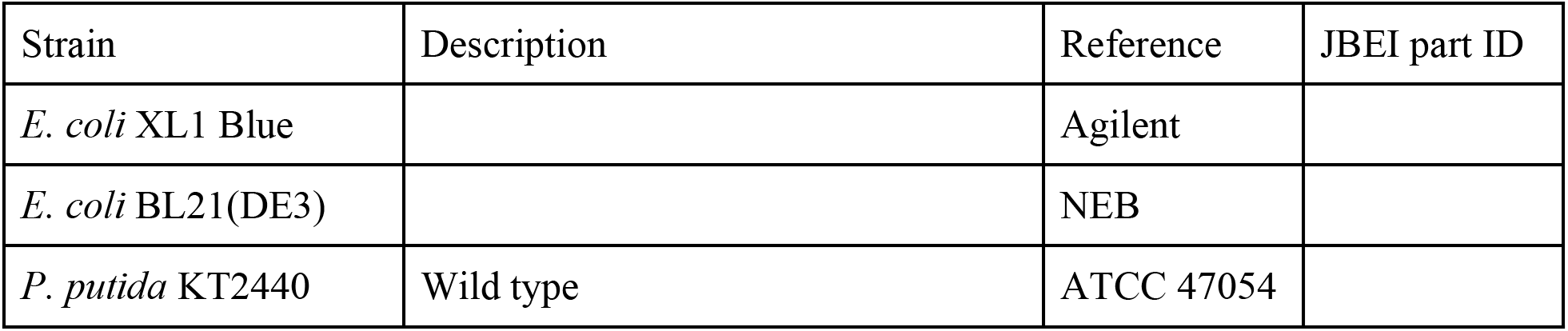
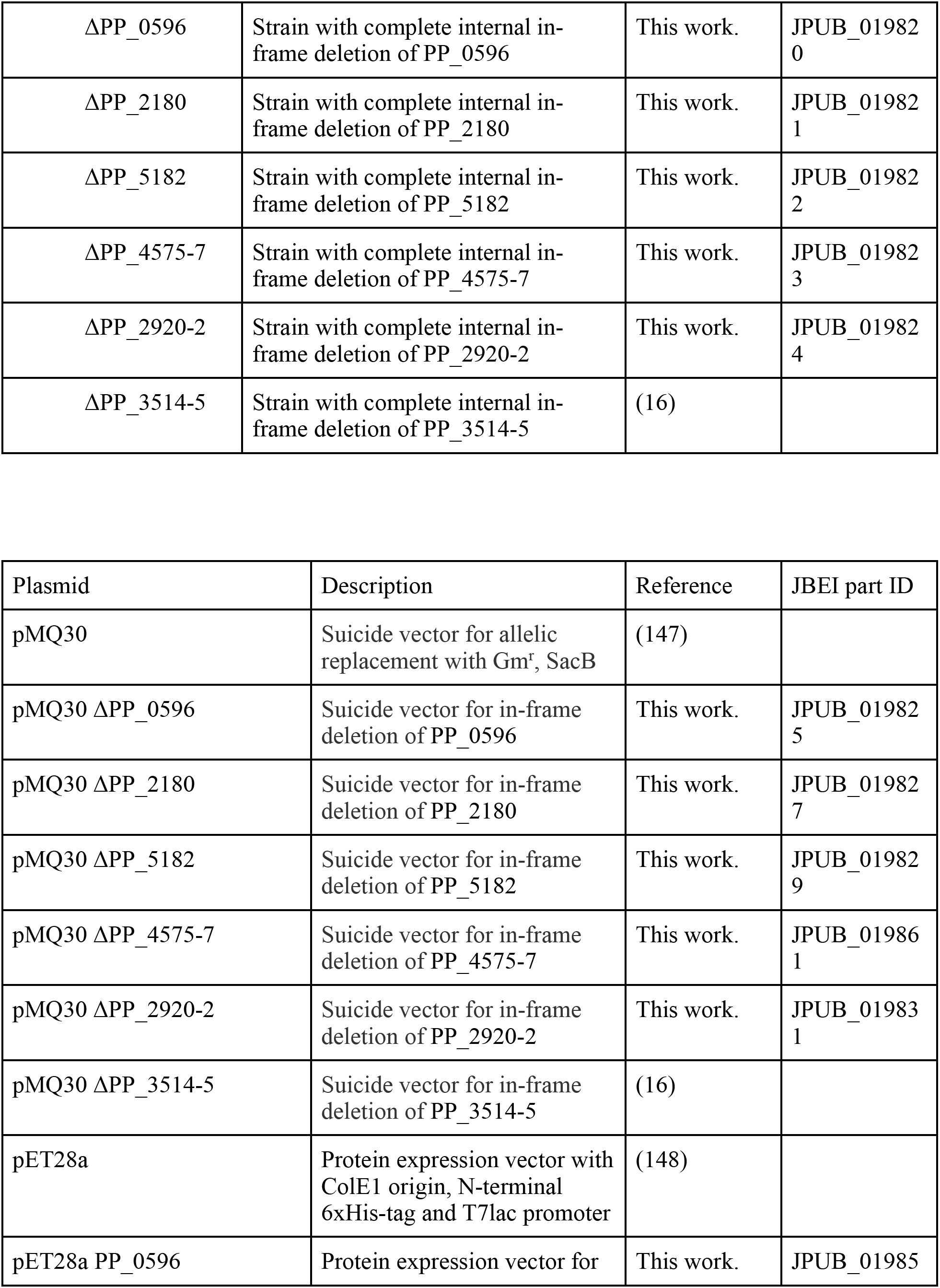
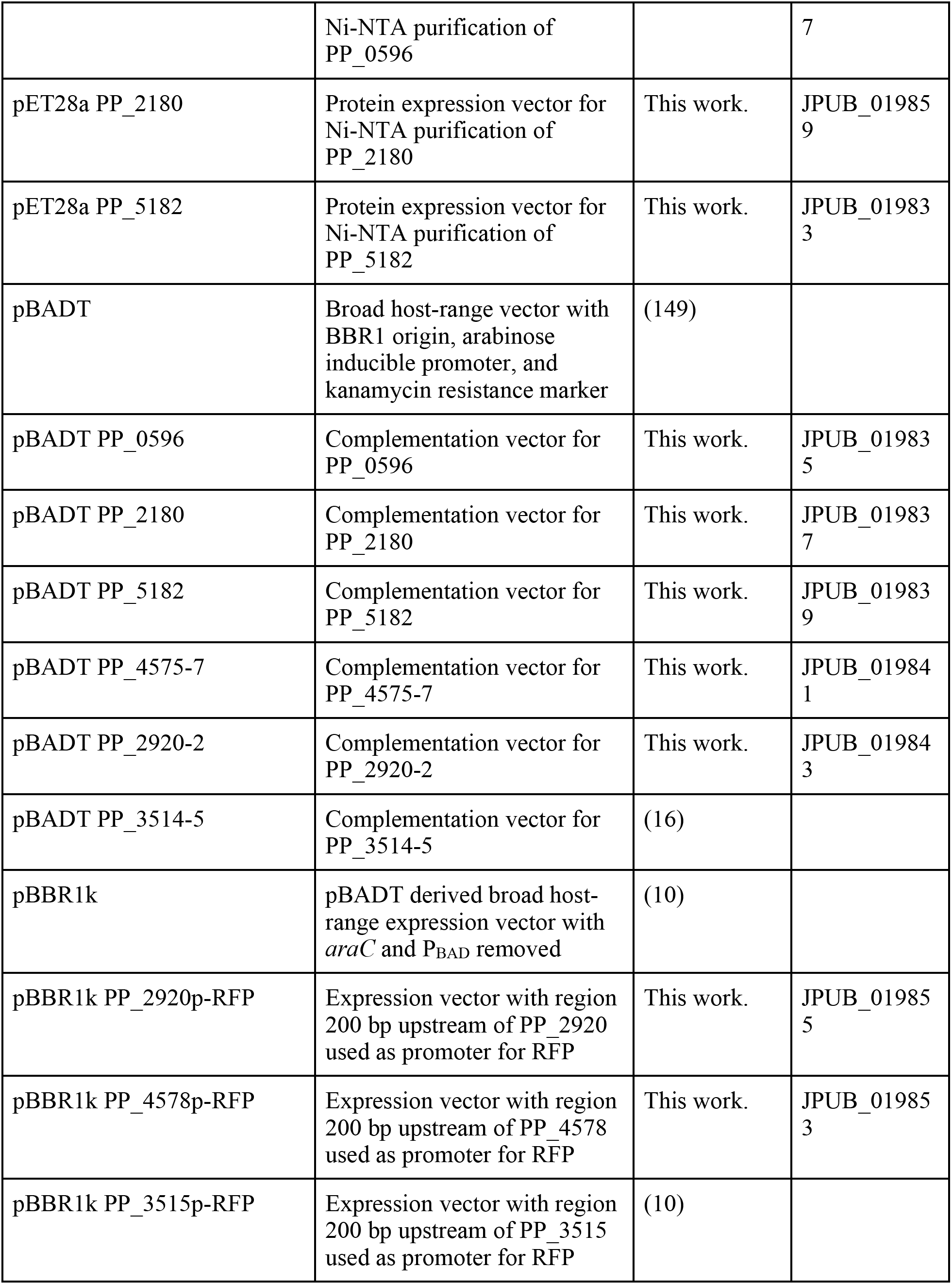
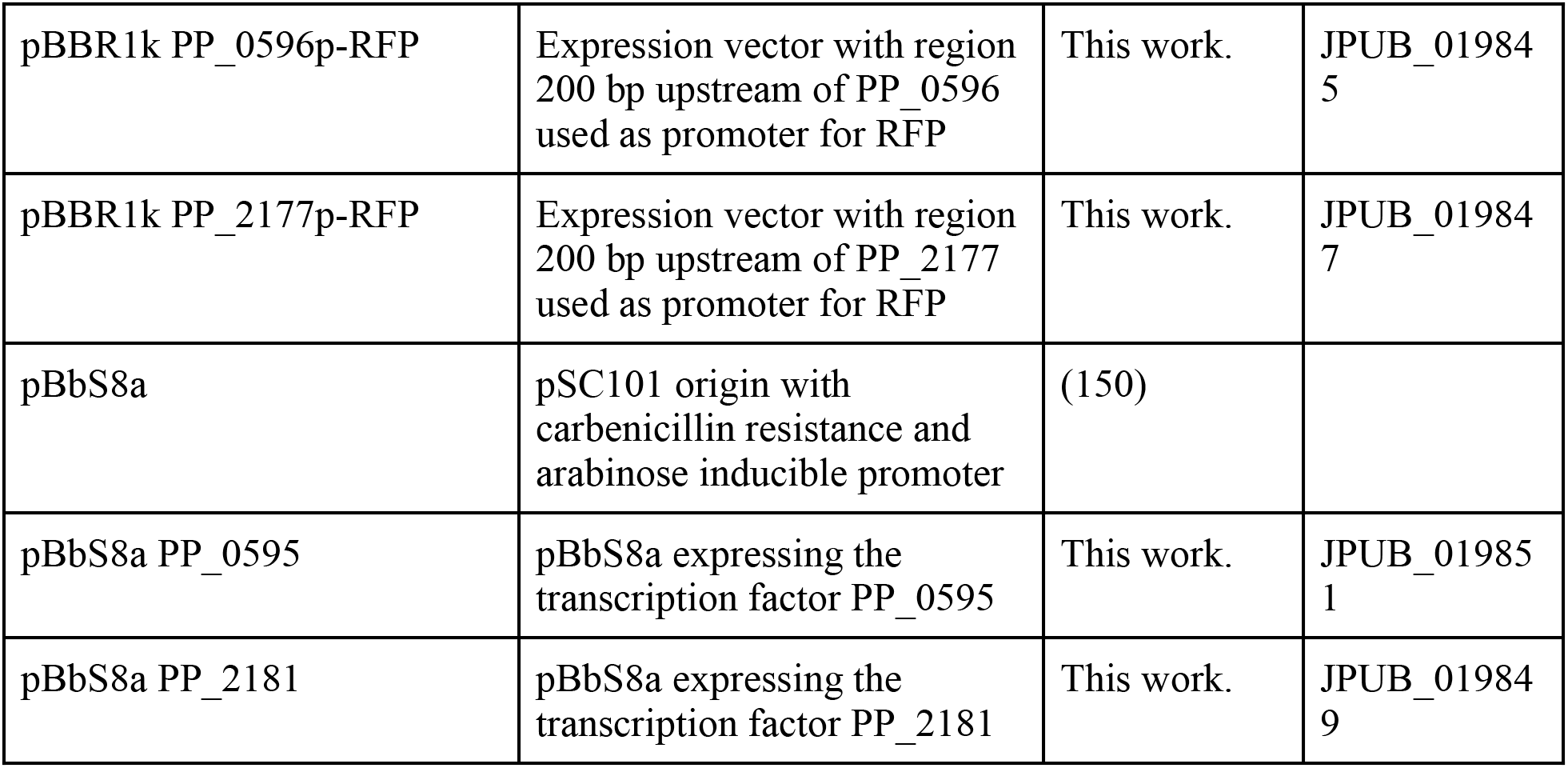

### Plate-based growth assays

Growth studies of bacterial strains were conducted using microplate reader kinetic assays as described previously (10). Overnight cultures were washed three times with nitrogen-free MOPS minimal medium and used to inoculate 48-well plates at a ratio of 1:100 (Falcon, 353072). Each well contained 500 μL of MOPS medium with 10 mM of the tested nitrogen source. Plates were sealed with a gas-permeable microplate adhesive film (VWR, USA), and then optical density and fluorescence were monitored for 24-72 hours in an Biotek Synergy H1M plate reader (BioTek, USA) at 30 °C with fast continuous shaking. Optical density was measured at 600 nm. RFP was measured with an excitation wavelength of 535 nm, an emission of 620 nm, and a gain of 100.

### BarSeq assays

BarSeq experiments utilized the *P. putida* library JBEI-1 and were completed as previously described (15). Aliquots (2 mL) of libraries of JBEI-1 were thawed on ice, added to 25 mL of LB medium with kanamycin, and then grown at 30 °C to an OD600 of 0.5. Then, three 1-mL aliquots were removed, pelleted, and stored at -80 °C as zero timepoints. The libraries were washed three times in MOPS minimal medium with no nitrogen source, and then used to inoculate each experiment at a ratio of 1:100. Experiments were conducted in 24-well plates; each well contained 2 mL of nitrogen-free MOPS minimal medium with 10 mM of each tested nitrogen source. Plates were grown at 30 °C with shaking at 200 rpm, and 1 mL samples were collected after 24-72 hours, depending on when cultures appeared sufficiently turbid for DNA extraction. Samples were pelleted and stored at -80 °C until DNA extraction, which was done with a DNeasy UltraClean Microbial kit (Qiagen, Germany). BarSeq analysis was performed as previously described (19, 40). Strain fitness is defined as the normalized log2 ratio of the barcode reads in the experimental sample to the barcode reads in the time zero sample. The fitness of a gene is defined as the weighted average of the strain fitness for insertions in the central 10% to 90% of the gene. The gene fitness values are normalized such that the typical gene has a fitness of zero. The primary statistic t value represents the form of fitness divided by the estimated variance across different mutants of the same gene. Statistic t values of >|4| were considered significant. A more detailed explanation of fitness score calculations can be found in Wetmore et al. (151). All experiments described here passed the quality testing metrics described previously. Experiments were conducted in biological duplicates, and the fitness data are publically available at http://fit.genomics.lbl.gov.

### Protein Production and Purification

The protocol for the ⍵-amino acid aminotransferases purification is a modified version of the previously described procedure by (152). Briefly, *E. coli* BL21(DE3) harboring the pET28a vectors for PP_0596, PP_2180 or PP_5182 were cultured in LB medium with kanamycin at 37 °C until OD600 reached 0.4. Protein expression was induced with 250 µM IPTG and cultures were grown at 18 °C for 24 h. Cells were harvested by centrifugation for 20 min at 5000*g* and the pellet was resuspended in 30 mL wash buffer (50 mM sodium phosphate pH 7.6, 300 mM NaCl, 10 mM imidazole, 4 °C). The cells were disrupted by sonication (8 x 30 sec) and cell debris was removed by three subsequent centrifugations (15 min, 8000*g*, 4 °C). The soluble fraction was mixed with 4 mL Nickel-NTA agarose beads (Thermo Fisher Scientific) for 1 h at 4 °C. The mixture was applied to a Nickel-NTA column and washed three times with wash buffer. The protein was eluted with 12 mL elution buffer (150 mM sodium phosphate buffer pH 7.6, 50 mM NaCl, 150 mM imidazole, 4 °C) and buffer was exchanged with stock buffer (100 mM sodium phosphate pH 7.5, 0.5 mM DTT, 10 % glycerol, 4 °C) by dialysis (SnakeSkin Dialysis Tubing, 10k MWCO, Thermo Fisher Scientific). Protein was concentrated in 30k MWCO Amicon Ultra-15 centrifugal filters (MilliporeSigma) and stored at -80 °C.

### *In vitro* pyruvate transamination and product quantification

Substrate specificity of the purified aminotransferases was determined by *in vitro* L-alanine production. The reaction was carried out in 100 mM sodium phosphate buffer pH 9 with 500 µM pyruvate, 1 mM PLP, 5 µM enzyme and 5 mM substrate. After addition of the substrate, the reaction mixture was immediately incubated at 30 °C for 30 min. The reaction was stopped by boiling the mixture for 10 min at 100 °C. Alanine concentrations were determined enzymatically using the Alanine Assay Kit (Cell Biolabs, Inc.) following the manufacturer’s instructions.

### Caprolactam degradation assay and LC-ESI-QTOF-MS analysis

To investigate the degradation of caprolactam in *P. putida* KT2440, wild type cells were grown in modified MOPS minimal media with either 10 mM ammonium chloride or caprolactam as the sole source of nitrogen. Cells were cultured as triplicates in 25 mL medium in shake flasks at 30 °C and 200 rpm. Samples were collected and OD600 was measured after 1, 6, 12, 24, 48 and 72 hours. For metabolomics analysis, cell culture samples were centrifuged at 16,000*g* for 1 min, and 300 µL supernatant was mixed with -80 °C cold methanol. After that the mixture was filtered by centrifugation (30 min, 14.000*g*, 4 °C) using 3 kDa Amicon Ultra-0.5 centrifugal filters (MilliporeSigma) and stored at -80 °C for further analysis.

For the measurement of caprolactam, liquid chromatographic separation was conducted with a Kinetex HILIC column (100-mm length, 4.6-mm internal diameter, 2.6-µm particle size; Phenomenex, Torrance, CA) using a 1260 HPLC system (Agilent Technologies, Santa Clara, CA, USA). The injection volume for each measurement was 2 µL. The sample tray and column compartment were set to 6 °C and 20 °C, respectively. The mobile phase was composed of 10 mM ammonium formate and 0.2% formic acid in water (solvent A) and 10 mM ammonium formate and 0.2% formic acid in 90% acetonitrile and 10% water (solvent B). Reagent grade and source are as follows: Ammonium formate (Analytical grade) and formic acid (98-100% chemical purity): Sigma-Aldrich, St. Louis, MO, USA. Acetonitrile (LC-MS grade) and water (LC-MS grade): Honeywell Burdick & Jackson, CA, USA. Analytes were separated with the following gradient: 90% to 70%B in 4 min, held at 70%B for 1.5 min, 70% to 40%B in 0.5 min, held at 40%B for 2.5 min, 40% to 90%B in 0.5 min, held at 90%B for 2 min. The flow rate was varied as follows: held at 0.6 mL/min for 6.5 min, linearly increased from 0.6 mL/min to 1 mL/min in 0.5 min, and held at 1 mL/min for 4 min. The total run time was 11 min.

The 1260 HPLC system was coupled to an Agilent Technologies 6520 QTOF-MS system (for quadrupole time-of-flight mass spectrometric detection). The LC column effluent was delivered to the 6520 QTOF-MS system’s electrospray ionization (ESI) ion source via a 1:4 post-column split ratio. ESI was used to facilitate the production of gas-phase [M + H]^+^ ions in the positive ion mode. The capillary voltage was set to 3500 V. Fragmentor, skimmer, and OCT 1 RF voltages were set to 100 V, 50 V, and 250 V, respectively. Drying gas temperature, drying gas flow rate, and nebulizer were set to 350 °C, 12 L/min, and 25 lb/In^2^, respectively. The instrument was tuned for a range of m/z 50 to 1,700 with the Agilent ESI-Low TOF tuning mix. High mass accuracy was achieved via reference mass correction with purine and HP-0921 reference solutions, which were purchased from Agilent Technologies. Data was acquired within the 50 to 1100 m/z range. Analytes were quantified via a six-point calibration curve following 2-fold serial dilutions from 25 µM to 0.78125 µM. MassHunter Workstation (Agilent Technologies), MassHunter Qualitative Analysis and MassHunter Profinder (Agilent Technologies) software were used for data acquisition and processing.

The LC-ESI-TOF-MS method for adipic acid has been described previously (153).

### t-Stochastic Neighbor Embedding of BarSeq results

The data used for the t-SNE visualization was filtered by t-scores and fitness scores (|t| > 5 and |fitness| > 1). This resulted in a file containing fitness scores for 615 significantly affected genes across 129 sole-nitrogen source growth assays in 51 different conditions, and 19 amino acid dropout conditions. 2-ABA was excluded from the analysis due to the large number of significantly affected genes in this condition. For each gene, the “significant condition” was defined as the condition that caused the largest change in fitness score. The fitness values were then used as the input into the t-SNE module supplied in the python package Scikit-Learn (19, 154). Clusters (n=100) were defined using the Ward-clustering module in Scikit-Learn. The most frequently occurring “significant condition” was used to name each cluster. A supplementary file (tsne.ipynb) contains the Jupyter notebook used in these analyses.

### Bioinformatic Analyses

All statistical analyses were carried out using either the Python Scipy or Numpy libraries (155, 156). To identify the number of aminotransferases, the HMMER version 3.3.2 (http://hmmer.org/) was used to scan the *P. putida* KT2440 genome against a profile database file consisting out of the Pfam HMM files of PF00155, PF00202, PF01063 and PF00266. The E-value cutoff was set to 1e-20, and significant hits annotated as transcriptional regulators were excluded from the final dataset. HMMER was also used to identify possible genes of the γ- glutamyl cycle, with the Pfam HMM files of PF00120, PF01266, PF00171, and PF07722 corresponding to the glutamyl-polyamine synthetase, the glutamyl-polyamine oxidase, the aldehyde dehydrogenase, and the amide hydrolase.

The databases MiST 3.0 and TransportDB 2.0 were used to extract transcription factors and transport associated proteins from the BarSeq dataset (17, 18). EggNOGmapper was used to generate the Clusters of Orthologous Groups (COGs) for *P. putida* KT2440 (21, 157). Additionally, manual analysis of the data and proposals of metabolic pathways relied heavily on BioCyc and PaperBlast (158, 159)

## ACKNOWLEDGEMENTS

We would like to thank Morgan Price and Dr. Megan Garber for their assistance in analyzing BarSeq data, Alberto Nava for his advice on manifold learning methods, Dr. Elias Englund for his help on protein purification and Dr. Namil Lee for his advice on gene annotations. Mitchell Thompson is a Simons Foundation Awardee of the Life Sciences Research Foundation. The laboratory of LMB is partially funded by the Deutsche Forschungsgemeinschaft (DFG, German Research Foundation) under Germany’s Excellence Strategy within the Cluster of Excellence FSC 2186 ‘The Fuel Science Center’. This work was part of the DOE Joint BioEnergy Institute (https://www.jbei.org) supported by the U. S. Department of Energy, Office of Science, Office of Biological and Environmental Research, supported by the U.S. Department of Energy, Energy Efficiency and Renewable Energy, Bioenergy Technologies Office, through contract DE-AC02- 05CH11231 between Lawrence Berkeley National Laboratory and the U.S. Department of Energy. The views and opinions of the authors expressed herein do not necessarily state or reflect those of the United States Government or any agency thereof. Neither the United States Government nor any agency thereof, nor any of their employees, makes any warranty, expressed or implied, or assumes any legal liability or responsibility for the accuracy, completeness, or usefulness of any information, apparatus, product, or process disclosed, or represents that its use would not infringe privately owned rights. The United States Government retains and the publisher, by accepting the article for publication, acknowledges that the United States Government retains a nonexclusive, paid-up, irrevocable, worldwide license to publish or reproduce the published form of this manuscript, or allow others to do so, for United States Government purposes. The Department of Energy will provide public access to these results of federally sponsored research in accordance with the DOE Public Access Plan (http://energy.gov/downloads/doe-public-access-plan).

## CONTRIBUTIONS

Conceptualization, M.S., A.N.P., M.R.I., M.G.T.; Methodology, M.S., A.N.P., M.R.I., M.G.T.; Investigation, M.S., A.N.P, M.R.I., E.E.K.B. M.G.T, R.K.; Writing – Original Draft, M.S., A.N.P., M.R.I.; Writing – Review and Editing, All authors.; Resources and supervision, M.G.T., L.M.B., A.M.D., P.M.S, J.D.K. M.S. and A.N.P. contributed equally to this work. Author order was determined by number of sesamoid bones.

## Bibliography

1. López-Farfán D, Reyes-Darias JA, Matilla MA, Krell T. 2019. Concentration Dependent Effect of Plant Root Exudates on the Chemosensory Systems of Pseudomonas putida KT2440. Front Microbiol 10:78.

2. Fernández M, Morel B, Corral-Lugo A, Krell T. 2016. Identification of a chemoreceptor that specifically mediates chemotaxis toward metabolizable purine derivatives. Mol Microbiol 99:34–42.

3. Neal AL, Ahmad S, Gordon-Weeks R, Ton J. 2012. Benzoxazinoids in root exudates of maize attract Pseudomonas putida to the rhizosphere. PLoS ONE 7:e35498.

4. Neal AL, Ton J. 2013. Systemic defense priming by Pseudomonas putida KT2440 in maize depends on benzoxazinoid exudation from the roots. Plant Signal Behav 8:e22655.

5. Hervás AB, Canosa I, Santero E. 2008. Transcriptome analysis of *Pseudomonas putida* in response to nitrogen availability. J Bacteriol 190:416–420.

6. Poblete-Castro I, Escapa IF, Jäger C, Puchalka J, Lam CMC, Schomburg D, Prieto MA, Martins dos Santos VAP. 2012. The metabolic response of P. putida KT2442 producing high levels of polyhydroxyalkanoate under single- and multiple-nutrient-limited growth: highlights from a multi-level omics approach. Microb Cell Fact 11:34.

7. Mozejko-Ciesielska J, Dabrowska D, Szalewska-Palasz A, Ciesielski S. 2017. Medium-chain-length polyhydroxyalkanoates synthesis by Pseudomonas putida KT2440 relA/spoT mutant: bioprocess characterization and transcriptome analysis. AMB Express 7:92.

8. Galman JL, Slabu I, Weise NJ, Iglesias C, Parmeggiani F, Lloyd RC, Turner NJ. 2017. Biocatalytic transamination with near-stoichiometric inexpensive amine donors mediated by bifunctional mono- and di- amine transaminases. Green Chem 19:361–366.

9. Han T, Kim GB, Lee SY. 2020. Glutaric acid production by systems metabolic engineering of an l-lysine- overproducing Corynebacterium glutamicum. Proc Natl Acad Sci USA 117:30328–30334.

10. Thompson MG, Pearson AN, Barajas JF, Cruz-Morales P, Sedaghatian N, Costello Z, Garber ME, Incha MR, Valencia LE, Baidoo EEK, Martin HG, Mukhopadhyay A, Keasling JD. 2020. Identification, Characterization, and Application of a Highly Sensitive Lactam Biosensor from Pseudomonas putida. ACS Synth Biol 9:53–62.

11. Banerjee D, Eng T, Lau AK, Sasaki Y, Wang B, Chen Y, Prahl J-P, Singan VR, Herbert RA, Liu Y, Tanjore D, Petzold CJ, Keasling JD, Mukhopadhyay A. 2020. Genome-scale metabolic rewiring improves titers rates and yields of the non-native product indigoidine at scale. Nat Commun 11:5385.

12. Nogales J, Mueller J, Gudmundsson S, Canalejo FJ, Duque E, Monk J, Feist AM, Ramos JL, Niu W, Palsson BO. 2020. High-quality genome-scale metabolic modelling of *Pseudomonas putida* highlights its broad metabolic capabilities. Environ Microbiol 22:255–269.

13. Thompson MG, Incha MR, Pearson AN, Schmidt M, Sharpless WA, Eiben CB, Cruz-Morales P, Blake- Hedges JM, Liu Y, Adams CA, Haushalter RW, Krishna RN, Lichtner P, Blank LM, Mukhopadhyay A, Deutschbauer AM, Shih PM, Keasling JD. 2020. Fatty acid and alcohol metabolism in Pseudomonas putida: functional analysis using random barcode transposon sequencing. Appl Environ Microbiol 86.

14. Incha MR, Thompson MG, Blake-Hedges JM, Liu Y, Pearson AN, Schmidt M, Gin JW, Petzold CJ, Deutschbauer AM, Keasling JD. 2020. Leveraging host metabolism for bisdemethoxycurcumin production in Pseudomonas putida. Metab Eng Commun 10:e00119.

15. Thompson MG, Blake-Hedges JM, Cruz-Morales P, Barajas JF, Curran SC, Eiben CB, Harris NC, Benites VT, Gin JW, Sharpless WA, Twigg FF, Skyrud W, Krishna RN, Pereira JH, Baidoo EEK, Petzold CJ, Adams PD, Arkin AP, Deutschbauer AM, Keasling JD. 2019. Massively parallel fitness profiling reveals multiple novel enzymes in Pseudomonas putida lysine metabolism. MBio 10.

16. Thompson MG, Valencia LE, Blake-Hedges JM, Cruz-Morales P, Velasquez AE, Pearson AN, Sermeno LN, Sharpless WA, Benites VT, Chen Y, Baidoo EEK, Petzold CJ, Deutschbauer AM, Keasling JD. 2019. Omics-driven identification and elimination of valerolactam catabolism in Pseudomonas putida KT2440 for increased product titer. Metab Eng Commun 9:e00098.

17. Gumerov VM, Ortega DR, Adebali O, Ulrich LE, Zhulin IB. 2020. MiST 3.0: an updated microbial signal transduction database with an emphasis on chemosensory systems. Nucleic Acids Res 48:D459–D464.

18. Elbourne LDH, Tetu SG, Hassan KA, Paulsen IT. 2017. TransportDB 2.0: a database for exploring membrane transporters in sequenced genomes from all domains of life. Nucleic Acids Res 45:D320–D324.

19. van der Maaten L, Hinton G. 2008. Visualizing Data using t-SNE. Journal of Machine Learning Research.

20. Price MN, Wetmore KM, Waters RJ, Callaghan M, Ray J, Liu H, Kuehl JV, Melnyk RA, Lamson JS, Suh Y, Carlson HK, Esquivel Z, Sadeeshkumar H, Chakraborty R, Zane GM, Rubin BE, Wall JD, Visel A, Bristow J, Blow MJ, Deutschbauer AM. 2018. Mutant phenotypes for thousands of bacterial genes of unknown function. Nature 557:503–509.

21. Huerta-Cepas J, Szklarczyk D, Heller D, Hernández-Plaza A, Forslund SK, Cook H, Mende DR, Letunic I, Rattei T, Jensen LJ, von Mering C, Bork P. 2019. eggNOG 5.0: a hierarchical, functionally and phylogenetically annotated orthology resource based on 5090 organisms and 2502 viruses. Nucleic Acids Res 47:D309–D314.

22. Sonawane AM, Röhm KH. 2004. A functional gltB gene is essential for utilization of acidic amino acids and expression of periplasmic glutaminase/asparaginase (PGA) by Pseudomonas putida KT2440. Mol Genet Genomics 271:33–39.

23. Eberl L, Ammendola A, Rothballer MH, Givskov M, Sternberg C, Kilstrup M, Schleifer KH, Molin S. 2000. Inactivation of gltB abolishes expression of the assimilatory nitrate reductase gene (nasB) in Pseudomonas putida KT2442. J Bacteriol 182:3368–3376.

24. Reitzer L. 2003. Nitrogen assimilation and global regulation in Escherichia coli. Annu Rev Microbiol 57:155–176.

25. Venturi V. 2003. Control of rpoS transcription in Escherichia coli and Pseudomonas: why so different? Mol Microbiol 49:1–9.

26. Lapouge K, Schubert M, Allain FH-T, Haas D. 2008. Gac/Rsm signal transduction pathway of gamma- proteobacteria: from RNA recognition to regulation of social behaviour. Mol Microbiol 67:241–253.

27. Martínez-Gil M, Ramos-González MI, Espinosa-Urgel M. 2014. Roles of cyclic Di-GMP and the Gac system in transcriptional control of the genes coding for the Pseudomonas putida adhesins LapA and LapF. J Bacteriol 196:1484–1495.

28. Bentley GJ, Narayanan N, Jha RK, Salvachúa D, Elmore JR, Peabody GL, Black BA, Ramirez K, De Capite A, Michener WE, Werner AZ, Klingeman DM, Schindel HS, Nelson R, Foust L, Guss AM, Dale T, Johnson CW, Beckham GT. 2020. Engineering glucose metabolism for enhanced muconic acid production in Pseudomonas putida KT2440. Metab Eng 59:64–75.

29. Eng T, Banerjee D, Lau AK, Bowden E, Herbert RA, Trinh J, Prahl J-P, Deutschbauer A, Tanjore D, Mukhopadhyay A. 2021. Engineering Pseudomonas putida for efficient aromatic conversion to bioproduct using high throughput screening in a bioreactor. Metab Eng 66:229–238.

30. 30. Demling P, Ankenbauer A, Klein B, Noack S, Tiso T, Takors R, Blank LM. 2021. Pseudomonas putida KT2440 endures temporary oxygen limitations. Biotechnol Bioeng https://doi.org/10.1002/bit.27938.

31. Merrick MJ, Edwards RA. 1995. Nitrogen control in bacteria. Microbiol Rev 59:604–622.

32. Pino C, Olmo-Mira F, Cabello P, Martínez-Luque M, Castillo F, Roldán MD, Moreno-Vivián C. 2006. The assimilatory nitrate reduction system of the phototrophic bacterium Rhodobacter capsulatus E1F1. Biochem Soc Trans 34:127–129.

33. Luque-Almagro VM, Gates AJ, Moreno-Vivián C, Ferguson SJ, Richardson DJ, Roldán MD. 2011. Bacterial nitrate assimilation: gene distribution and regulation. Biochem Soc Trans 39:1838–1843.

34. Caballero A, Esteve-Núñez A, Zylstra GJ, Ramos JL. 2005. Assimilation of nitrogen from nitrite and trinitrotoluene in Pseudomonas putida JLR11. J Bacteriol 187:396–399.

35. Romeo A, Sonnleitner E, Sorger-Domenigg T, Nakano M, Eisenhaber B, Bläsi U. 2012. Transcriptional regulation of nitrate assimilation in Pseudomonas aeruginosa occurs via transcriptional antitermination within the nirBD-PA1779-cobA operon. Microbiology (Reading, Engl) 158:1543–1552.

36. Wang B, Rensing C, Pierson LS, Zhao H, Kennedy C. 2014. Translational coupling of nasST expression in Azotobacter vinelandii prevents overexpression of the nasT gene. FEMS Microbiol Lett 361:123–130.

37. Maeda S, Okamura M, Kobayashi M, Omata T. 1998. Nitrite-specific active transport system of the cyanobacterium Synechococcus sp. strain PCC 7942. J Bacteriol 180:6761–6763.

38. Gao S-H, Fan L, Yuan Z, Bond PL. 2015. The concentration-determined and population-specific antimicrobial effects of free nitrous acid on Pseudomonas aeruginosa PAO1. Appl Microbiol Biotechnol 99:2305–2312.

39. Kusumawardhani H, van Dijk D, Hosseini R, de Winde JH. 2020. Novel Toxin-Antitoxin Module SlvT-SlvA Regulates Megaplasmid Stability and Incites Solvent Tolerance in Pseudomonas putida S12. Appl Environ Microbiol 86.

40. Skjerning RB, Senissar M, Winther KS, Gerdes K, Brodersen DE. 2019. The RES domain toxins of RES- Xre toxin-antitoxin modules induce cell stasis by degrading NAD+. Mol Microbiol 111:221–236.

41. Fuhrer T, Fischer E, Sauer U. 2005. Experimental identification and quantification of glucose metabolism in seven bacterial species. J Bacteriol 187:1581–1590.

42. Nikel PI, Fuhrer T, Chavarría M, Sánchez-Pascuala A, Sauer U, de Lorenzo V. 2021. Reconfiguration of metabolic fluxes in Pseudomonas putida as a response to sub-lethal oxidative stress. ISME J 15:1751–1766.

43. del Castillo T, Ramos JL, Rodríguez-Herva JJ, Fuhrer T, Sauer U, Duque E. 2007. Convergent peripheral pathways catalyze initial glucose catabolism in Pseudomonas putida: genomic and flux analysis. J Bacteriol 189:5142–5152.

44. Roon RJ, Levenberg B. 1968. An adenosine triphosphate-dependent, avidin-sensitive enzymatic cleavage of urea in yeast and green algae. J Biol Chem 243:5213–5215.

45. Roon RJ, Levenberg B. 1972. Urea amidolyase. I. Properties of the enzyme from Candida utilis. J Biol Chem 247:4107–4113.

46. Mackay EM, Pateman JA. 1982. The regulation of urease activity in Aspergillus nidulans. Biochem Genet 20:763–776.

47. Kanamori T, Kanou N, Atomi H, Imanaka T. 2004. Enzymatic characterization of a prokaryotic urea carboxylase. J Bacteriol 186:2532–2539.

48. Nicholas DJD, Nason A. 1954. Molybdenum and nitrate reductase II. Molybdenum as a constituent of nitrate reductase. Journal of Biological Chemistry.

49. Scott E, Peter F, Sanders J. 2007. Biomass in the manufacture of industrial products--the use of proteins and amino acids. Appl Microbiol Biotechnol 75:751–762.

50. Hermann T. 2003. Industrial production of amino acids by coryneform bacteria. J Biotechnol 104:155–172.

51. Nijkamp K, Westerhof RGM, Ballerstedt H, de Bont JAM, Wery J. 2007. Optimization of the solvent- tolerant Pseudomonas putida S12 as host for the production of p-coumarate from glucose. Appl Microbiol Biotechnol 74:617–624.

52. Craig JW, Chang F-Y, Kim JH, Obiajulu SC, Brady SF. 2010. Expanding small-molecule functional metagenomics through parallel screening of broad-host-range cosmid environmental DNA libraries in diverse proteobacteria. Appl Environ Microbiol 76:1633–1641.

53. Verhoef S, Ballerstedt H, Volkers RJM, de Winde JH, Ruijssenaars HJ. 2010. Comparative transcriptomics and proteomics of p-hydroxybenzoate producing Pseudomonas putida S12: novel responses and implications for strain improvement. Appl Microbiol Biotechnol 87:679–690.

54. 54. Xing X, Jiang P. July 2011. Recombinant bacteria for producing deoxyviolacein and uses thereof. US20110183384A1. United States.

55. Schmitz S, Nies S, Wierckx N, Blank LM, Rosenbaum MA. 2015. Engineering mediator-based electroactivity in the obligate aerobic bacterium Pseudomonas putida KT2440. Front Microbiol 6:284.

56. Molina-Santiago C, Cordero BF, Daddaoua A, Udaondo Z, Manzano J, Valdivia M, Segura A, Ramos J-L, Duque E. 2016. Pseudomonas putida as a platform for the synthesis of aromatic compounds. Microbiology (Reading, Engl) 162:1535–1543.

57. Aboulmagd E, Voss I, Oppermann-Sanio FB, Steinbüchel A. 2001. Heterologous expression of cyanophycin synthetase and cyanophycin synthesis in the industrial relevant bacteria Corynebacterium glutamicum and Ralstonia eutropha and in Pseudomonas putida. Biomacromolecules 2:1338–1342.

58. Voss I, Diniz SC, Aboulmagd E, Steinbüchel A. 2004. Identification of the Anabaena sp. strain PCC7120 cyanophycin synthetase as suitable enzyme for production of cyanophycin in gram-negative bacteria like Pseudomonas putida and Ralstonia eutropha. Biomacromolecules 5:1588–1595.

59. Wiefel L, Bröker A, Steinbüchel A. 2011. Synthesis of a citrulline-rich cyanophycin by use of Pseudomonas putida ATCC 4359. Appl Microbiol Biotechnol 90:1755–1762.

60. Loeschcke A, Thies S. 2015. Pseudomonas putida-a versatile host for the production of natural products. Appl Microbiol Biotechnol 99:6197–6214.

61. Radkov AD, Moe LA. 2013. Amino acid racemization in Pseudomonas putida KT2440. J Bacteriol 195:5016–5024.

62. Mercenier A, Simon JP, Haas D, Stalon V. 1980. Catabolism of L-arginine by Pseudomonas aeruginosa. J Gen Microbiol 116:381–389.

63. Stalon V, Vander Wauven C, Momin P, Legrain C. 1987. Catabolism of arginine, citrulline and ornithine by Pseudomonas and related bacteria. J Gen Microbiol 133:2487–2495.

64. Li C, Lu C-D. 2009. Arginine racemization by coupled catabolic and anabolic dehydrogenases. Proc Natl Acad Sci USA 106:906–911.

65. Patil MD, Rathod VP, Bihade UR, Banerjee UC. 2019. Purification and characterization of arginine deiminase from Pseudomonas putida: Structural insights of the differential affinities of l-arginine analogues. J Biosci Bioeng 127:129–137.

66. Vander Wauven C, Jann A, Haas D, Leisinger T, Stalon V. 1988. N2-succinylornithine in ornithine catabolism of Pseudomonas aeruginosa. Arch Microbiol 150:400–404.

67. Tricot C, Vander Wauven C, Wattiez R, Falmagne P, Stalon V. 1994. Purification and properties of a succinyltransferase from Pseudomonas aeruginosa specific for both arginine and ornithine. Eur J Biochem 224:853–861.

68. Radkov AD, Moe LA. 2018. A Broad Spectrum Racemase in Pseudomonas putida KT2440 Plays a Key Role in Amino Acid Catabolism. Front Microbiol 9:1343.

69. Chang YF, Adams E. 1971. Induction of separate catabolic pathways for L- and D-lysine in Pseudomonas putida. Biochem Biophys Res Commun 45:570–577.

70. Espinosa-Urgel M, Ramos JL. 2001. Expression of a Pseudomonas putida aminotransferase involved in lysine catabolism is induced in the rhizosphere. Appl Environ Microbiol 67:5219–5224.

71. Revelles O, Espinosa-Urgel M, Molin S, Ramos JL. 2004. The davDT operon of Pseudomonas putida, involved in lysine catabolism, is induced in response to the pathway intermediate delta-aminovaleric acid. J Bacteriol 186:3439–3446.

72. Revelles O, Espinosa-Urgel M, Fuhrer T, Sauer U, Ramos JL. 2005. Multiple and interconnected pathways for L-lysine catabolism in Pseudomonas putida KT2440. J Bacteriol 187:7500–7510.

73. Revelles O, Wittich R-M, Ramos JL. 2007. Identification of the initial steps in D-lysine catabolism in Pseudomonas putida. J Bacteriol 189:2787–2792.

74. Lambert MP, Neuhaus FC. 1972. Factors affecting the level of alanine racemase in Escherichia coli. J Bacteriol 109:1156–1161.

75. Radkov AD, Moe LA. 2014. Bacterial synthesis of D-amino acids. Appl Microbiol Biotechnol 98:5363– 5374.

76. Duque E, Daddaoua A, Cordero BF, De la Torre J, Antonia Molina-Henares M, Ramos J-L. 2017. Identification and elucidation of in vivo function of two alanine racemases from Pseudomonas putida KT2440. Environ Microbiol Rep 9:581–588.

77. Boulette ML, Baynham PJ, Jorth PA, Kukavica-Ibrulj I, Longoria A, Barrera K, Levesque RC, Whiteley M. 2009. Characterization of alanine catabolism in Pseudomonas aeruginosa and its importance for proliferation in vivo. J Bacteriol 191:6329–6334.

78. Zhi J, Mathew E, Freundlich M. 1999. Lrp binds to two regions in the dadAX promoter region of Escherichia coli to repress and activate transcription directly. Mol Microbiol 32:29–40.

79. Schwanemann T, Otto M, Wierckx N, Wynands B. 2020. Pseudomonas as versatile aromatics cell factory. Biotechnol J 15:e1900569.

80. Whitaker RJ, Gaines CG, Jensen RA. 1982. A multispecific quintet of aromatic aminotransferases that overlap different biochemical pathways in Pseudomonas aeruginosa. J Biol Chem 257:13550–13556.

81. Molina-Henares MA, García-Salamanca A, Molina-Henares AJ, de la Torre J, Herrera MC, Ramos JL, Duque E. 2009. Functional analysis of aromatic biosynthetic pathways in Pseudomonas putida KT2440. Microb Biotechnol 2:91–100.

82. Pittard J, Yang J. 2008. Biosynthesis of the aromatic amino acids. Ecosal Plus 3.

83. Nogales J, Palsson BØ, Thiele I. 2008. A genome-scale metabolic reconstruction of Pseudomonas putida KT2440: iJN746 as a cell factory. BMC Syst Biol 2:79.

84. Leprince A, de Lorenzo V, Völler P, van Passel MWJ, Martins dos Santos VAP. 2012. Random and cyclical deletion of large DNA segments in the genome of Pseudomonas putida. Environ Microbiol 14:1444–1453.

85. Gadilohar BL, Shankarling GS. 2017. Choline based ionic liquids and their applications in organic transformation. J Mol Liq 227:234–261.

86. Wood JM. 2011. Bacterial osmoregulation: a paradigm for the study of cellular homeostasis. Annu Rev Microbiol 65:215–238.

87. Wargo MJ. 2013. Homeostasis and catabolism of choline and glycine betaine: lessons from Pseudomonas aeruginosa. Appl Environ Microbiol 79:2112–2120.

88. Geiger O, López-Lara IM, Sohlenkamp C. 2013. Phosphatidylcholine biosynthesis and function in bacteria. Biochim Biophys Acta 1831:503–513.

89. Belda E, van Heck RGA, José Lopez-Sanchez M, Cruveiller S, Barbe V, Fraser C, Klenk H-P, Petersen J, Morgat A, Nikel PI, Vallenet D, Rouy Z, Sekowska A, Martins Dos Santos VAP, de Lorenzo V, Danchin A, Médigue C. 2016. The revisited genome of Pseudomonas putida KT2440 enlightens its value as a robust metabolic chassis. Environ Microbiol 18:3403–3424.

90. Leys D, Basran J, Scrutton NS. 2003. Channelling and formation of “active” formaldehyde in dimethylglycine oxidase. EMBO J 22:4038–4048.

91. Wargo MJ, Hogan DA. 2009. Identification of genes required for Pseudomonas aeruginosa carnitine catabolism. Microbiology (Reading, Engl) 155:2411–2419.

92. Bastard K, Smith AAT, Vergne-Vaxelaire C, Perret A, Zaparucha A, De Melo-Minardi R, Mariage A, Boutard M, Debard A, Lechaplais C, Pelle C, Pellouin V, Perchat N, Petit J-L, Kreimeyer A, Medigue C, Weissenbach J, Artiguenave F, De Berardinis V, Vallenet D, Salanoubat M. 2014. Revealing the hidden functional diversity of an enzyme family. Nat Chem Biol 10:42–49.

93. 93. Meadows JA, Wargo MJ. 2018. Transcriptional Regulation of Carnitine Catabolism in Pseudomonas aeruginosa by CdhR. mSphere 3.

94. Lee S, Takahashi Y, Oura H, Suzuki-Minakuchi C, Okada K, Yamane H, Nomura N, Nojiri H. 2016. Effects of carbazole-degradative plasmid pCAR1 on biofilm morphology in Pseudomonas putida KT2440. Environ Microbiol Rep 8:261–271.

95. Nuccio ML, Russell BL, Nolte KD, Rathinasabapathi B, Gage DA, Hanson AD. 1998. The endogenous choline supply limits glycine betaine synthesis in transgenic tobacco expressing choline monooxygenase. Plant J 16:487–496.

96. Rontein D, Nishida I, Tashiro G, Yoshioka K, Wu WI, Voelker DR, Basset G, Hanson AD. 2001. Plants synthesize ethanolamine by direct decarboxylation of serine using a pyridoxal phosphate enzyme. J Biol Chem 276:35523–35529.

97. Lundgren BR, Sarwar Z, Pinto A, Ganley JG, Nomura CT. 2016. Ethanolamine Catabolism in *Pseudomonas aeruginosa* PAO1 Is Regulated by the Enhancer-Binding Protein EatR (PA4021) and the Alternative Sigma Factor RpoN. J Bacteriol 198:2318–2329.

98. Roof DM, Roth JR. 1988. Ethanolamine utilization in Salmonella typhimurium. J Bacteriol 170:3855–3863.

99. Chang GW, Chang JT. 1975. Evidence for the B12-dependent enzyme ethanolamine deaminase in Salmonella. Nature 254:150–151.

100. Scarlett FA, Turner JM. 1976. Microbial metabolism of amino alcohols. Ethanolamine catabolism mediated by coenzyme B12-dependent ethanolamine ammonia-lyase in Escherichia coli and Klebsiella aerogenes. J Gen Microbiol 95:173–176.

101. Cameron B, Briggs K, Pridmore S, Brefort G, Crouzet J. 1989. Cloning and analysis of genes involved in coenzyme B12 biosynthesis in Pseudomonas denitrificans. J Bacteriol 171:547–557.

102. 102. LaBauve AE, Wargo MJ. 2012. Growth and laboratory maintenance of Pseudomonas aeruginosa. Curr Protoc Microbiol Chapter 6:Unit 6E.1.

103. Heller K, Mann BJ, Kadner RJ. 1985. Cloning and expression of the gene for the vitamin B12 receptor protein in the outer membrane of Escherichia coli. J Bacteriol 161:896–903.

104. Cadieux N, Bradbeer C, Reeger-Schneider E, Köster W, Mohanty AK, Wiener MC, Kadner RJ. 2002. Identification of the periplasmic cobalamin-binding protein BtuF of Escherichia coli. J Bacteriol 184:706– 717.

105. Warren MJ, Raux E, Schubert HL, Escalante-Semerena JC. 2002. The biosynthesis of adenosylcobalamin (vitamin B12). Nat Prod Rep 19:390–412.

106. Martens JH, Barg H, Warren MJ, Jahn D. 2002. Microbial production of vitamin B12. Appl Microbiol Biotechnol 58:275–285.

107. Zayas CL, Escalante-Semerena JC. 2007. Reassessment of the late steps of coenzyme B12 synthesis in Salmonella enterica: evidence that dephosphorylation of adenosylcobalamin-5’-phosphate by the CobC phosphatase is the last step of the pathway. J Bacteriol 189:2210–2218.

108. Mori K, Bando R, Hieda N, Toraya T. 2004. Identification of a reactivating factor for adenosylcobalamin- dependent ethanolamine ammonia lyase. J Bacteriol 186:6845–6854.

109. Vogels GD, Van der Drift C. 1976. Degradation of purines and pyrimidines by microorganisms. Bacteriol Rev 40:403–468.

110. Tipton PA. 2006. Urate to allantoin, specifically (S)-allantoin. Nat Chem Biol 2:124–125.

111. Ramazzina I, Folli C, Secchi A, Berni R, Percudani R. 2006. Completing the uric acid degradation pathway through phylogenetic comparison of whole genomes. Nat Chem Biol 2:144–148.

112. Hassan KA, Liu Q, Henderson PJF, Paulsen IT. 2015. Homologs of the Acinetobacter baumannii AceI transporter represent a new family of bacterial multidrug efflux systems. MBio 6.

113. Juttukonda LJ, Green ER, Lonergan ZR, Heffern MC, Chang CJ, Skaar EP. 2019. Acinetobacter baumannii OxyR Regulates the Transcriptional Response to Hydrogen Peroxide. Infect Immun 87.

114. Zimmer DP, Soupene E, Lee HL, Wendisch VF, Khodursky AB, Peter BJ, Bender RA, Kustu S. 2000. Nitrogen regulatory protein C-controlled genes of *Escherichia coli*: scavenging as a defense against nitrogen limitation. Proc Natl Acad Sci USA 97:14674–14679.

115. Ware E. 1950. The chemistry of the hydantoins. Chem Rev 46:403–470.

116. Shorvon S. 1994. Status Epilepticus. Cambridge University Press, Cambridge.

117. 117. Drauz K, Grayson I, Kleemann A, Krimmer H-P, Leuchtenberger W, Weckbecker C. 2007. Amino AcidsUllmann’s encyclopedia of industrial chemistry. Wiley-VCH Verlag GmbH & Co. KGaA, Weinheim, Germany.

118. Ogawa J, Kaimura T, Yamada H, Shimizu S. 1994. Evaluation of pyrimidine- and hydantoin-degrading enzyme activities in aerobic bacteria. FEMS Microbiol Lett 122:55–60.

119. Solem E. 1974. The absolute configuration of beta-aminoisobutyric acid formed by degradation of thymine in man. Clin Chim Acta 53:183–190.

120. 120. Unable to find information for 8244409.

121. Zhang J, Barajas JF, Burdu M, Wang G, Baidoo EE, Keasling JD. 2017. Application of an Acyl-CoA Ligase from Streptomyces aizunensis for Lactam Biosynthesis. ACS Synth Biol 6:884–890.

122. Niehaus TD, Elbadawi-Sidhu M, de Crécy-Lagard V, Fiehn O, Hanson AD. 2017. Discovery of a widespread prokaryotic 5-oxoprolinase that was hiding in plain sight. J Biol Chem 292:16360–16367.

123. Miller DL, Rodwell VW. 1971. Metabolism of basic amino acids in Pseudomonas putida. Intermediates in L-arginine catabolism. J Biol Chem 246:5053–5058.

124. Bandounas L, Ballerstedt H, de Winde JH, Ruijssenaars HJ. 2011. Redundancy in putrescine catabolism in solvent tolerant Pseudomonas putida S12. J Biotechnol 154:1–10.

125. Chae TU, Ko Y-S, Hwang K-S, Lee SY. 2017. Metabolic engineering of Escherichia coli for the production of four-, five- and six-carbon lactams. Metab Eng 41:82–91.

126. Tiso T, Narancic T, Wei R, Pollet E, Beagan N, Schröder K, Honak A, Jiang M, Kenny ST, Wierckx N, Perrin R, Avérous L, Zimmermann W, O’Connor K, Blank LM. 2021. Towards bio-upcycling of polyethylene terephthalate. Metab Eng 66:167–178.

127. Tabor CW, Tabor H. 1985. Polyamines in microorganisms. Microbiol Rev 49:81–99.

128. Gaymans RJ, Van Utteren TEC, Van Den Berg JWA, Schuyer J. 1977. Preparation and some properties of nylon 46. J Polym Sci Polym Chem Ed 15:537–545.

129. Xu F, Zhong L, Xu Y, Zhang C, Zhang F, Zhang G. 2019. Highly efficient flame-retardant and soft cotton fabric prepared by a novel reactive flame retardant. Cellulose 26:4225–4240.

130. Roberts MR. 2007. Does GABA Act as a Signal in Plants?: Hints from Molecular Studies. Plant Signal Behav 2:408–409.

131. Bouché N, Fromm H. 2004. GABA in plants: just a metabolite? Trends Plant Sci 9:110–115.

132. 132. Trouillon J, Ragno M, Simon V, Attrée I, Elsen S. 2021. Transcription Inhibitors with XRE DNA-Binding and Cupin Signal-Sensing Domains Drive Metabolic Diversification in Pseudomonas. mSystems 6.

133. Reyes-Darias JA, García V, Rico-Jiménez M, Corral-Lugo A, Lesouhaitier O, Juárez-Hernández D, Yang Y, Bi S, Feuilloley M, Muñoz-Rojas J, Sourjik V, Krell T. 2015. Specific gamma-aminobutyrate chemotaxis in pseudomonads with different lifestyle. Mol Microbiol 97:488–501.

134. Kinnersley AM, Turano FJ. 2000. Gamma Aminobutyric Acid (GABA) and Plant Responses to Stress. CRC Crit Rev Plant Sci 19:479–509.

135. Bartsch K, von Johnn-Marteville A, Schulz A. 1990. Molecular analysis of two genes of the Escherichia coli gab cluster: nucleotide sequence of the glutamate:succinic semialdehyde transaminase gene (gabT) and characterization of the succinic semialdehyde dehydrogenase gene (gabD). J Bacteriol 172:7035–7042.

136. Weber N, Hatsch A, Labagnere L, Heider H. 2017. Production of (S)-2-aminobutyric acid and (S)-2- aminobutanol in Saccharomyces cerevisiae. Microb Cell Fact 16:51.

137. Zhang K, Li H, Cho KM, Liao JC. 2010. Expanding metabolism for total biosynthesis of the nonnatural amino acid L-homoalanine. Proc Natl Acad Sci USA 107:6234–6239.

138. Soga T, Baran R, Suematsu M, Ueno Y, Ikeda S, Sakurakawa T, Kakazu Y, Ishikawa T, Robert M, Nishioka T, Tomita M. 2006. Differential metabolomics reveals ophthalmic acid as an oxidative stress biomarker indicating hepatic glutathione consumption. J Biol Chem 281:16768–16776.

139. Ewering C, Heuser F, Benölken JK, Brämer CO, Steinbüchel A. 2006. Metabolic engineering of strains of Ralstonia eutropha and Pseudomonas putida for biotechnological production of 2-methylcitric acid. Metab Eng 8:587–602.

140. Cohen YR. 2002. β-Aminobutyric Acid-Induced Resistance Against Plant Pathogens. Plant Dis 86:448– 457.

141. Jumper J, Evans R, Pritzel A, Green T, Figurnov M, Ronneberger O, Tunyasuvunakool K, Bates R, Žídek A, Potapenko A, Bridgland A, Meyer C, Kohl SAA, Ballard AJ, Cowie A, Romera-Paredes B, Nikolov S, Jain R, Adler J, Back T, Hassabis D. 2021. Highly accurate protein structure prediction with AlphaFold. Nature 596:583–589.

142. Baek M, DiMaio F, Anishchenko I, Dauparas J, Ovchinnikov S, Lee GR, Wang J, Cong Q, Kinch LN, Schaeffer RD, Millán C, Park H, Adams C, Glassman CR, DeGiovanni A, Pereira JH, Rodrigues AV, van Dijk AA, Ebrecht AC, Opperman DJ, Baker D. 2021. Accurate prediction of protein structures and interactions using a three-track neural network. Science 373:871–876.

143. Ham TS, Dmytriv Z, Plahar H, Chen J, Hillson NJ, Keasling JD. 2012. Design, implementation and practice of JBEI-ICE: an open source biological part registry platform and tools. Nucleic Acids Res 40:e141.

144. 144. Chen J, Densmore D, Ham TS, Keasling JD, Hillson NJ. 2012. DeviceEditor visual biological CAD canvas. J Biol Eng 6:1.

145. Hillson NJ, Rosengarten RD, Keasling JD. 2012. j5 DNA assembly design automation software. ACS Synth Biol 1:14–21.

146. Gibson DG, Young L, Chuang R-Y, Venter JC, Hutchison CA, Smith HO. 2009. Enzymatic assembly of DNA molecules up to several hundred kilobases. Nat Methods 6:343–345.

147. Shanks RMQ, Kadouri DE, MacEachran DP, O’Toole GA. 2009. New yeast recombineering tools for bacteria. Plasmid 62:88–97.

148. Rosenberg AH, Lade BN, Chui DS, Lin SW, Dunn JJ, Studier FW. 1987. Vectors for selective expression of cloned DNAs by T7 RNA polymerase. Gene 56:125–135.

149. Bi C, Su P, Müller J, Yeh Y-C, Chhabra SR, Beller HR, Singer SW, Hillson NJ. 2013. Development of a broad-host synthetic biology toolbox for Ralstonia eutropha and its application to engineering hydrocarbon biofuel production. Microb Cell Fact 12:107.

150. Lee TS, Krupa RA, Zhang F, Hajimorad M, Holtz WJ, Prasad N, Lee SK, Keasling JD. 2011. BglBrick vectors and datasheets: A synthetic biology platform for gene expression. J Biol Eng 5:12.

151. Wetmore KM, Price MN, Waters RJ, Lamson JS, He J, Hoover CA, Blow MJ, Bristow J, Butland G, Arkin AP, Deutschbauer A. 2015. Rapid quantification of mutant fitness in diverse bacteria by sequencing randomly bar-coded transposons. MBio 6:e00306–15.

152. 152. Unable to find information for 912504.

153. Eudes A, Berthomieu R, Hao Z, Zhao N, Benites VT, Baidoo EEK, Loqué D. 2018. Production of muconic acid in plants. Metab Eng 46:13–19.

154. Pedregosa F, Varoquaux G, Gramfort A, Michel V, Thirion B, Grisel O, Blondel M, Prettenhofer P, Weiss R, Dubourg V, Vanderplas J, Passos A, Cournapeau D, Brucher M, Perrot M, Duchesnay É. 2011. Scikit- learn: Machine Learning in Python. Journal of Machine Learning Research.

155. Virtanen P, Gommers R, Oliphant TE, Haberland M, Reddy T, Cournapeau D, Burovski E, Peterson P, Weckesser W, Bright J, van der Walt SJ, Brett M, Wilson J, Millman KJ, Mayorov N, Nelson ARJ, Jones E, Kern R, Larson E, Carey CJ, SciPy 1.0 Contributors. 2020. SciPy 1.0: fundamental algorithms for scientific computing in Python. Nat Methods 17:261–272.

156. Harris CR, Millman KJ, van der Walt SJ, Gommers R, Virtanen P, Cournapeau D, Wieser E, Taylor J, Berg S, Smith NJ, Kern R, Picus M, Hoyer S, van Kerkwijk MH, Brett M, Haldane A, Del Río JF, Wiebe M, Peterson P, Gérard-Marchant P, Oliphant TE. 2020. Array programming with NumPy. Nature 585:357–362.

157. 157. Cantalapiedra CP, Hernández-Plaza A, Letunic I, Bork P, Huerta-Cepas J. 2021. eggNOG-mapper v2: Functional Annotation, Orthology Assignments, and Domain Prediction at the Metagenomic Scale. Mol Biol Evol https://doi.org/10.1093/molbev/msab293.

158. Karp PD, Billington R, Caspi R, Fulcher CA, Latendresse M, Kothari A, Keseler IM, Krummenacker M, Midford PE, Ong Q, Ong WK, Paley SM, Subhraveti P. 2019. The BioCyc collection of microbial genomes and metabolic pathways. Brief Bioinformatics 20:1085–1093.

159. Price MN, Arkin AP. 2017. PaperBLAST: Text Mining Papers for Information about Homologs. mSystems 2.

