## Supplemental Figures for "Nitrogen metabolism in *Pseudomonas putida*: functional analysis using random barcode transposon sequencing"

### SUPPLEMENTAL MATERIAL

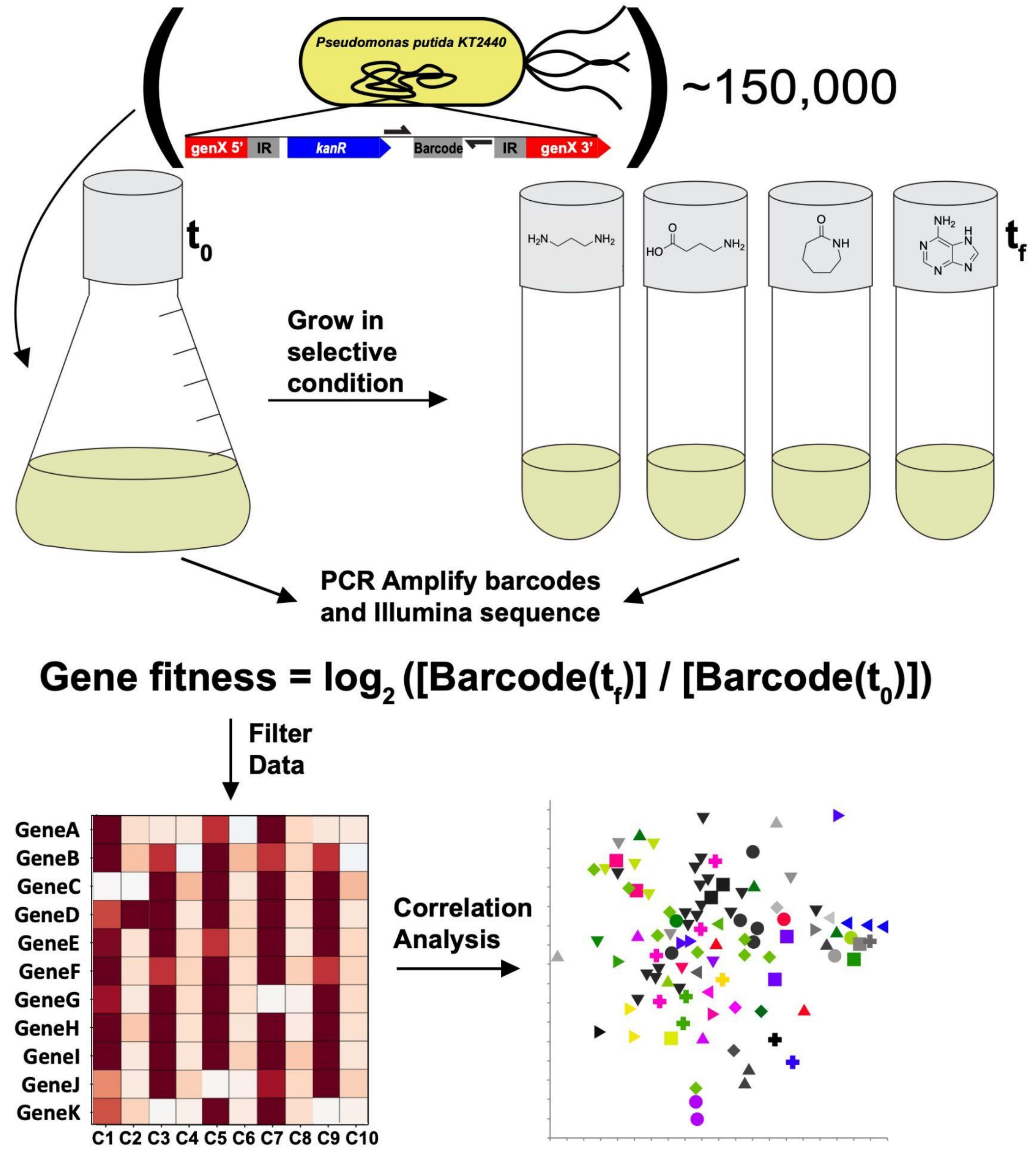


Figure S1: Overview of the barcode abundance sequencing (BarSeq) and data processing workflow.

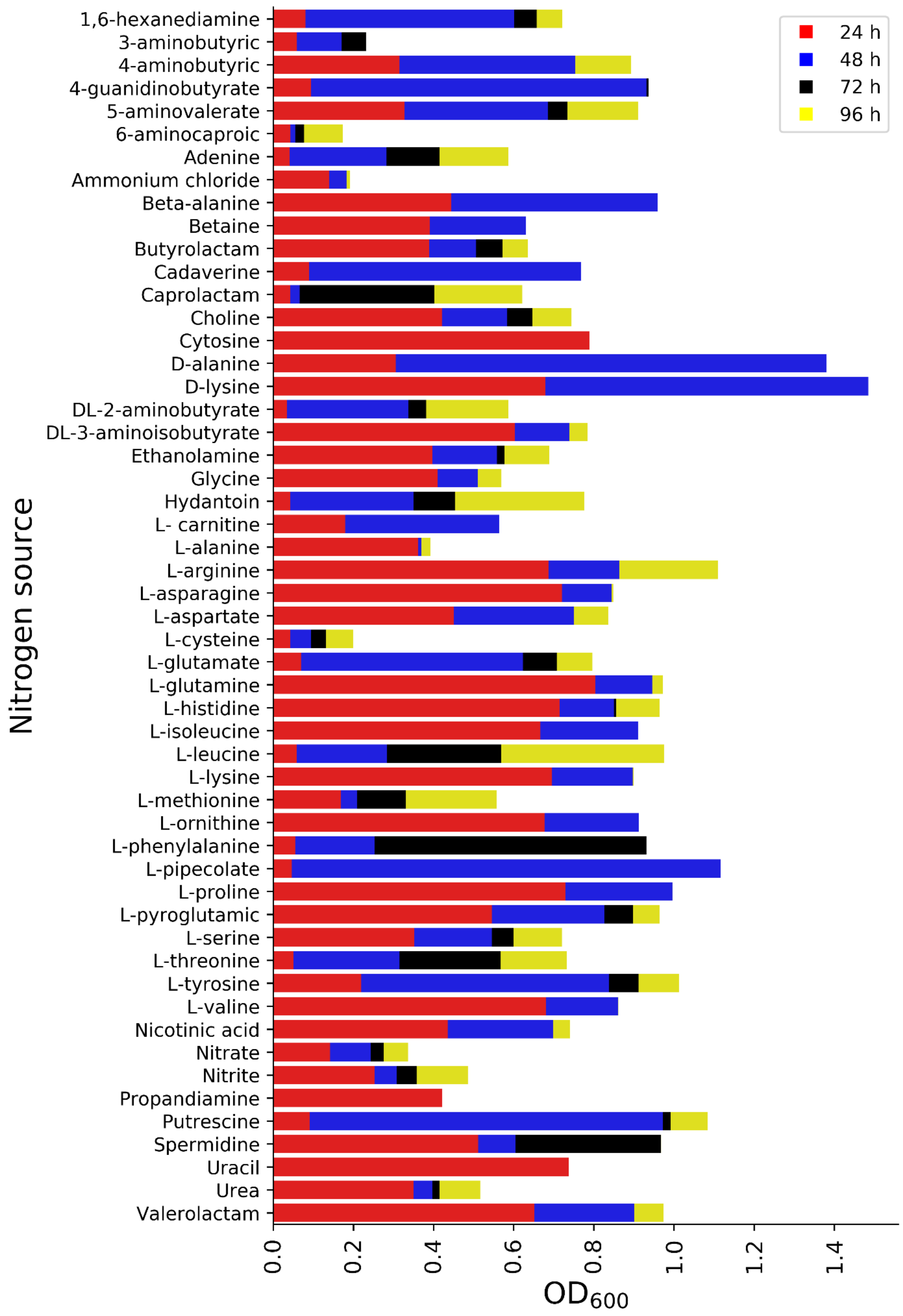


Figure S2: Plate-based growth assay showing the maximal OD for all the tested nitrogen sources. Cells were grown as triplicates in a 24-deep well plate and OD was taken after 24, 48, 72 and 96 hours.

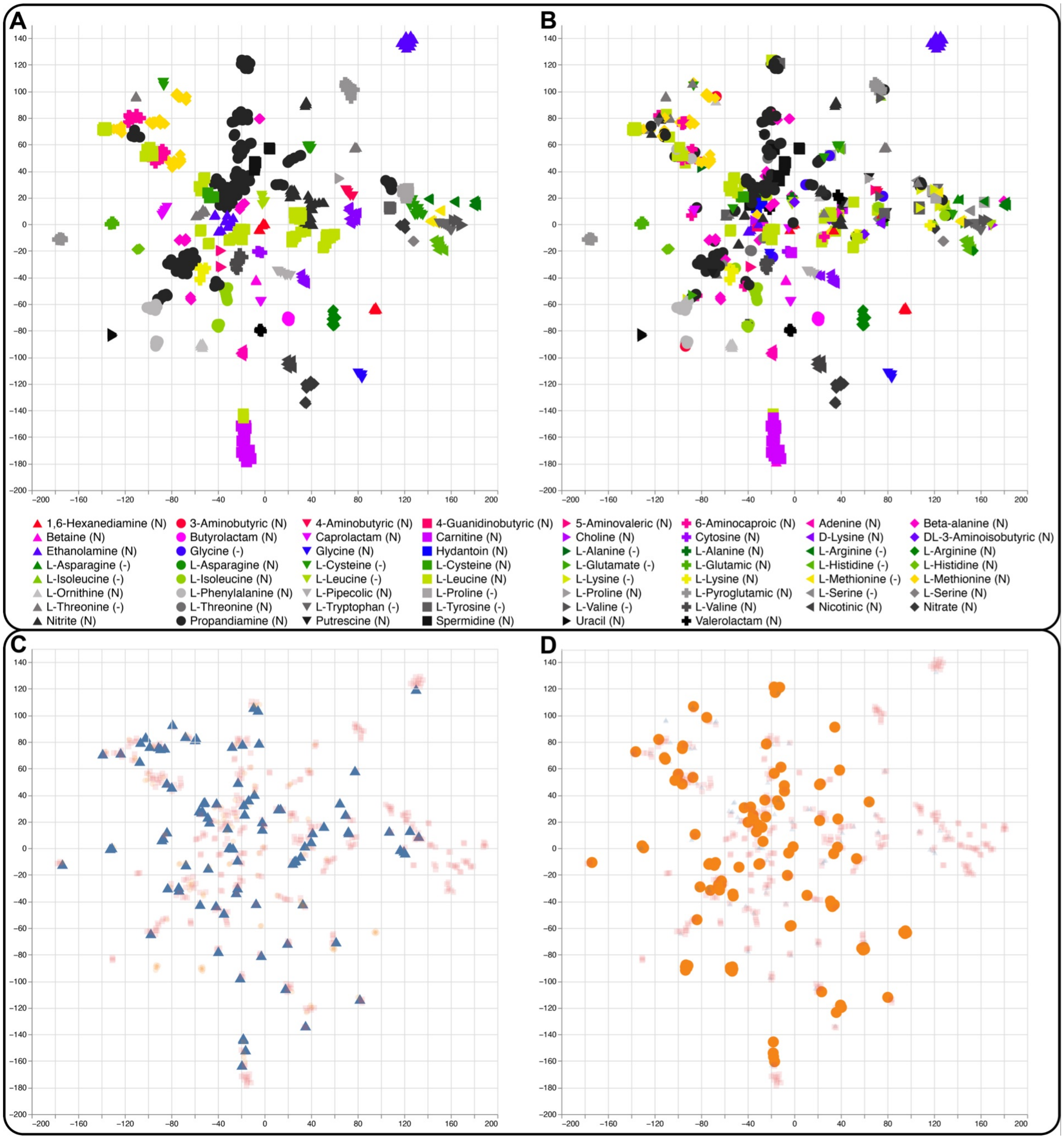


Figure S3: A) t-SNE visualization of the nitrogen source and amino acid dropout BarSeq assays. Individual points represent genes with significant fitness changes in the dataset. The x and y axes are non-dimensional and distances between points within clusters are representative of the Euclidean distance between the genes in the dataset. Clusters were defined using Ward clustering of the t-SNE results. B) Depiction of t-SNE results without Ward cluster definitions. C) Depiction of cluster locations for transcriptional regulators (blue triangles identified from MiSTdb

3.0. D) Depiction of cluster locations for genes predicted to be involved in transport reactions (orange circles) from TransportDB 2.0. Interactive versions of these plots are available in the supplementary materials (FigureS1.html). (Figure I2): An interactive version of the t-SNE analysis shown in Figure 1B with an extra panel (top right) excluding Ward clustering. Clicking in the top left chart highlights a cluster and generates a list of cluster components in the bottom right. Clicking in the top right panel highlights a specific gene. Clicking a point in the bottom left panel will open a comparison of the clustered genes in the Fitness Browser (https://fit.genomics.lbl.gov). (Figure I3): In this interactive figure, the shapes and colors are determined by the COG identifier for each individual gene. Clicking in the legend highlights all genes that are associated with a particular COG, and clicking in the scatterplot generates a list co-clustered genes

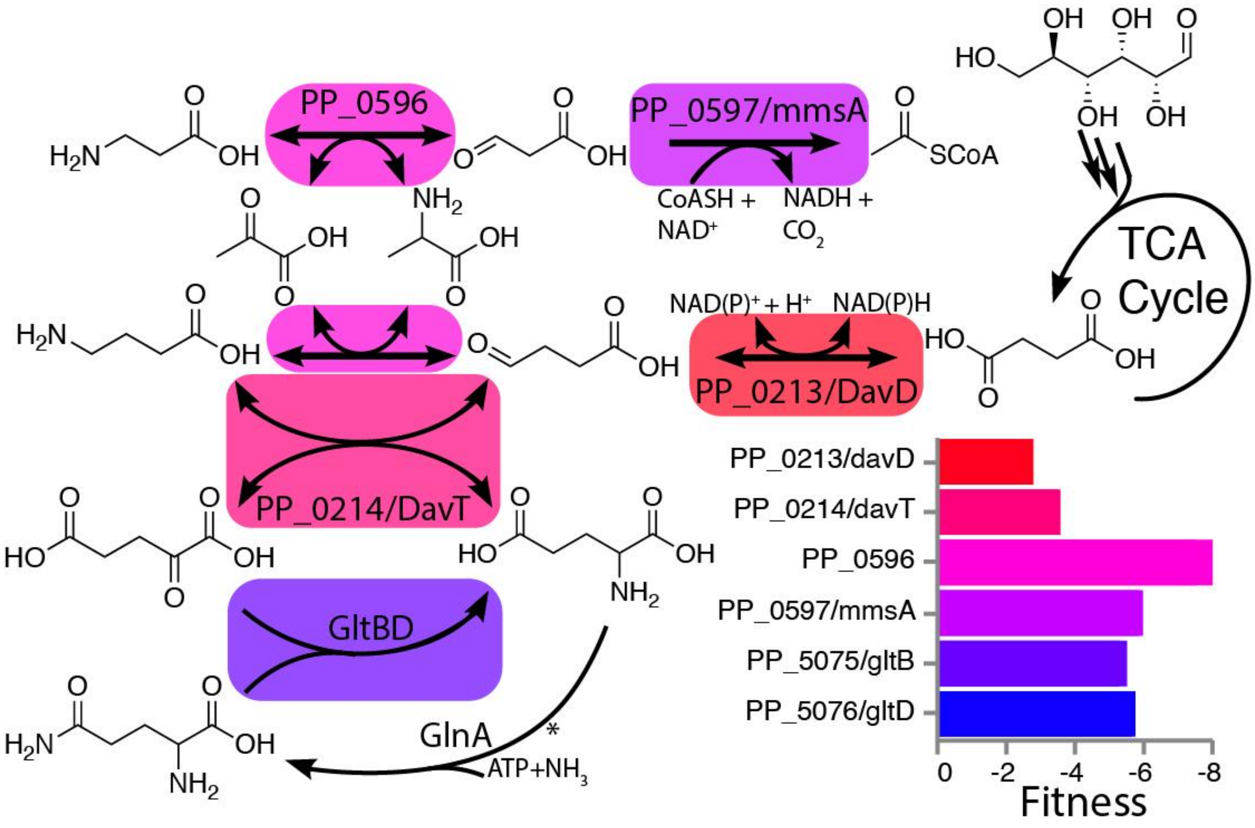


Figure S4: Proposed pathway for beta-alanine metabolism in *P. putida* that satisfies the requirements for *davD*, *davT*, *gltBD*, *mmsA*, and PP_0596. Inset bar chart shows the fitness values for the genes depicted in the pathway. While oxidation of 3-oxopropionate (malonyl-semialdehyde) to malonate is also hypothetically possible via DavD, *P. putida* cannot use malonate as a sole carbon source indicating it may be a dead-end metabolite (unpublished data). Deeper biochemical or genetic analyses of this pathway are necessary to validate this mechanism of alanine and glutamate balancing during beta-alanine metabolism.

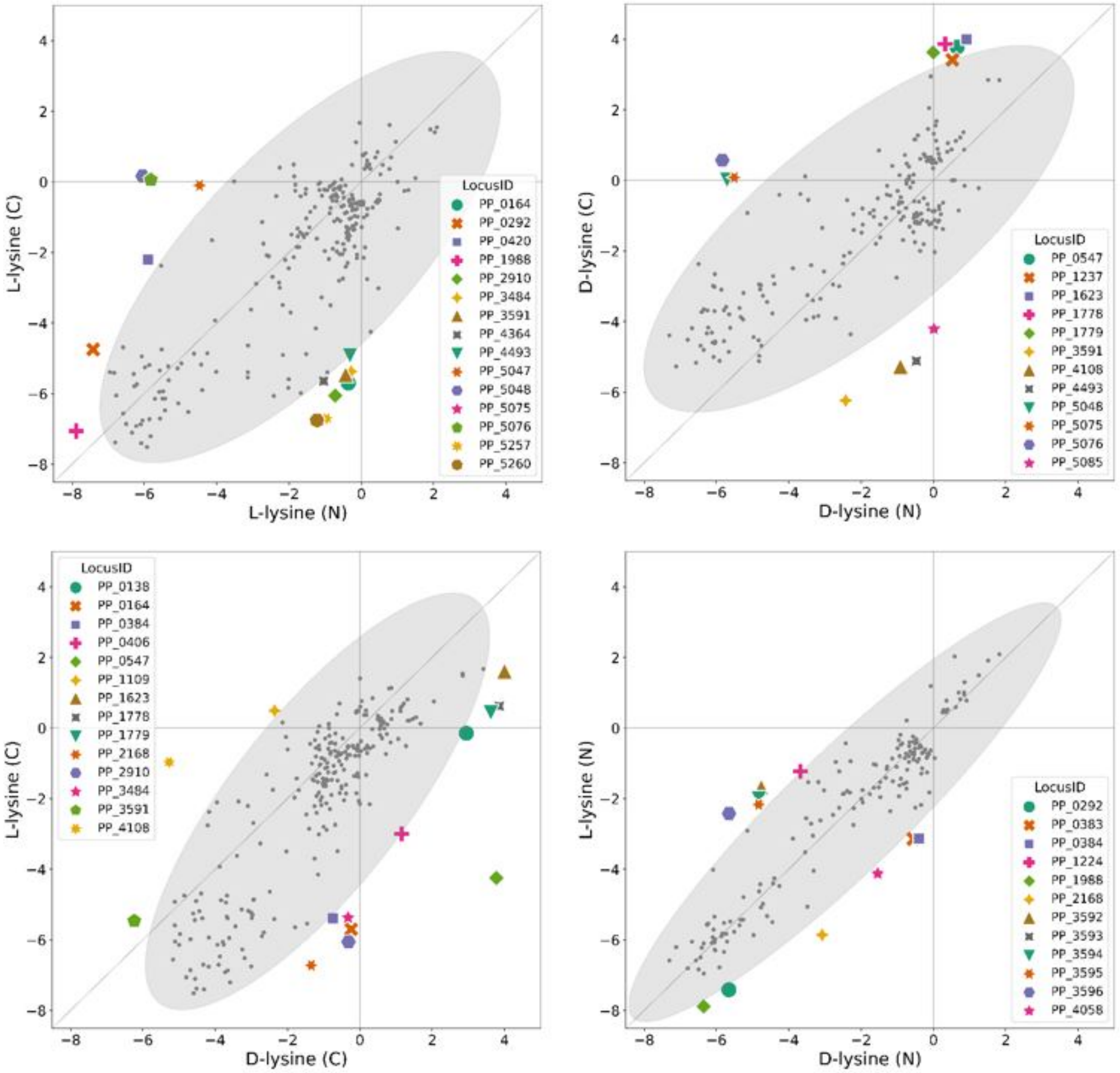


Figure S5: Mahalanobis distance and outlier detection for the comparison of all the tested lysine conditions. Prior to analysis, fitness scores were filtered by t-values (t > |4|). Outliers are defined as data points outside of the 95% confidence interval of the distribution, marked by the grey ellipsoid.

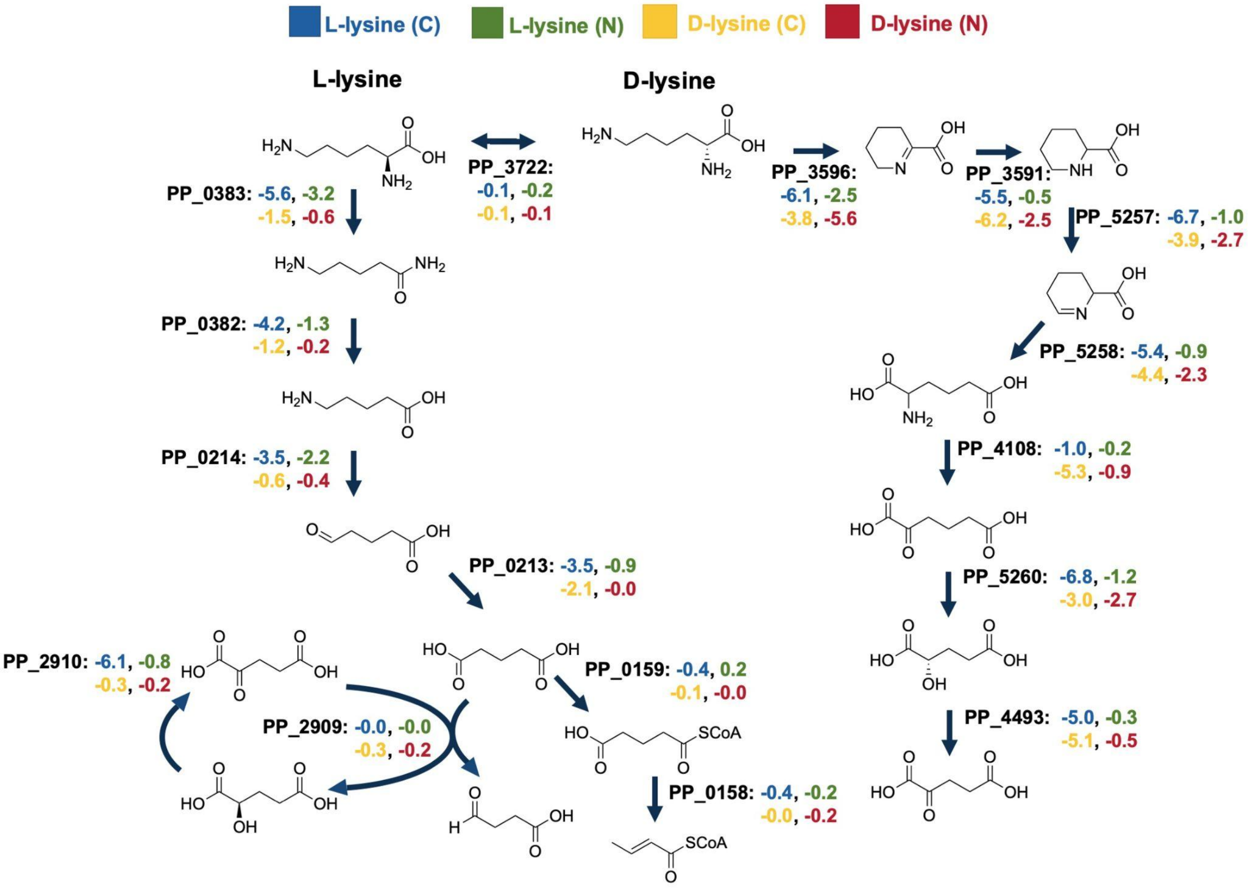


Figure S6: D and L-lysine degradation pathways and fitness values. Color indicates whether fitness value is from L-lysine carbon source (blue), L-lysine nitrogen source (green), D-lysine carbon source (yellow), or D-lysine nitrogen source (red) experiments.

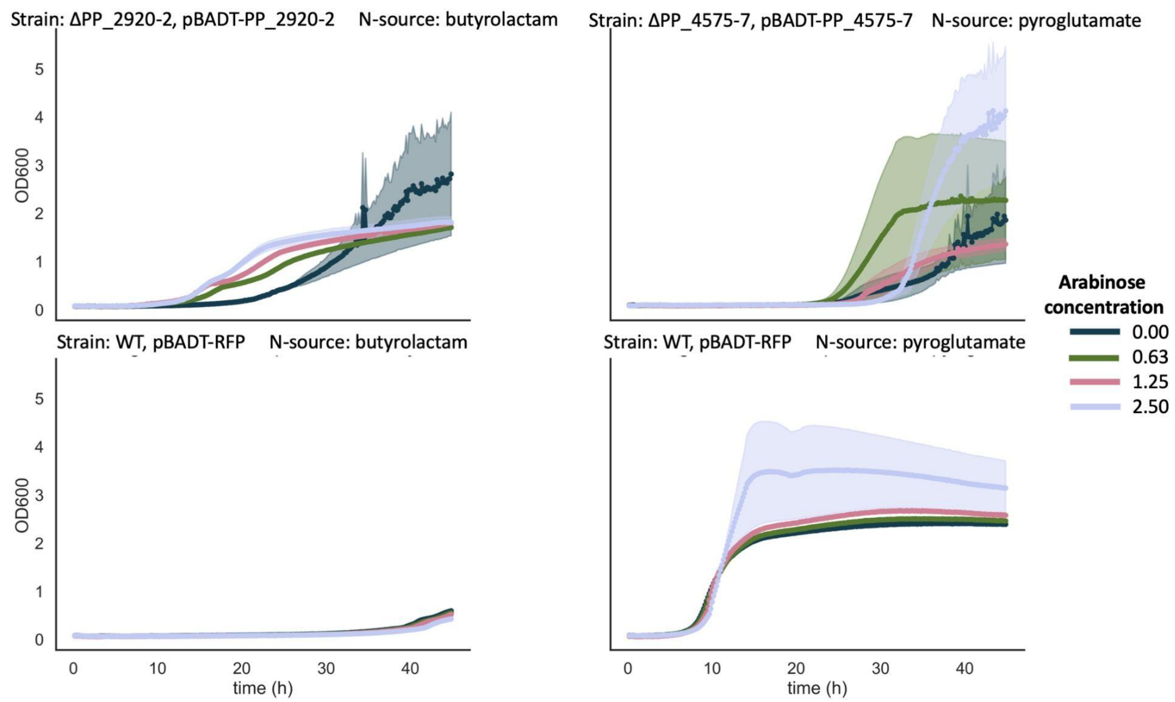


Figure S7: Complementation assays of lactamase knockouts. Knockout strains were complemented with arabinose induced expression of the lactamase from a pBADT kanamycin resistant backbone. Wild type controls carried RFP expressed from the same backbone. It appeared that leaky expression of the lactamase was sufficient to restore growth in the deletion strains.

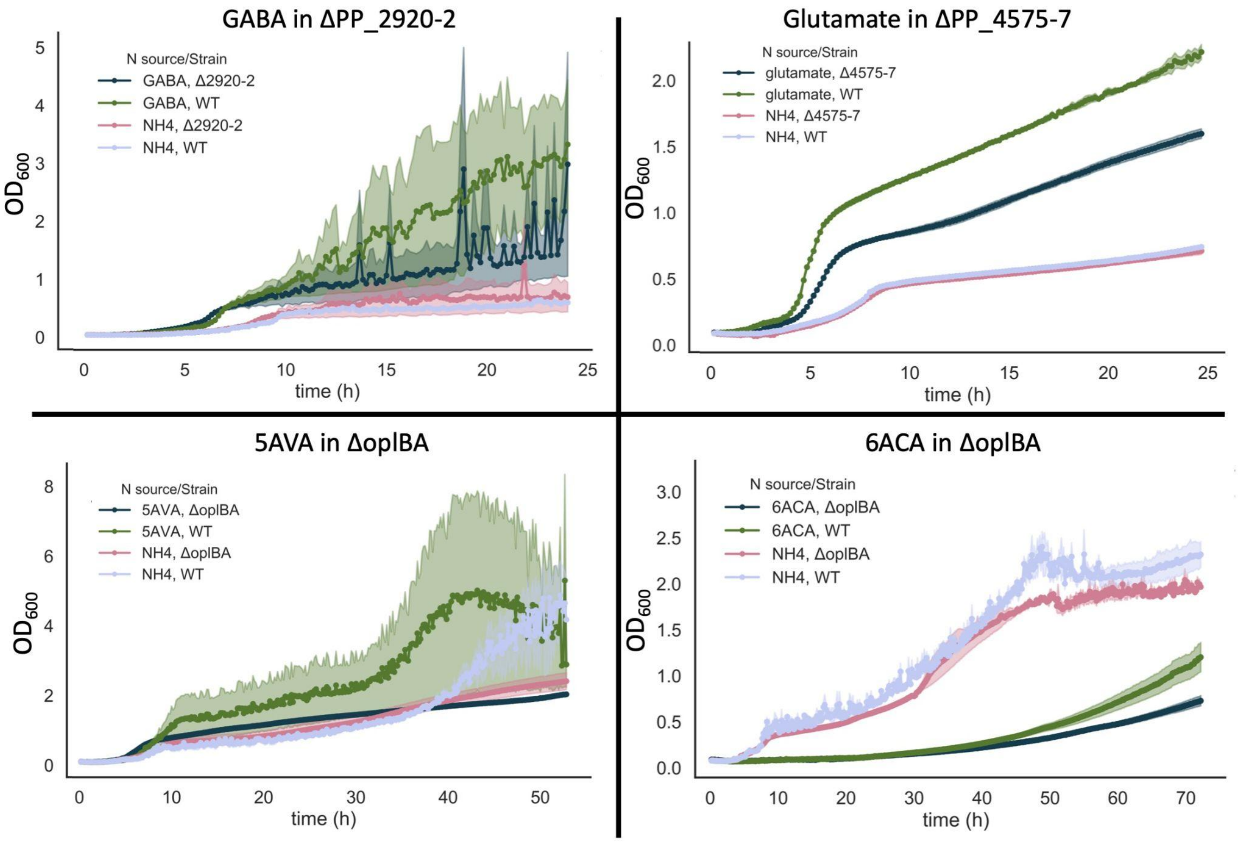


Figure S8: Growth curves of lactamase knockout strains with the ω-amino acid corresponding to the lactam substrate of the deleted lactamase as a nitrogen source. Growth with ammonium chloride as a nitrogen source appears the same in the wild-type and knockout strains, while growth with the ω-amino acid appears slightly hindered in the lactamase knockout strains.

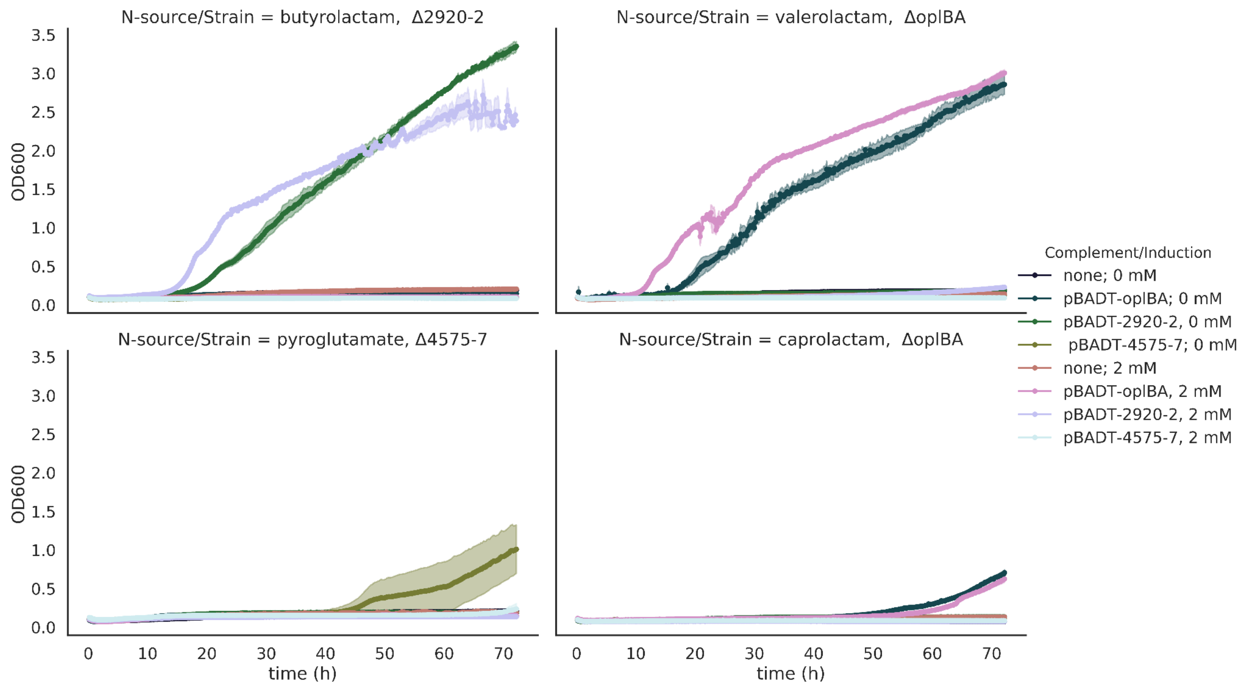


Figure S9: Complementation test of lactamase knockouts with plasmid-based expression of three different lactamases. For each lactam substrate, deletion strains of the native lactamase could only have growth restored with plasmid-based expression of the lactamase that was deleted, indicating high specificity of the lactamases.

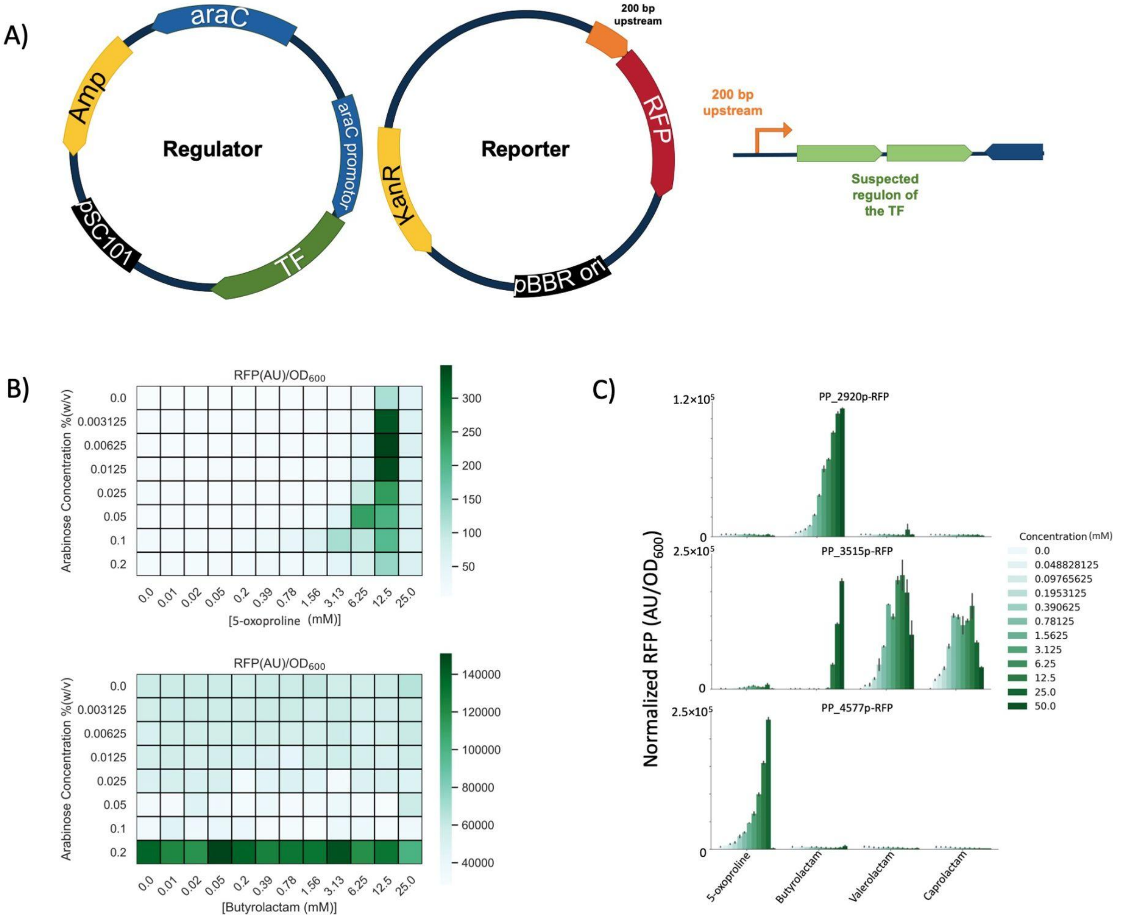


Figure S10: A) Genetic layout of two-plasmid biosensor systems. B) Initial two-plasmid test in *E. coli* XL1 blue, with varying expression of transcription factor and concentration of inducer. C) Reporter plasmid assay in *P. putida.* Two-fold dilutions of valerolactam, caprolactam, 5-oxoproline, and butyrolactam were tested at a concentration range from 0 to 50 mM (n = 3). Lactams were added to MOPS minimal medium with 20 mM glucose and 10 mM ammonium.

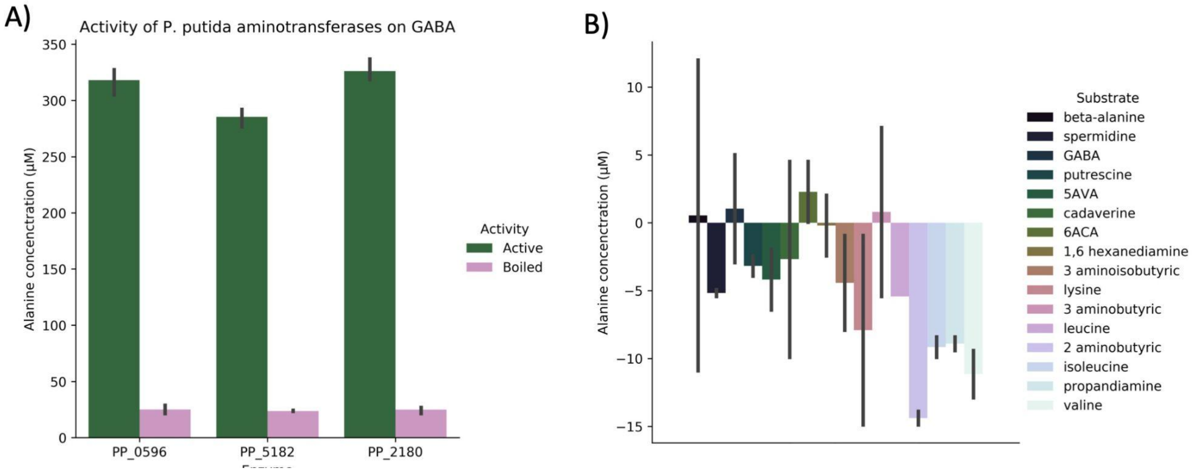


Figure S11: Biochemistry controls. A) Boiled enzyme control. Enzymes were boiled at 100C for 10 minutes prior to addition of substrate (n=3). B) No enzyme control to test if alanine assay responded to untransformed substrates (n=3)

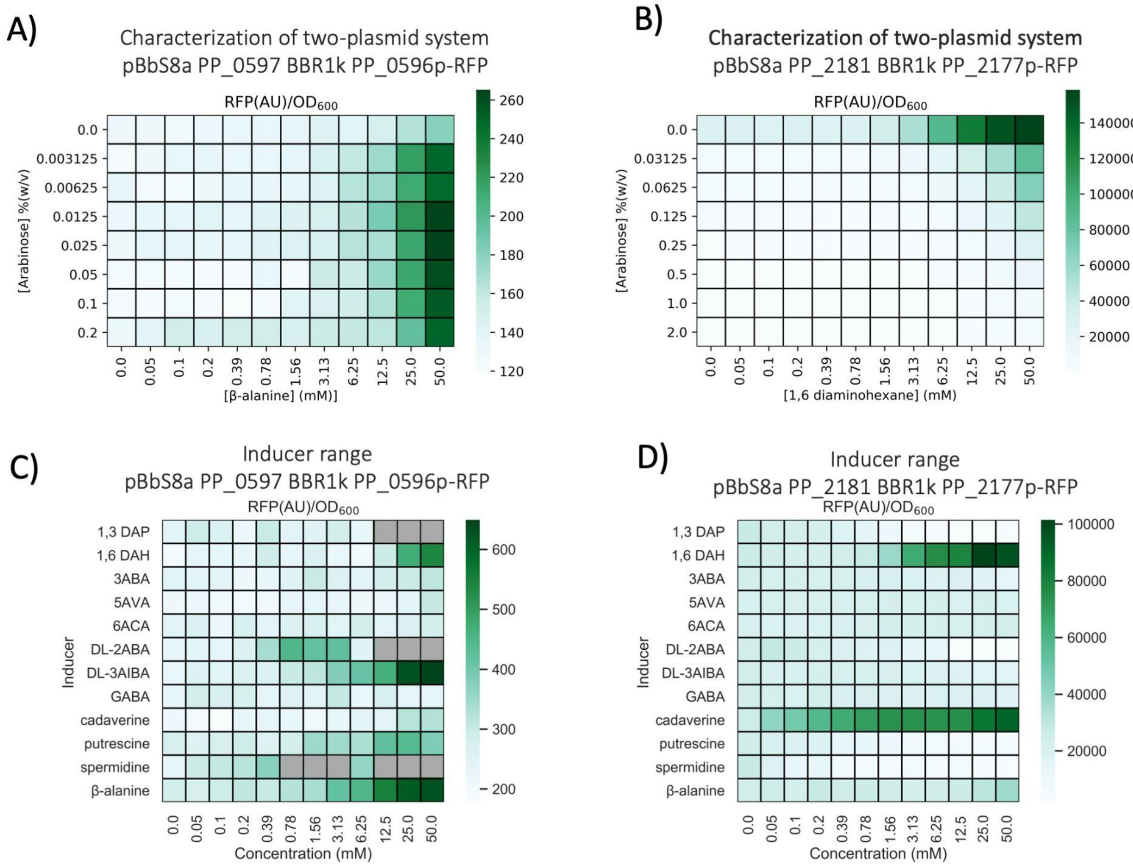


Figure S12: A) Two-plasmid biosensor system with the transcription factor PP_0597 and the promoter of PP_0596 B) Two-plasmid biosensor system with the transcription factor PP_2181 and the promoter of PP_2177 C) Inducer range of the pBbS8a PP_0597 BBR1k PP_0596p RFP system. Expression of PP_0596 was induced with 0.0125 wt% arabinose. Since RFP signal is normalized by OD, low OD can artificially inflate the apparent response. Gray squares indicate at the end of the 24 hour assay, final OD was less than double the starting OD. It appears that the presence of this biosensor system increases the toxicity of some of the inducers tested. This could be due to cross talk with native *E. coli* regulation. D) Inducer range of the pBbS8a PP_2181 BBR1k PP_2177p RFP system. Expression of PP_2181 relied on promoter leakiness, as was indicated optimal in part B.

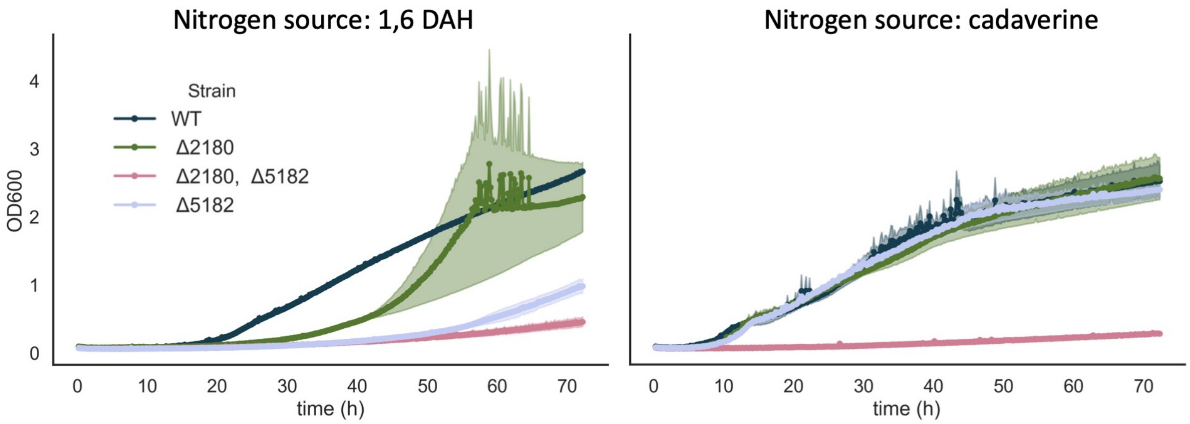


Figure S13: Growth of PP_2180 and PP_5182 deletion strains on the diamines cadaverine and 1,6-DAH. Shown are the WT (blue), ΔPP_2180 (green), ΔPP_2180 + ΔPP_5182 (pink), and ΔPP_5182 (grey) strains.

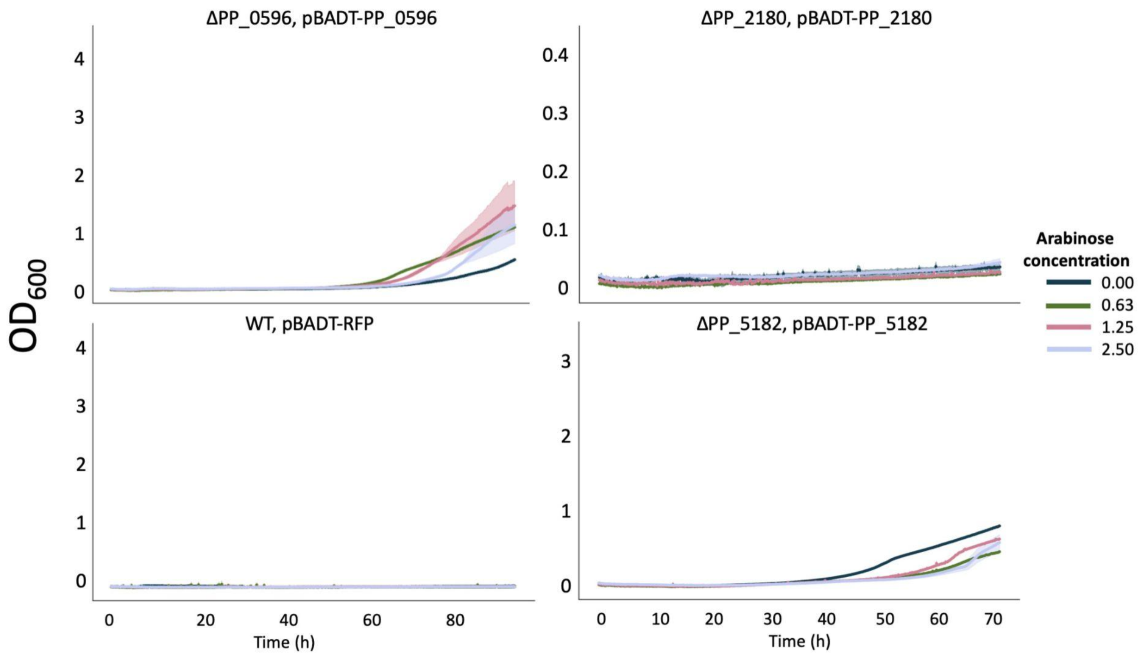

Figure S14: Complemented aminotransferase knockout strains grown with 6ACA as a sole nitrogen source.

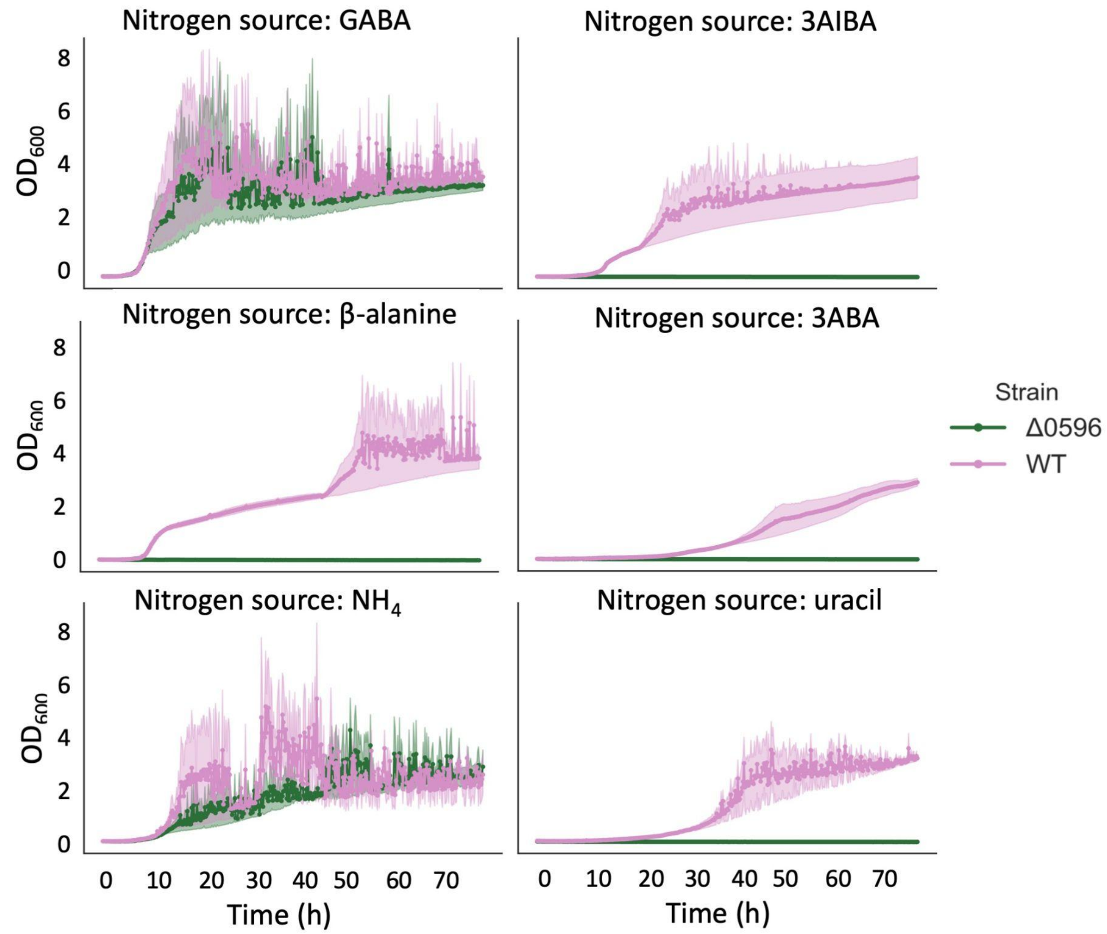


Figure S15: Comparison of growth of ΔPP_0596 (pink) vs WT (green) on various nitrogen sources. ΔPP_0596 appeared necessary for utilization of uracil, 3ABA, 3AIBA, and β-alanine.

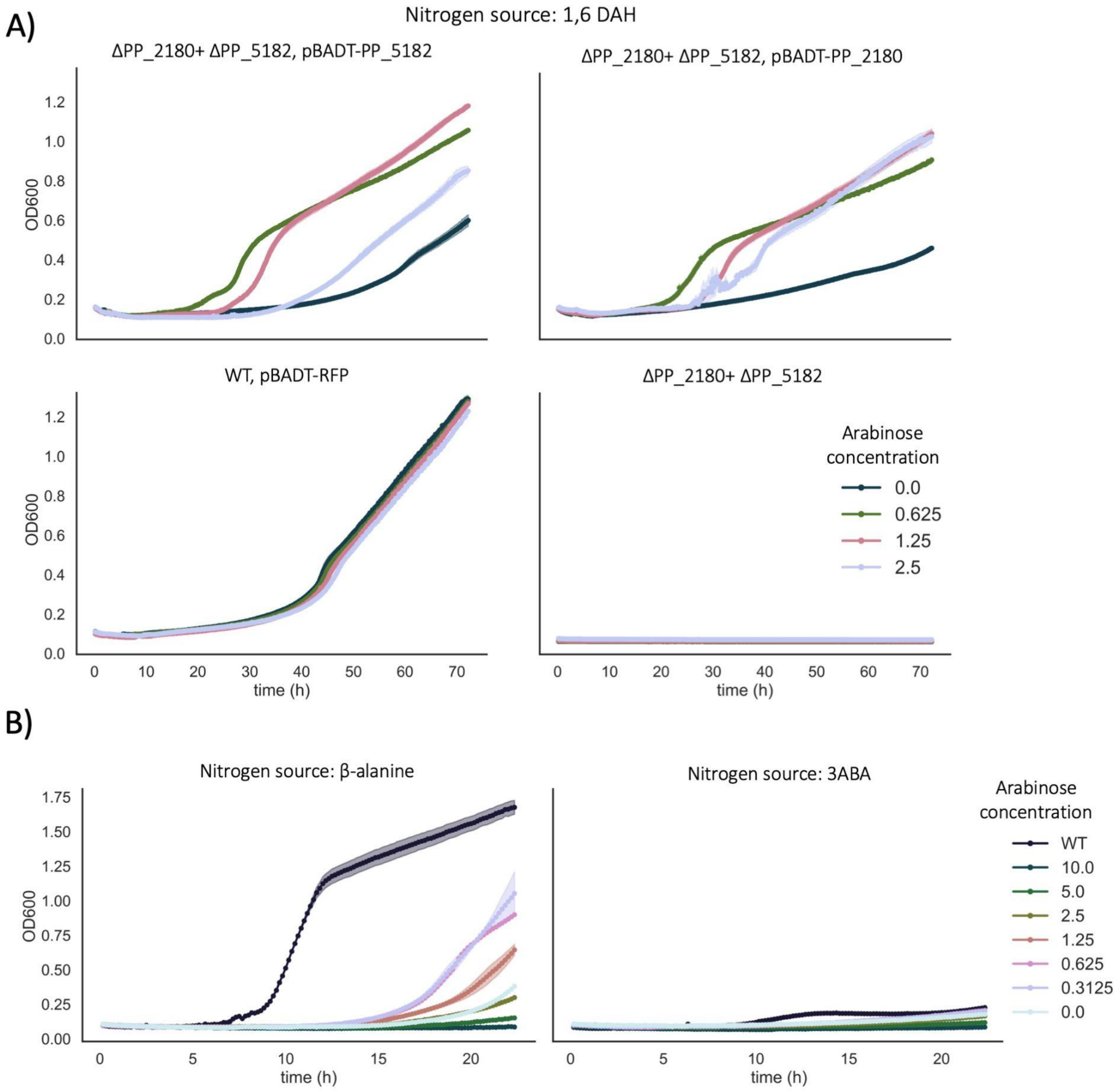


Figure S16: Complementation assays of aminotransferase knockouts. A) Complementation assays of ΔPP_2180 + ΔPP_5182 assay with either pBADT-PP_2180 or pBADT-PP_5182. Both singly expressed aminotransferases restore wild-type growth on 1,6-DAH. B) Complementation assays of ΔPP_0596 with pBADT-PP_0596. Complementation restores growth but not to wild type levels.
