## Additional Files for "Nitrogen metabolism in *Pseudomonas putida*: functional analysis using random barcode transposon sequencing": tsne.ipynb

```
# %% [code]
import math
import numpy as np
import pandas as pd
import time
import sklearn
from sklearn.cluster import AgglomerativeClustering
from sklearn.manifold import TSNE
import collections
from collections import OrderedDict
import itertools
import subprocess
import sys
from colour import Color
import matplotlib.pyplot as plt
import seaborn as sns
import altair as alt


# %% [code]

df_all=pd.read_csv('df_615.csv')
df_nits=df_all
df_nits.index=df_nits.locusId
df_nits.columns = df_nits.columns.str.split('.').str[0]
df_nits=df_nits.drop(columns='desc').iloc[:,2:-1]
df_nits
npdf = df_nits.to_numpy()
time2 = time.time()
e_ex=1
perplexity=5
n_iter=10000
lr=100
angle=0.1
models = TSNE(n_components=2, 
                      perplexity=perplexity, 
                      early_exaggeration=e_ex, 
                      learning_rate=lr, 
                      n_iter=n_iter, 
                      n_iter_without_progress=5000, 
                      min_grad_norm=1e-08, 
                      metric='euclidean', 
                      init='pca', 
                      verbose=1, 
                      random_state=None, 
                      method='exact', 
                      angle=angle, 
                      n_jobs=8)
tsne4=models.fit(npdf)
it=tsne4.n_iter_
tsne1=tsne4.embedding_
dftsnex=pd.DataFrame(tsne1)
dftsnex=dftsnex.rename(columns={0:'x',1:'y','0':'x','1':'y'})
df_nits['x']=dftsnex['x'].values
df_nits['y']=dftsnex['y'].values
df_nits['desc']=df_all['desc'].values
df_nits['sigcond']=df_all['sigcond'].values
df_nits.to_csv('tsneout.csv')
# %run tsne2.py
# %run tsne3.py

# %% [code]
dftsne=pd.read_csv('tsneout.csv')
dftsne.columns=dftsne.columns.str.split('.').str[0]
dftsne.index=dftsne['locusId']
dftsne2=dftsne.iloc[:,-4:-2]
dftsnex=dftsne2
ward_fit = AgglomerativeClustering(n_clusters=100, linkage='complete',distance_threshold=None,compute_full_tree=True).fit(dftsnex)
len(set(ward_fit.labels_))
dftsne['cluster'] = ward_fit.labels_


dic=dftsne.groupby(['cluster'])['sigcond'].agg(
    lambda x: pd.Series.mode(x)[0])
dic=dic.to_frame()
dicx=dftsne.groupby(['cluster'])['x'].agg(
    lambda x: pd.Series.mean(x))
dicx=dicx.to_frame()
dicx.head()
dicy=dftsne.groupby(['cluster'])['y'].agg(
    lambda x: pd.Series.mean(x))
dicy=dicy.to_frame()
dicy.head()
dicloc=dftsne.groupby(['cluster'])['locusId'].agg(
    lambda x: pd.Series.to_list(x))

dicloc=dicloc.to_frame()

dicloc['cluster'] = dicloc.index.values
dicloc['URL2'] = ['&locusId='.join(map(str, l)) for l in dicloc['locusId']]
dicloc['URL2'] = 'https://fit.genomics.lbl.gov/cgi-bin/genesFit.cgi?orgId=Putida&showAll=1&around=0&locusId=' + dicloc['URL2'].astype(str)
dftsny = dicy.join(dicx)
dftsny = dftsny.join(dicloc)
dftsny['clusternum'] = dftsny.cluster
dftsny = dftsny.replace({'cluster':dict(zip(dic.index, dic['sigcond']))})

dftsne['clusternum']=dftsne.cluster
dftsne = dftsne.replace({'cluster':dict(zip(dic.index, dic['sigcond']))})
dftsne = dftsne.rename(columns={0:'x',1:'y'})

dftsne.index=dftsne['clusternum']

dftsne = dftsne.sort_index()
from sklearn.decomposition import PCA
pca = PCA(n_components=60)
principalComponents = pca.fit_transform(dftsne.iloc[:,1:-4])
principalDf = pd.DataFrame(data = principalComponents)
principalDf.index = dftsne.locusId

ppdic=pd.read_csv('ppgene_dictionary.csv')

dftsne['gene'] = dftsne.locusId.values
dftsne['status'] = dftsne.locusId.values
dftsne['uniprot'] = dftsne.locusId.values
dftsne=dftsne.replace({'uniprot':dict(zip(ppdic.PP,ppdic.Entry))})
dftsne = dftsne.replace({'gene':dict(zip(ppdic.PP, ppdic.gene))})
dftsne = dftsne.replace({'status':dict(zip(ppdic.PP, ppdic.Status))})

dftsne['PCAnul']=dftsne.locusId
dftsne = dftsne.replace({'PCAnul':dict(zip(principalDf.index,principalDf[0]))})
dftsne['PCAone']=dftsne.locusId
dftsne = dftsne.replace({'PCAone':dict(zip(principalDf.index,principalDf[1]))})
dftsne.index=dftsne.locusId

dicond=dftsne.groupby(['clusternum'])['sigcond'].agg(
    lambda x: pd.Series.to_list(x))
dicond=dicond.to_frame()
dftsny=dftsny.join(dicond)
dicgen=dftsne.groupby(['clusternum'])['gene'].agg(
    lambda x: pd.Series.to_list(x))
dicgen=dicgen.to_frame()
dftsny=dftsny.join(dicgen)
dftsny=dftsny.rename(columns={'cluster':'class'})
dftsne=dftsne.rename(columns={'cluster':'class'})
dftsne.iloc[:,1:].to_csv('dftsnew.csv')
dftsny.to_csv('dftsnyw.csv')

# %% [code]
dftsne=pd.read_csv('dftsnew.csv')
dftsny=pd.read_csv('dftsnyw.csv')
classlen = len(set(dftsne["sigcond"].unique()))
classlen

grey = Color('#c2c0c0')
red = Color('#ff001e')
green = Color('#006918')
colg=math.trunc(classlen/3)
colb=classlen-2*colg


colors = list(grey.range_to(Color('#000000'),colb))
colorsc =list(green.range_to(Color('#ffd900'),colg)) #toggle these based on classlen
colorsb= list(red.range_to(Color("#0d00ff"),colg))
             
colors = [str(i) for i in colors]
colorsb= [str(i) for i in colorsb]
colorsc=[str(i) for i in colorsc]
             
stroke_palette = colorsb +colorsc+ colors 

chart1=0
chart2=0
lst1=0
lst2=0


selection1 = alt.selection_multi(fields=['class'], bind='legend')
selection2 = alt.selection_multi(fields=['sigcond'],bind='legend')
lst1 = list(dftsne.locusId.values)
lst2 = list(dftsne.locusId.values)
#str2 = str1.insert(0,None)
lst2.sort()
lst1.sort()
lst2.insert(0,None)
lst1.insert(0,'Show All')
lst2.insert(0,'PP_XXXX')
lst1.insert(0,'Hide All')


input_dropdown = alt.binding_select(options=lst2,labels=lst1)
gene = alt.selection_single(name='gene',fields=['locusId'], bind=input_dropdown)
click = alt.selection_multi(on='click', fields=['locusId'])
click2 = alt.selection_multi(on='click', fields=['clusternum'])
brush = alt.selection_interval(
    on="[mousedown[event.altKey], mouseup] > mousemove")
ranked_text = alt.Chart(dftsne).mark_text().encode(
#    x=10,
    y=alt.Y('row_number:O',axis=None)
).transform_window(
    row_number='row_number()'
).transform_filter(
    selection2|gene|brush|click|selection1|click2
).transform_window(
    rank='rank(row_number)'
).transform_filter(
    alt.datum.rank<60
)
genelst = ranked_text.encode(text='locusId').properties(title='LocusId')
loclst = ranked_text.encode(text='gene').properties(title='Gene')
stat = ranked_text.encode(text='status').properties(title='Uniprot Status')
desc = ranked_text.encode(text='desc').properties(title='Description')
entry = ranked_text.encode(text='uniprot').properties(title='Uniprot Entry')
cond2 = ranked_text.encode(text='sigcond').properties(title='Max |fit|')
cond1c = ranked_text.encode(text='x').properties(title='x')
cond2c = ranked_text.encode(text='y').properties(title='y')

dftsne=dftsne.sort_values('clusternum')
chart2 = alt.Chart(dftsne,title='t-SNE clustering of published BarSeq data (w/ PCA initialization)').mark_point(size=90,filled=True).encode(
     x=alt.X('x',axis=alt.Axis(labels=True),title=None#scale=alt.Scale(domain=(-100, 120))
            ),
     y=alt.Y('y',axis=alt.Axis(labels=True),title=None#scale=alt.Scale(domain=(-95, 80))
            ),
     tooltip=['locusId','gene','sigcond','clusternum'],
     color=alt.Color('sigcond', legend=alt.Legend(columns=2,symbolLimit=0,orient='left'), 
         scale=alt.Scale(range=stroke_palette)),
     #href='URL',
     shape=alt.Shape('sigcond', 
            scale=alt.Scale(range=["triangle-up","circle","triangle-down","square",
                        "triangle-right","cross","triangle-left","diamond"]),
            legend=alt.Legend(columns=1,symbolLimit=0,title='')),
     size=alt.condition(selection2|click|gene|brush|click2|selection1, alt.value(200), alt.value(90)), 
     opacity=alt.condition(selection2|click|gene|brush|click2|selection1, alt.value(1.), alt.value(0.05))                                                             
     ).properties(width=600,height=600).add_selection(selection2,click,gene,brush,click2).interactive()

f=str(len(set(dftsne.clusternum)))

chart1 = alt.Chart(dftsne, title='t-SNE clustering of published BarSeq data (w/ PCA initialization), clusters='+f).mark_point(size=90,filled=True).encode(
     x=alt.X('x',axis=alt.Axis(labels=True),title=None#scale=alt.Scale(domain=(-100, 120))
            ),
     y=alt.Y('y',axis=alt.Axis(labels=True),title=None#scale=alt.Scale(domain=(-95, 80))
            ),
     tooltip=['locusId','gene','class','clusternum','sigcond'],
     color=alt.Color('class', legend=alt.Legend(columns=2,symbolLimit=0,orient='left',title=''), 
         scale=alt.Scale(range=stroke_palette)),
     #href='URL',
     shape=alt.Shape('class', 
            scale=alt.Scale(range=["triangle-up","circle","triangle-down","square",
                        "triangle-right","cross","triangle-left","diamond"]),
                    ),#legend=None),
            #legend=alt.Legend(columns=1,symbolLimit=0)),
     size=alt.condition(gene|brush|click2|click|selection1, alt.value(200), alt.value(90)), 
     opacity=alt.condition(gene|brush|click2|click|selection1, alt.value(1.), alt.value(0.05))                                                             
     ).properties(width=600,height=600).add_selection(gene,brush,click2,selection1)#.interactive()
chartz = alt.Chart(dftsny, title='Cluster centroids with links to fitness browser').mark_point(size=90,filled=True).encode(
     x=alt.X('x',axis=alt.Axis(labels=True),title=None#scale=alt.Scale(domain=(-100, 120))
            ),
     y=alt.Y('y',axis=alt.Axis(labels=True),title=None
            ),
     tooltip=['locusId','class','clusternum'],
     color=alt.Color('class', legend=None,title='Condition', 
         scale=alt.Scale(range=stroke_palette)),
     href='URL2',
     shape=alt.Shape('class', 
            scale=alt.Scale(range=["triangle-up","circle","triangle-down","square",
                        "triangle-right","cross","triangle-left","diamond"])),
            #legend=alt.Legend(columns=1,symbolLimit=0)),
     size=alt.condition(gene|brush|click2, alt.value(200), alt.value(90)), 
     opacity=alt.condition(gene|brush|click2, alt.value(1.), alt.value(0.05))                                                             
     ).properties(width=600,height=600).add_selection(click2,gene,brush)#.interactive()


chart5=alt.hconcat(chart1,chart2,)
chart6=alt.hconcat(chartz,loclst,genelst,cond2,stat,entry,desc)
chart4 = alt.vconcat(chart1,chart6)
chart9 = alt.hconcat(chart5,chart4)
chartx = alt.vconcat(chart5,chart6)

chart8=alt.vconcat(chart2,chart6)


chartx.configure_view(strokeOpacity=0)#.save('Schmidt_Pearson_t-SNE-Nit-BarSeq.html')

# %% [code]


# %% [code]


# %% [code]
```
